# A metabolic labelling-based lipid imaging technology establishes VPS13A as a phosphatidylethanolamine lipid transfer protein

**DOI:** 10.64898/2026.06.09.731182

**Authors:** Saswati Biswal, Manas, Tanveer Alam Khan, Shreya, Gandhi Bhukya, Shivangi Singh, Bhupali Thakuria, Avinil Das Sharma, Nachiket Dhonnar, Jeet Kalia

**Affiliations:** Department of Biological Sciences, Indian Institute of Science Education and Research (IISER) Bhopal Bhopal Bypass Road, Bhauri, Bhopal–462066, Madhya Pradesh, India; Department of Chemistry, Indian Institute of Science Education and Research (IISER) Bhopal Bhopal Bypass Road, Bhauri, Bhopal–462066, Madhya Pradesh, India

## Abstract

The ability to image specific lipid subtypes within cells can have a transformative impact on the study of lipid dynamics and trafficking mechanisms. Herein, we describe a technology for imaging phosphatidylethanolamine (PE) lipids in live mammalian cells that involved screening a library of ethanolamine derivatives to identify an azido compound that efficiently metabolically labels PE. Crucially, this probe evades the cellular methylation machinery specifically labelling PE without forming labelled methylated PE and phosphatidylcholine (PC) lipids. The administration of cyclooctyne dyes to cells metabolically labelled with this probe rendered azido PE lipids fluorescent via strain-promoted click chemistry, enabling imaging. We employed this technology to image PE in various cellular organelles, visualize PE externalization during apoptosis, and discover that the VPS13A protein transports PE from the endoplasmic reticulum to the mitochondria. This technology will facilitate addressing fundamental questions in PE biology and studying dysregulation of PE dynamics and trafficking in disease states.

## Introduction

Phosphatidylethanolamine (PE) lipids account for ∼15–25% of the mammalian cellular lipidome^1^. They contribute towards inducing negative curvature on the plasma membrane to orchestrate cytokinesis and the maintenance of extensive invaginations in the inner mitochondrial membrane maximizing membrane surface area for optimal ATP production^2^. Additionally, PE functions as a lipid chaperone for folding membrane proteins and is intimately associated with important cellular processes such as apoptosis, ferroptosis and autophagy^2^. Considering its diverse important roles in cellular physiology, it is not surprising that the dysregulation of cellular PE levels is associated with serious diseases including spastic paraplegia, Liberfarb syndrome, and Parkinson’s disease^3^. Moreover, alterations in the phosphatidylcholine (PC)/PE ratio in the endoplasmic reticulum (ER) membrane triggers ER stress associated with obesity, fatty liver disease, and diabetes^4,5^.

To fully understand the roles of PE lipids in cellular physiology and disease pathophysiology, molecular-level insights into their cellular dynamics (for example, inter-organellar exchange) are required. Although PE biosynthesis has been extensively studied^2,6,7^, the cellular dynamics of PE remains poorly understood primarily due to the lack of technologies for imaging PE within live cells. Fluorescently labelled cyclic peptides such as duramycin and cinnamycin lantibiotics that specifically bind to the headgroup of PE lipids have been used to image PE externalized to the outer cell surface during apoptosis and cytokinesis^8–10^. This approach is ideal for imaging cell surface PE but is not compatible with imaging PE within organellar membranes of live cells owing to the lack of membrane permeability of these cyclic peptide probes. Clearly, a robust technology for imaging PE lipids in live cells is urgently required to study PE cellular dynamics.

Inspired by previous success in metabolic labelling-based approaches for lipid labelling^11–27^, we envisaged a PE imaging technology involving the cellular administration of their alkynyl/azido metabolic precursors to install these chemical handles into PE lipids rendering them amenable to fluorescent labelling via azide–alkyne click chemistry (**Figure 1A**). Most PE lipids within mammalian cells are biosynthesized via the Kennedy pathway that entails the uptake of ethanolamine followed by its sequential conversion into phosphoethanolamine, cytidine diphospho (CDP) ethanolamine and finally into PE lipids^7^ (**Figure S1**, Supporting Information section). To implement our metabolic labelling strategy, we planned on administering alkynyl/azido ethanolamine analogs to mammalian cells in culture with the goal of forming PE lipids labelled on their headgroups with these bioorthogonal chemical handles.

**Figure 1.**
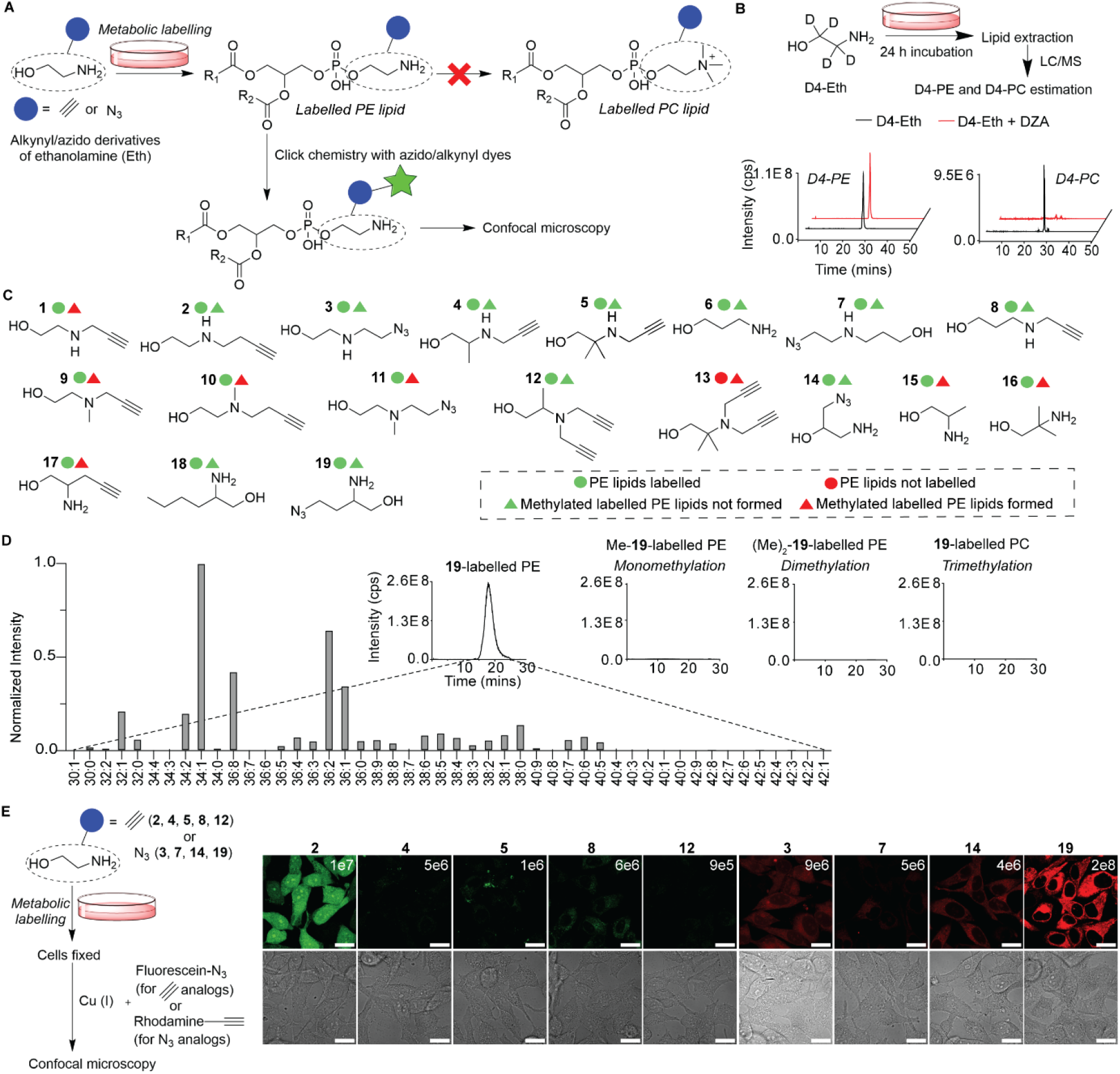
Identification of suitable probes for PE imaging. A) A schematic representation of our metabolic labeling strategy for PE imaging involving the administration of ethanolamine analogs labelled with biorthogonal alkynyl/azido groups to mammalian cells followed by tagging with fluorophores via click chemistry. An ideal probe should efficiently label PE lipids, and not form methylated derivatives of labelled PE, including PC. B) Top: Workflow for evaluating the propensity of mammalian cells to form D4-PE and D4-PC lipids upon administration of D4-ethanolamine (Eth). Bottom: Results of LC/MS lipidomics experiments performed on lipids isolated from HeLa cells subjected to the above workflow demonstrating biosynthesis of D4-PE lipids and also D4-PC lipids (black traces in the LC/MS profiles). Co-administering DZA, a generic inhibitor of methyltransferase activity, potently inhibits D4-PC formation, and has negligible effect on D4-PE formation (red traces in the LC/MS profiles). Bar graphs representing quantitative comparisons over three replicates are provided in Supporting Information section **Figure S5**. C) Summary of our lipidomics-based screening efforts involving the cellular administration of a library of 19 ethanolamine analogs to HeLa and HEK293 cells. The circular symbols represent the results of the PE labelling experiments with green circles annotating successful PE labelling and red denoting failure to metabolically label PE lipids. Triangular symbols denote the methylation results (green: methylated versions of PE lipids labelled with analogs not detected; red: methylated versions of PE lipids labelled with analogs detected). D) Lipidomics data for compound **19**, our most efficient PE labelling probe depicting proficient PE labelling and no detectable formation of monomethylated/dimethylated/trimethylated versions of **19**-labelled PE. E) Evaluation of the ability of PE labelling probes that do not undergo methylation identified above as PE imaging probes. Left panel: Our fixed cell-based imaging workflow. Right: Imaging results (top panel: fluorescence, bottom panel: bright field) obtained with HeLa cells metabolically labelled with the probes (2 mM) with the lipidomic peak intensities of probe-labelled PE lipids formed depicted on the top-right of each of the fluorescent imaging data (averaged over three replicates). Scale bars = 20 μm.

Although previous experiments on flies have demonstrated that the Kennedy pathway is capable of accepting ethanolamine analogs^28^, a major caveat of this approach is the possibility that it would yield not just labelled PE lipids but also labelled PC and/or methylated versions of PE lipids due to the well-established propensity of PE to undergo sequential methylation within mammalian cells^29–31^. Subsequent treatment with clickable dyes would, therefore, yield not just fluorescent PE lipids but also its methylated derivatives within cells precluding the specific imaging of PE lipids. Consequently, a key goal of ours was to develop clickable ethanolamine analogs that would be efficiently processed by the Kennedy pathway to form labelled PE lipids, and yet, not form methylated PE lipids within cells (**Figure 1A**).

We envisioned that our PE imaging technology would provide a valuable toolkit to unravel the poorly understood ER–mitochondria lipid crosstalk mechanisms, which hold particular significance for PE lipids. The mitochondrial PE pool has contributions from PE synthesized in the ER via the Kennedy pathway as well as that synthesized by the mitochondrial PS decarboxylase (PISD) enzyme that converts PS to PE^32,33^. These two major PE biosynthesis pathways in mammalian cells generate different PE species, with the Kennedy pathway primarily forming mono- and di-unsaturated fatty acid-containing PE lipids, and the PS decarboxylation preferably synthesizing polyunsaturated PE lipids^34^. Not surprisingly, therefore, these two pathways are mutually non-compensatory and deleting either *pisd* or the Kennedy pathway gene, *pcyt2*, causes embryonic lethality in mice^2^. Dysfunction of either pathway is pathological—the disruption of the Kennedy pathway is associated with hereditary spastic paraplegia and childhood-onset neurodegeneration (CONATOC) whereas mutations in *pisd* are linked to the mitochondrial disease, Liberfarb syndrome^3^. Clearly, an in-depth understanding of the crosstalk between the Kennedy and PS decarboxylation pathways will contribute tremendously towards unraveling the pathophysiology of these diseases and understanding their roles in PE biology. In this context, a key unanswered question is how PE synthesized in the ER is transported to the mitochondria especially considering that mitochondria do not partake in cellular vesicular trafficking routes and rely on lipid trafficking proteins for inter-organellar lipid exchange^35^. Recently, novel PC imaging technologies have been employed to identify proteins involved in PC lipid trafficking from the ER to the mitochondria^19,20^. No such studies have been reported on PE lipids due to the lack of a robust PE imaging platform capable of imaging lipids within live cells. Herein, we fill this knowledge gap by employing our PE imaging technology to discover that the VPS13A protein expressed at ER–mitochondria contact sites traffics PE lipids from the ER to the mitochondria.

## Results and Discussion

### HEK293 and HeLa cells form PC lipids when administered with ethanolamine

We began our studies by characterizing the propensity of two commonly used mammalian cell lines, HeLa and HEK293, to form methylated PE lipids. LC/MS analysis on lipids isolated from cultures of both of these cell types incubated with D4-ethanolamine for 24 h revealed the formation of D4-PE lipids as well as D4-PC lipids (HeLa: LC/MS traces in black in **Figure 1B**, HEK293: **Figure S4**, and **Figure S5** summarizing experiments performed in three replicates). Including the *S*-adenosylmethionine (SAM) cycle inhibitor, DZA, which has been previously demonstrated to inhibit PE to PC conversion in mammalian cells^36^ dramatically reduced the levels of D4-PC without altering D4-PE formation (HeLa: traces in red in **Figure 1B** and **Figure S5**, HEK293: **Figures S4 and S5**). These results establish that both HeLa and HEK293 cell lines are adept at taking up ethanolamine and biosynthesizing methylated PE species. Therefore, the success of our metabolic labelling-based approach for specifically imaging PE was contingent on the development of alkynyl and/or azido ethanolamine analogs that would label PE efficiently, and yet, not form methylated versions of labelled PE lipids.

### Lipidomics screening of ethanolamine analogs to discover specific PE labelling probes

In the quest for an ideal ethanolamine probe for PE-specific labelling, we were encouraged by a previous report that suggested that the mammalian PE methyl transferase enzyme responsible for methylation of PE in mammalian cells demonstrates specificity for turning over PE derivatives possessing certain ethanolamine analogs over others^30,31^. Seeking to build on these studies, we subjected a library of ethanolamine analogs possessing a range of chemical architectures (synthesis: **Schemes S1–S11** in the Supporting Information section) to metabolic labelling followed by LC/MS-based lipidomics analysis on HeLa and HEK293 mammalian cells in culture (details of MS scan modes and other specifics pertaining to lipidomics are described in the Supporting Information section).

Our lipidomics experiments resulted in the identification of several compounds that specifically labelled PE in HEK293 and HeLa cells (annotated with green circular as well as green triangular symbols in **Figure 1C**) with both cell lines yielding identical trends for the probes (**Figure 1C** and **Figures S6, S7**). Examples of such molecules include *N*-alkyl ethanolamines containing the *N*-butynyl substituent (compound **2, Figure 1C**), the *N*-azidoethyl group (**3**), and compounds containing both the *N*-propargyl as well as *C*-alkyl substitutions (**4** and **5**).

Propanolamine derivatives also resisted the formation of *N*-methylated lipid derivatives (**6**–**8**). In contrast, *N,N*-dialkyl ethanolamine compounds were not as adept at evading methylation with the alkynes **9** and **10**, and the azide **11** forming labelled PC lipids in addition to PE. The structurally more complex analog, **12**, containing two *N*-propargyl substituents and a methyl group on the carbon adjacent to the nitrogen of the ethanolamine labelled PE specifically without undergoing methylation. Introducing another methyl group at the carbon atom yielded probe **13** which did not label PE and also did not yield methylated versions of labelled PE.

We also tested a range of *C*-alkyl ethanolamines including **14** wherein an azidomethylene group is appended to the carbon adjacent to the hydroxyl group of ethanolamine, and **15**–**19** that are substituted at the carbon atom adjacent to amino group of ethanolamine. Our lipidomics experiments revealed that **14** exclusively labelled PE, and among **15**–**19**, only those analogs that contained a long alkyl substitution (**18** and **19**) evaded methylation. In addition to forming the desired labelled PE lipids, **15** formed dimethylated derivatives of labelled PE and also underwent trimethylation to form labelled PC lipids, whereas **16** yielded monomethylated labelled PE, and **17** formed both mono and dimethylated labelled PE derivatives (**Figures S6– S9** contain all the LC/MS profiles and lipidview analysis depicting individual labelled lipid species detected with all probes in HeLa and HEK293 cells). These results demonstrate that formation of methylated PE lipids is extremely sensitive to ethanolamine structures.

Among all compounds tested, **19** labelled PE most efficiently, yielding an LC/MS intensity of ∼2e8 for labelled PE lipids enabling us to detect 48 PE lipid species labelled with **19** (**Figure 1D**). MS scans aimed at measuring methylated PE, dimethylated PE and trimethylated PE (labelled PC) yielded no detectable signals for these species demonstrating that **19** labels PE exclusively.

### Identification of an ideal probe for PE imaging

With several alkynyl and azido ethanolamine analogs that specifically label PE in hand, we proceeded to evaluate them for imaging PE lipids in HeLa cells. Our initial survey involved administering the compounds at a concentration of 2 mM, allowing metabolic labelling to proceed for 24 h, fixing the cells and subjecting them to click chemistry conditions with either azido fluorescein (in case of alkynyl probes **2, 4, 5, 8**, and **12**) or alkynyl rhodamine (in case of azido probes **3, 7, 14**, and **19**). These experiments yielded the highest fluorescent signals for **2** among the alkynyl compounds tested, and for **19** among the azides (**Figure 1E**). Importantly, these results followed the same trend as that obtained in the lipidomics experiments as compounds **2** and **19** yielded the highest lipidomics intensity signal among the alkynyl and azido probes respectively.

Whereas click chemistry with terminal alkynes entails the use of cytotoxic Cu (I) rendering them not ideal for live cell applications, azides undergo Cu-free strain-promoted click chemistry-mediated labelling with cyclooctyne dyes without requiring any other reagents in live cells. Consequently, we performed our subsequent imaging experiments with the azido probe, **19** by employing bicyclononyne (BCN)-BODIPY as a cyclooctyne dye (**Figure 2A**, left panel).

**Figure 2.**
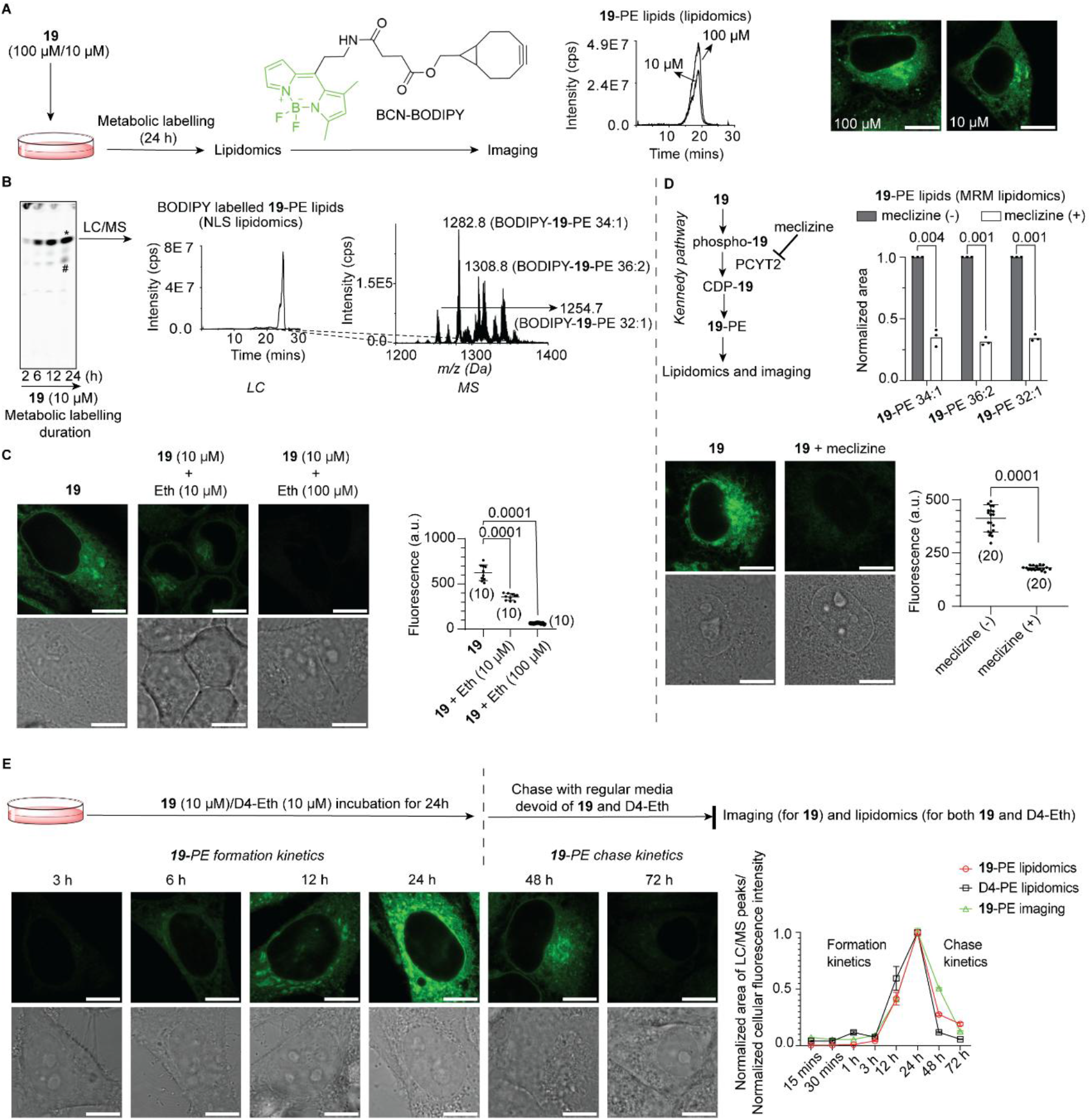
Live-cell imaging of PE lipids. A) Left: A schematic representation of our live HeLa cell imaging workflow employing probe **19** at concentrations within the range of that of native ethanolamine. Lipidomics (middle panel) and imaging (right) results for **19**-PE formed in HeLa cells incubated with **19** (100/10 μM). B) TLC analysis on fluorescently labelled **19**-PE lipids isolated from metabolically labelled cells post *in cellulo* click chemistry with BCN-BODIPY. The prominent fluorescent spot denoted with an asterisk was scraped and subjected to LC/MS analysis (middle panel: LC trace, and MS analysis of the LC peak on the right) conforming the presence of BODIPY-**19**-PE lipids. The minor spot on the TLC denoted by a hash was characterized by LC/MS and annotated as BODIPY-**19**-carboxymethyl PE species formed as a result of condensation between **19**-PE and glucose (**Scheme S12** and **Figure S11**). C) Competition experiments with native ethanolamine depicting fluorescence images on the left panel and quantitative analysis under each condition on the right. The numbers within parenthesis denote the number of cells for which the fluorescence intensities were measured under each condition. D) Kennedy pathway inhibition experiments employing meclizine, an inhibitor of the PCYT2 enzyme. Multiple reaction monitoring (MRM)-based lipidomics (top-right) and imaging (bottom) experiments depicting strong inhibition of **19**-PE formation in the presence of meclizine (50 μM) over a 6 h incubation period. E) Time course-based experiments to compare **19**-PE formation and catabolism (chase) with that for D4-PE upon metabolic labelling with either **19** or D4-Eth (both administered at 10 μM). The workflow is depicted schematically on the top; the imaging data is depicted on the bottom-left panel; and quantitative analysis by imaging (for **19**-PE) and lipidomics (for both **19**-PE and D4-PE) is depicted on the plot on the bottom-right. Scale bars = 10 μm.

### Imaging PE in live HeLa cells using 19 as a probe

An ideal metabolic labelling probe must operate within a concentration range similar to that of the native metabolite. Since ethanolamine is typically present between ∼10 and 100 μM in the extracellular milieu^7,37^, we evaluated the amenability of imaging PE using **19** within this concentration range. Lipidomic characterization of PE labelling demonstrated **19**-PE lipid formation upon the cellular administration of both 100 μM as well as 10 μM concentrations of **19** (**Figure 2A**, middle panel). Imaging experiments yielded high intensity fluorescent signals even for cultures treated with 10 μM of **19** (**Figure 2A**, right panel), demonstrating that our technology can image PE lipids in mammalian cells by administering the probe at concentrations close to the lower end of the physiological native metabolite concentrations. Moreover, cellular cytotoxicity experiments on HeLa cells incubated with 10 μM–4 mM **19** revealed that **19** demonstrated negligible cytotoxicity in this wide range of concentrations (**Figure S10**), establishing its compatibility for use as a probe on mammalian cells. Taken together, these experiments establish the technical robustness of our technology.

Administering **19** (10 μM) to HeLa cells in culture for different time periods followed by treatment with BCN-BODIPY and TLC analysis on extracted lipids yielded a major fluorescent spot (denoted by an asterisk in **Figure 2B**) whose intensity increased in an incubation time-dependent fashion. Scraping this spot off the TLC followed by LC/MS (middle and right panels of **Figure 2B**) yielded masses for BCN-BODIPY conjugated **19**-PE lipids, establishing that our workflow does indeed result in the formation of fluorescently labelled PE lipids in HeLa cells.

To investigate the PE labelling specificity of our technology, we performed competition with ethanolamine by co-administering it with **19**. Administering 10 μM of both **19** and ethanolamine caused a ∼2-fold reduction in the fluorescence intensity of BCN-BODIPY labelled PE lipids as imaged by confocal microscopy whereas ten-fold excess ethanolamine profoundly ablated **19**-PE formation yielding barely detectable fluorescence signals (**Figure 2C**).

Another experiment that we performed to validate our technology focused on establishing that **19** was forming **19**-PE lipids via the Kennedy pathway as designed. This experiment involved administering HeLa cells with **19** and then inhibiting the Kennedy pathway by introducing meclizine, an inhibitor of the PCYT2 enzyme that catalyzes the rate determining step of the Kennedy pathway for PE biosynthesis—the conversion of phosphoethanolamine to CDP-ethanolamine (**Figure 2D**, top-left panel and **Figure S1**). Lipidomics demonstrated a ∼70% inhibition in the biosynthesis of three major labelled PE species in presence of meclizine (**Figure 2D**, top-right panel) and also resulted in a dramatic attenuation of fluorescence signals from **19**-administered HeLa cells treated with BCN-BODIPY as evident from both the representative image and fluorescence intensities averaged over 20 cells (**Figure 2D**, bottom panel). These competition and biosynthesis inhibition experiments established that **19** is an effective substrate for the Kennedy pathway.

To more rigorously compare **19** and ethanolamine with respect to PE biosynthesis and to compare the rates of catabolism of **19**-PE with native PE, we performed time course experiments wherein we incubated HeLa cells in culture with either **19** or D4-ethanolamine (both 10 μM) and subjected them to our imaging workflow (for **19**-administered samples), and lipidomics characterization (for both **19** and D4-ethanolamine administered samples). After a 24 h incubation with **19** or D4-ethanolamine, we performed chase experiments with regular media devoid of these metabolic labeling agents (**Figure 2E**, top panel). The results of the imaging experiments demonstrate a time-dependent enhancement in the fluorescence intensity over 24 h of incubation time with **19** which disappears over 48 h of chase (the period between the 24 h and 72 h time points in **Figure 2E**). A time course plot depicting normalized fluorescent intensities for BCN-BODIPY-**19**-PE and lipidomic quantification for **19**-PE formation and chase (**Figure 2E**, bottom-right panel) yielded overlapping traces establishing that the fluorescence intensities observed in our imaging experiments directly correlated with PE labelling by **19**. Moreover, these traces also overlapped with the lipidomics data for formation and chase for D4-ethanolamine labelled PE lipids (**Figure 2E**, bottom-right panel) demonstrating that **19** is an ideal surrogate for ethanolamine with respect to PE biosynthesis and catabolism.

### Employing 19 to image PE externalization during apoptosis

We next employed our technology to visualize the cellular dynamics of PE lipids by focusing on apoptosis—a physiological process that is accompanied by the redistribution of PE lipids between the inner and outer leaflets of the plasma membrane^38^. Under normal physiological conditions, PE is present at much higher levels in the inner leaflet but during apoptosis, it is upregulated in the outer leaflet. PE lipids externalized during apoptosis have been previously imaged using the lantibiotic-mediated imaging approach discussed above ^8,38^, enabling us to use it as an ideal positive control for our experiments.

To employ our technology for imaging PE externalization during apoptosis, we metabolically labelled PE with **19** using our standard protocol, and then triggered apoptosis by administering the calcium ionophore, A23187^39^ (**Figure 3A**). Subsequently, to image **19**-PE lipids exclusively in the outer leaflet of the plasma membrane, we employed a previously reported cell-impermeable cyclooctyne dye, AF647-DBCO^20,40^ (**Figure 3B**). We then treated these cells with duramycin-LC-biotin-streptavidin AF594, a PE-binding lantibiotic fluorescent probe^38^, and subjected the cells to confocal imaging. These experiments yielded fluorescent signals on the cell surface for both the AF647 and AF594 dyes (top panel, **Figure 3C**), which overlapped completely as evident in the line intensity plot depicted on the right. A control experiment wherein apoptosis was not triggered in **19**-administered cells yielded no fluorescence signals above background for both dyes (middle panel, **Figure 3C**). When apoptosis was triggered in cells that were not metabolically labelled with **19**, fluorescent intensities for the duramycin probe were obtained but not for the cyclooctyne probe (bottom panel, **Figure 3**).

**Figure 3.**
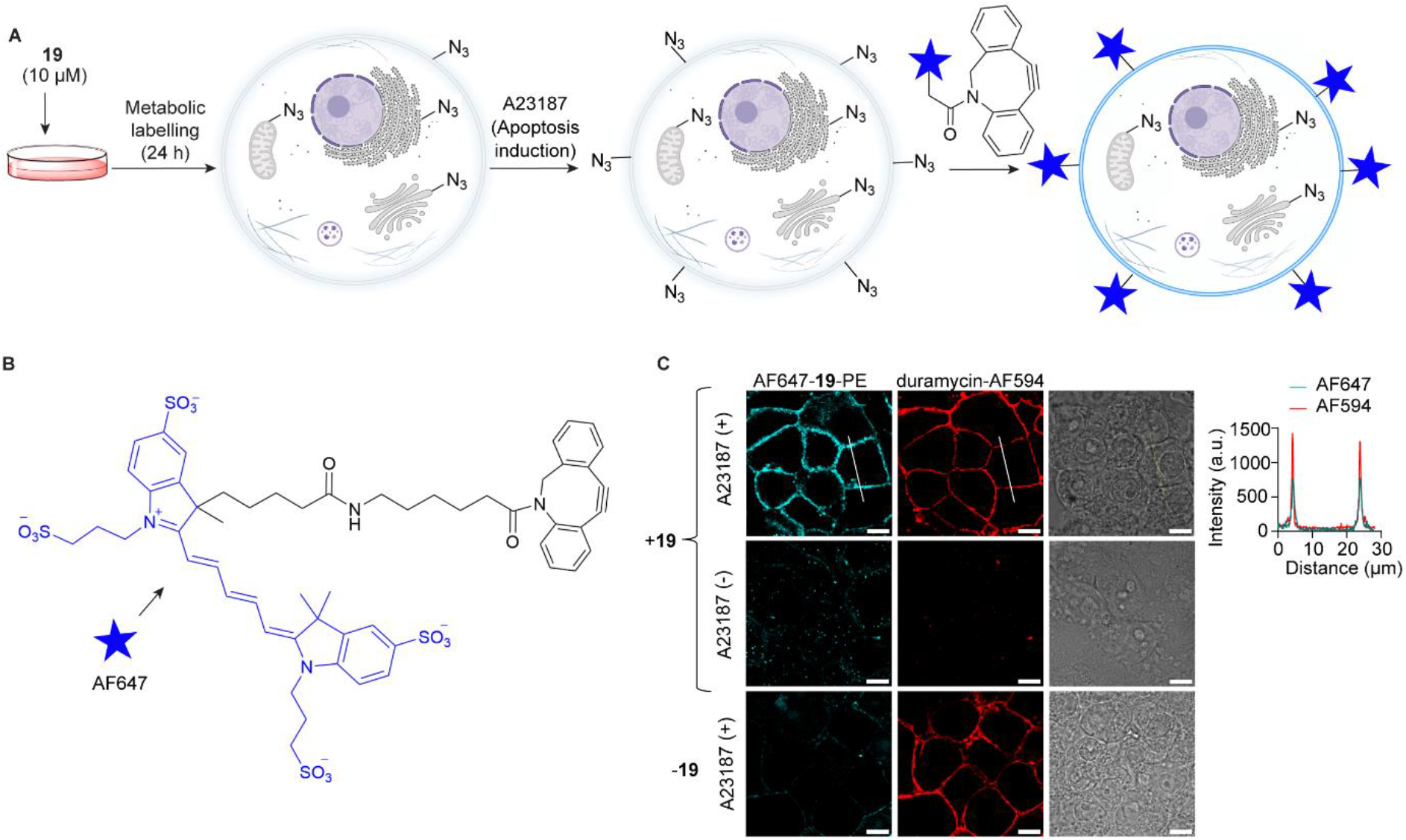
Visualizing PE externalization during apoptosis. A) A schematic of the workflow involving induction of apoptosis by administering the calcium ionophore A23187 to HeLa cells metabolically labelled with **19** followed by the use of cell impermeable cyclooctyne dye, AF647-DBCO to image externalized **19**-PE lipids. B) Structure of AF647-DBCO, C) Imaging data obtained. Top: **19**-administered cells treated with A23187 imaged with AF647-DBCO (left panel), the PE headgroup-binding probe, duramycin-AF594 (middle) and a bright field image (right). The plot on the right depicts line intensity fluorescence analysis performed across a line segment shown in white that traverses a cell. Middle: **19**-administered cells not treated with A23187 imaged after treatment with AF647-DBCO and duramycin-AF594. Bottom: Cells treated with A23187, and not **19** imaged as in the other two panels. Scale bars = 10 μm.

The similarity between these imaging results obtained with the AF647-DBCO dye that imaged externalized **19**-PE lipids, and the duramycin probe that imaged externalized native PE lipids establish that the dynamics of **19**-PE lipids are similar to those of native PE strongly suggesting that our probe is capable of faithfully recapitulating native PE dynamics in live mammalian cells.

#### Employing 19 to image PE in organelles in live cells

Having established **19** as a suitable probe for imaging PE lipids in live cells, we employed it to image PE within organellar membranes by performing co-localization experiments with organelle-selective dyes. We treated cells metabolically labelled with **19** with BCN-BODIPY to fluorescently label **19**-PE lipids, and then administered organelle trackers for performing Z stack-based co-localization imaging experiments. ER tracker yielded a high Pearson’s correlation coefficient (PCC) value of 0.8 (**Figure 4A**) establishing the presence of **19**-PE lipids in this organelle. Next, to investigate if **19**-PE lipids biosynthesized in the ER are transported to other cellular organelles, we performed co-localization experiments on lysosomes, Golgi and mitochondria. PCC analyses on the resulting confocal imaging data (**Figure 4B, C, D**) revealed that **19**-PE lipids were also present in these organelles establishing that **19**-PE is not only efficiently biosynthesized in mammalian cells but is also proficiently trafficked by the cellular machinery from the site of its biosynthesis to other organelles.

**Figure 4.**
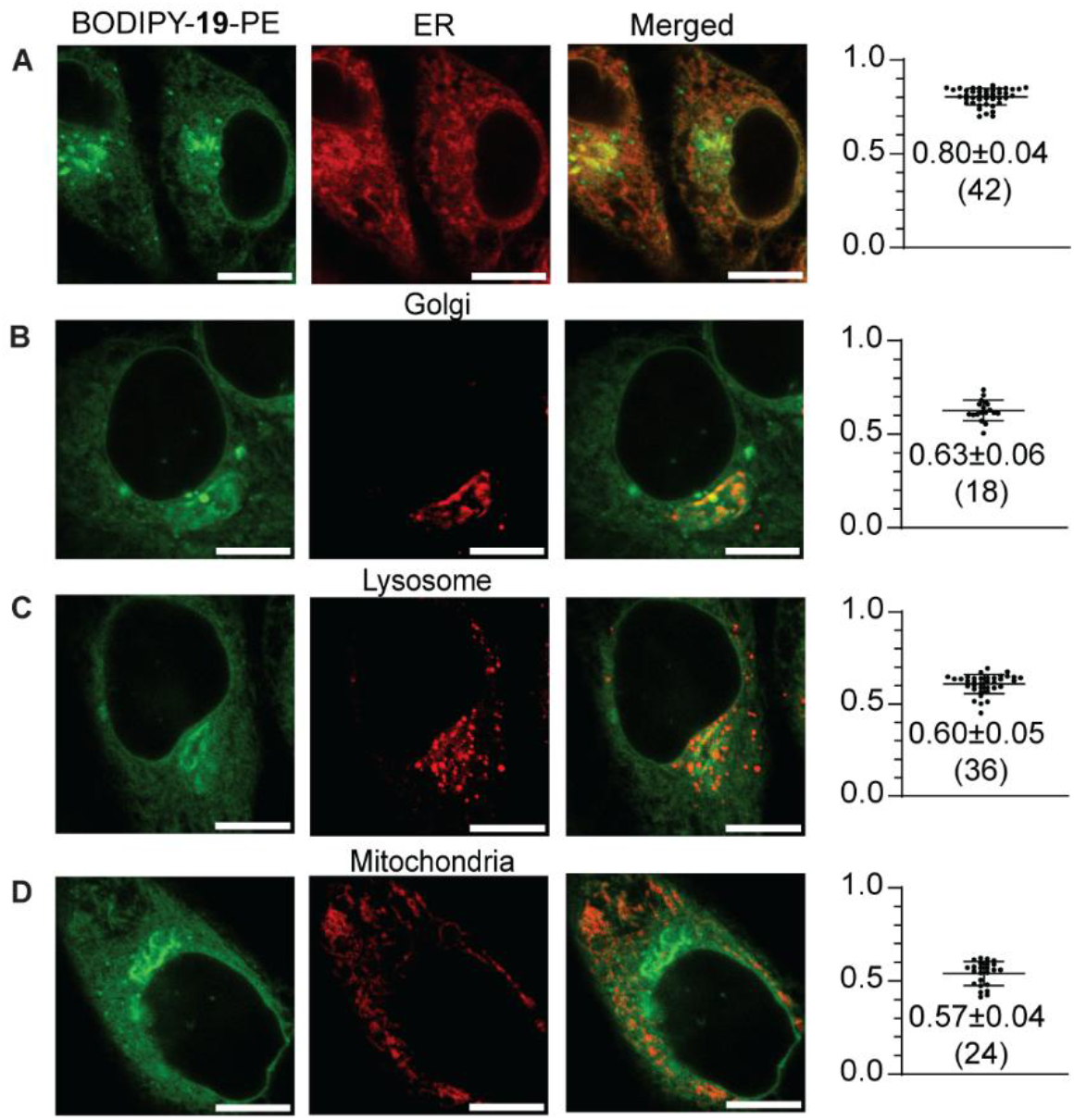
Imaging PE lipids in live cellular organelles. Co-localization experiments involving administering A) ER tracker, B) Golgi tracker, C) lysotracker, and D) mitotracker dyes to HeLa cells metabolically labelled with **19**. Pearson’s correlation values are depicted in plots on the right adjacent to representative images with the number of cells they were performed on within parenthesis. Scale bars = 10 μm.

Taken together, the lipidomic and imaging studies on **19** summarized in **Figures 2, 3**, and **4** strongly suggest that this probe is an ideal surrogate for ethanolamine with respect to PE biosynthesis, turnover, and cellular dynamics.

#### Our technology establishes VPS13A as a PE trafficking protein

Success in imaging Kennedy pathway-synthesized **19**-PE lipids in the mitochondria (**Figure 4D**) demonstrated unequivocally that these Kennedy pathway-synthesized lipids do indeed get transported from the ER to the mitochondria, and set the stage for using our technology to discover proteins that orchestrate lipid exchange between these two organelles. One such candidate protein is VPS13A, an ER–mitochondria contact site protein that is intimately associated with the neurodegenerative disorder, chorea-acanthocytosis, with one study reporting as many as 57 mutations in the *vps13a* gene in 43 unrelated patients afflicted with this disease^41^.

VPS13A has been recently demonstrated to transport PC lipids from the ER to the mitochondria^19^, and although it has been previously reported that VPS13A binds to PE^42^, no reports on its ability to transport PE are published to the best of our knowledge. We tested this hypothesis using our technology.

To specifically image **19**-PE in the mitochondria, we replaced BCN-BODIPY, a pan-organelle diffusing cyclooctyne dye that we had employed for organellar imaging (**Figure 4**), with Cy5-DBCO (**Figure 5A,B**), a mitochondria-localizing cyclooctyne dye that was recently employed to image PC lipids within the mitochondria^20,40^. The positively charged Cy5 moiety of this dye drives it into the negative membrane potential environment of the mitochondria where it undergoes click chemistry with azide-tagged lipids. Subsequently, the unreacted dye is quenched by adding Cy7-N_3_ followed by washing with high K^+^ media in presence of valinomycin and the mitochondrial proton gradient disrupting agent, CCCP.

**Figure 5.**
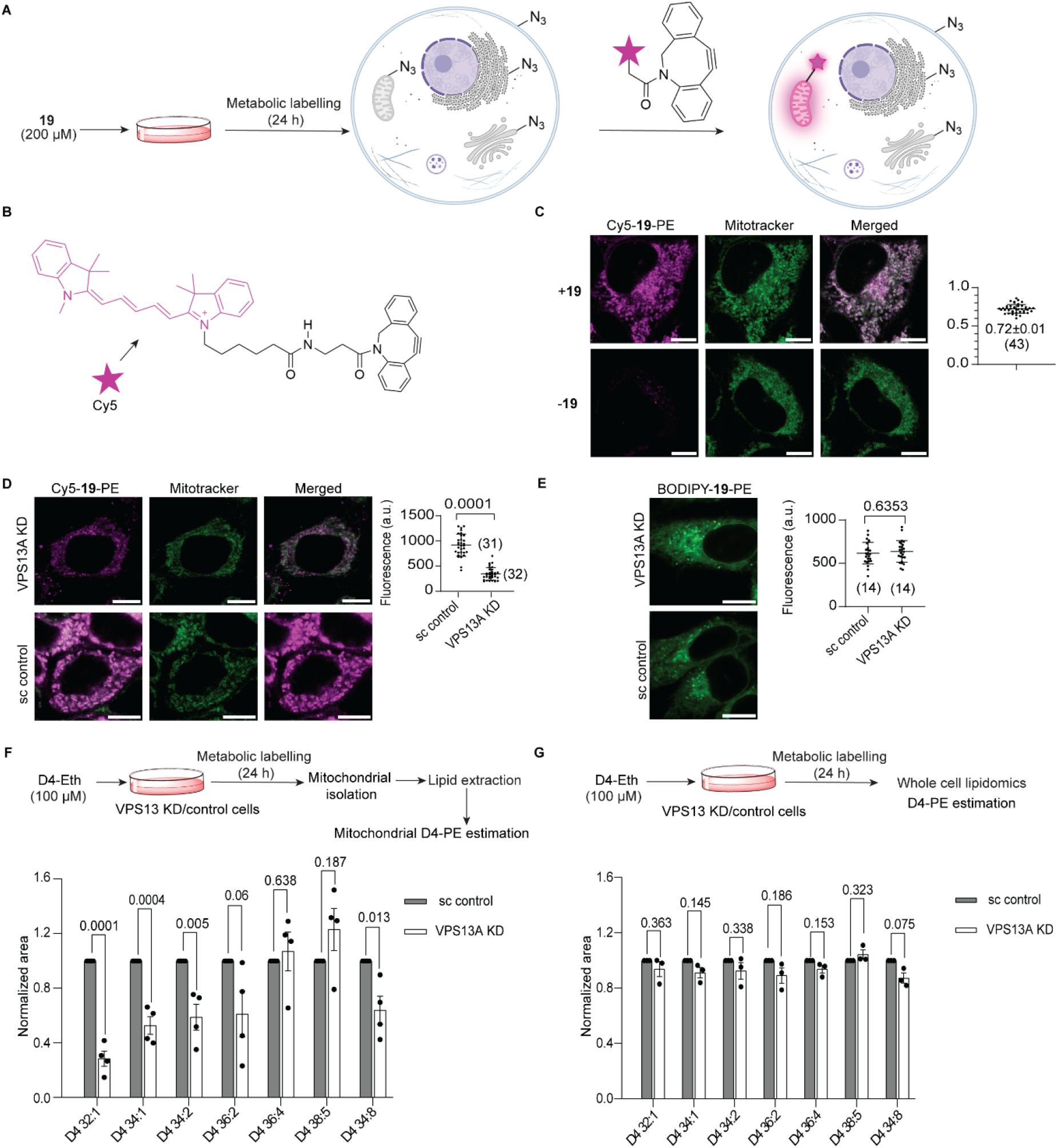
Mitochondria-specific imaging of PE lipids and VPS13A knockdown studies. A) Schematic depicting the use of the Cy5-DBCO dye for imaging **19**-PE lipids specifically within the mitochondria. B) Structure of Cy5-DBCO. C) Imaging results on HeLa cells either treated with **19** (top) or not (bottom) by employing Cy5-DBCO to image **19**-PE (left) and mitotracker green to image mitochondria (middle). Co-localization analysis results are depicted on the plot on the right. D) Imaging VPS13A knockdown (KD) HeLa cells (top) and control cells administered with a scrambled construct (bottom). Fluorescence intensity plots are depicted on right. E) Imaging using the pan-organelle diffusing dye, BCN-BODIPY in **19**-administered VPS13A KD and control HeLa cells with representative images depicted on the left and statistical analysis on the right. F) MRM lipidomics comparing the levels of prominent PE lipids present in the mitochondria of VPS13A KD and control cells metabolically labelled with D4-Eth. G) MRM lipidomics on the same PE species analyzed above present in lipids isolated from D4-Eth administered VPS13A KD and control cells (whole cell lipidomes). Scale bars = 10 μm.

Our imaging experiments on **19**-administered HeLa cells yielded Cy5 fluorescence intensities that demonstrated a high PCC value of 0.72 (**Figure 5C**, top panel) with the mitotracker dye, and control experiments where cells not administered with **19** were treated with Cy5-DBCO did not yield any fluorescent signals above background (**Figure 5C**, bottom panel).

To investigate whether the VPS13A protein contributes to transporting **19**-PE lipids from the ER to the mitochondria, we knocked down VPS13A using shRNA, achieving ∼47% knockdown as demonstrated by Western blotting (**Figure S12**), and subjected these cells to the imaging workflow summarized above. The Cy5 fluorescence intensities measured in knockdown cells were ∼3-fold lower than that in the control cells treated with a scrambled shRNA construct, as depicted in the representative images and the statistical analysis over more than 30 cells (**Figure 5D**). These results suggest that trafficking of PE from the ER to the mitochondria is disrupted in the VPS13A knockdown cells. An alternate explanation of these results is that VPS13A knockdown disrupts PE biosynthesis in the ER resulting in lower levels of PE available for transport to the mitochondria. To address this possibility, we employed the pan-organelle diffusing dye BCN-BODIPY instead of Cy5-DBCO to image **19**-PE lipids and obtained indistinguishable fluorescence intensities between the control and knockdown cells demonstrating that VPS13A knockdown does not affect PE lipid biosynthesis (**Figure 5E**).

To evaluate whether the results above obtained for **19**-PE lipids translate to native PE lipids, we performed D4-ethanolamine labelling experiments on both VPS13A knockdown cells as well as control cells. Subsequently, we purified mitochondria from these cells by optimizing a percoll gradient centrifugation workflow (**Figure S3** and **S13**), isolated mitochondrial lipids, and performed MRM lipidomics to quantitate seven D4-PE lipids that our MS experiments revealed were present in high abundance in the mitochondria (**Figure 5F**). These experiments revealed that mono- and di-unsaturated PE lipids: PE 32:1, PE 34:1, PE 34:2 and PE 36:2, were significantly depleted in the mitochondria of the knockdown cells whereas the levels of polyunsaturated fatty acid-containing PE lipids: PE 36:4 and PE 38:5 were not significantly altered. These results are consistent with the observation that the Kennedy pathway preferably synthesizes mono-and di-unsaturated PE whereas the mitochondrial PISD-mediated PE synthesis primarily generates polyunsaturated fatty acid-containing PE lipids^34^. Since VPS13A transports PE lipids from the ER to the mitochondria, the levels of ER-synthesized PE lipids (mono- and di-unsaturated species) would be expected to be depleted in the mitochondria of VPS13A knockdown cells, and not the polyunsaturated ones, in line with the results of our experiments. Furthermore, whole cell lipidomics on D4-ethanolamine treated cells did not reveal any significant alterations in the cellular levels of these lipids (**Figure 5G**) demonstrating that VPS13A knockdown does not impact PE biosynthesis. Taken together, our lipidomics data support the imaging data and establish that VPS13A transports PE lipids from the ER to the mitochondria.

### Conclusions

We have developed a robust technology for imaging PE lipids within live cells. This technology leverages the promiscuity of the Kennedy pathway with respect to accepting structurally diverse ethanolamine derivatives as substrates for PE biosynthesis, enabling us to develop an azido ethanolamine probe that efficiently generated azido PE lipids when administered to cells at concentrations similar to those of native ethanolamine. Crucially, this probe did not form methylated PE lipids when administered to mammalian cells, enabling exclusive labelling (and therefore, cellular imaging) of PE lipids and not their methylated versions (including PC). We employed this probe to successfully visualize PE in various cellular organelles, and image PE externalization during apoptosis in live cells. Furthermore, by employing our technology together with a mitochondria-targeting cyclooctyne dye, we discovered that the therapeutically attractive VPS13A protein traffics PE lipids from the mitochondria to the ER. We anticipate that our technology will be employed for addressing various aspects of PE biology and shed light on its roles in cellular physiology and disease pathophysiology.

## Acknowledgements

J. K. thanks IISER Bhopal and DBT/Wellcome Trust India Alliance Fellowship [grant number IA/S/23/2/506977] for funding. The imaging data shown was obtained from the imaging facility at IISER Bhopal supported by DST-FIST. S. B., S. S. and B. T. thank IISER Bhopal for fellowship. A. D. S. thanks IISER Bhopal and CSIR for fellowship. Manas and Shreya thank DBT and CSIR respectively for fellowships. T. A. K. and G. B. thank IISER Bhopal for providing Institute post-doctoral fellowships. We thank former Kalia lab PhD student, Aditi Dixit, for insightful discussions.

