## Supporting Information for "A metabolic labelling-based lipid imaging technology establishes VPS13A as a phosphatidylethanolamine lipid transfer protein"

[2]Department of Chemistry

Professor, Department of Biological Sciences

Associate Faculty, Department of Chemistry

DBT Wellcome India Alliance Senior Fellow

**Table of Contents**


| Section No. | Contents | Pg. No. | Table, figures, and schemes in the sub-section |
|---|---|---|---|
| I | General information | 3 | - |
| II | Kennedy pathway for PE biosynthesis | 4 | Figure S1 |
| III | Synthetic schemes and procedures | 4–18 | Schemes S1–S11 |
| IV | Antibodies used in this study | 18 | Table S1 |
| V | Lipidomics procedures | 18–21 | Table S2 |
| VI | Imaging procedures | 21–23 | Table S3 |
| VII | Procedure for TLC analysis followed by MS analysis on fluorescent spots | 24–26 | Figure S2, Scheme S12 |
| VIII | Procedure for cytotoxicity assay on HeLa cells treated with **19** | 27 | |
| IX | shRNA-mediated knockdown of VPS13A | 27–28 | |
| X | Mitochondria isolation using percoll gradient | 28–29 | Figure S3 |
| XI | Lipidomics data | 29–55 | Table S4, Figures S4–S9 |
| XII | Data for cytotoxicity assay on HeLa cells treated with **19** | 56 | Figure S10 |
| XIII | Data for MS analysis on fluorescent TLC spot denoted by "#" in Figure 2B of the main text | 56 | Figure S11 |
| XIV | Western blot on VPS13A knockdown cells | 56 | Figure S12 |
| XV | Western blot analysis on purified mitochondrial fractions obtained by performing percoll gradient fractionation on VPS13AKD and control HeLa cells | 57 | Figure S13 |
| XVI | References | 58 | - |
| XVII | $^{1}H$ and $^{13}C$ NMR spectra | 59–80 | - |

### I. General information

*Organic synthesis and reagents:* All organic synthesis reactions were performed in oven-dried glassware. Chemical reagents for organic synthesis were purchased either from Sigma-Aldrich, Alfa Aesar or Tokyo Chemical Industry Co. Ltd. and were used without purification. Distilled water was used for reaction work-ups, and ultrapure Type 1 water was used for buffer preparations and LC-MS experiments. The progress of chemical reactions was monitored by thin layer chromatography (TLC) using 0.25 mm Merck precoated (60 F254) silica gel plates. For visualization of TLC spots, UV light of wavelength 254 nm (for UV active compounds) and 365 nm (for fluorescent compounds) was used. In some cases, TLC spots were visualized using ninhydrin and phosphomolybdic acid (PMA) staining solutions. Purifications were performed using column chromatography on silica gel (100-200 mesh). $^{1}$H and $^{13}$C NMR spectra were recorded on Bruker's AVANCE-III 500 MHz and 400 MHz NMR spectrometers. Chemical shifts are reported as parts per million (δ) relative to tetramethylsilane (TMS) as internal standard and coupling constants (J values) in Hertz (Hz). Multiplicities are indicated as follows: s (singlet), d (doublet), t (triplet), q (quartet), dd (doublet of doublet), bs (broad singlet), td (triplet of a doublet) and m (multiplet). High-resolution mass spectra (HRMS) were recorded on LCMS High Resolution Bruker's MicrOTOF-Q III and Agilent 6546 HRLC-QTOF ESI.

*Cell culture:* HeLa and HEK293 cells were maintained in complete medium consisting of Dulbecco's Modified Eagle Medium (DMEM) supplemented with 10% fetal bovine serum (FBS), 100 units/mL penicillin, and 100 µg/mL streptomycin, in a humidified $CO_2$ incubator maintained at 5% $CO_2$ and 37 °C. All treatments and incubations of mammalian cells were performed under the same conditions.

*Lipidomics:* Lipidomics was performed on an ion trap triple quadrupole mass spectrometer (QTRAP® 4500 system from AB SCIEX) equipped with an Exion LC. All lipidomics mass spectra were analysed on the Analyst software and lipid species were identified by using the LipidView software. PE lipid standard (17:0/20:4) was purchased from Avanti polar lipids. 3-Deazaadenosine (DZA) and D4-ethanolamine were purchased from Santa Cruz Biotechnology and Cambridge Isotope respectively. Dulbecco's modified Eagle medium (DMEM), phosphate-buffered saline (PBS), Fetal bovine serum (FBS), and 0.05% trypsin-EDTA were purchased from Thermo Fisher, polyethylenimine (PEI) linear (MW 25,000) from Kyfora Bio, the calcium Ionophore A23187 from Sigma, and meclizine dihydrochloride from Merck.

*Imaging:* Confocal imaging was performed on an Olympus FV3000 confocal laser scanning microscope equipped with 40XO 1.4 NA and 100XO 1.4 NA UPLSAPO objectives, 488, 561, 594 and 640 nm solid-state lasers, and PMT detectors with a Tokai Hit stage-top incubator. Image analysis was performed using FIJI/ImageJ software. The shRNA construct employed for knocking down the VPS13A protein in HeLa cells was purchased from an in-house facility at the Department of Biological Sciences, IISER Bhopal. Details of antibodies used for western blots are provided in **Table S1**. All the western blots were visualized using a ChemiDoc imaging system (Bio-Rad).

### II. Kennedy pathway for PE biosynthesis

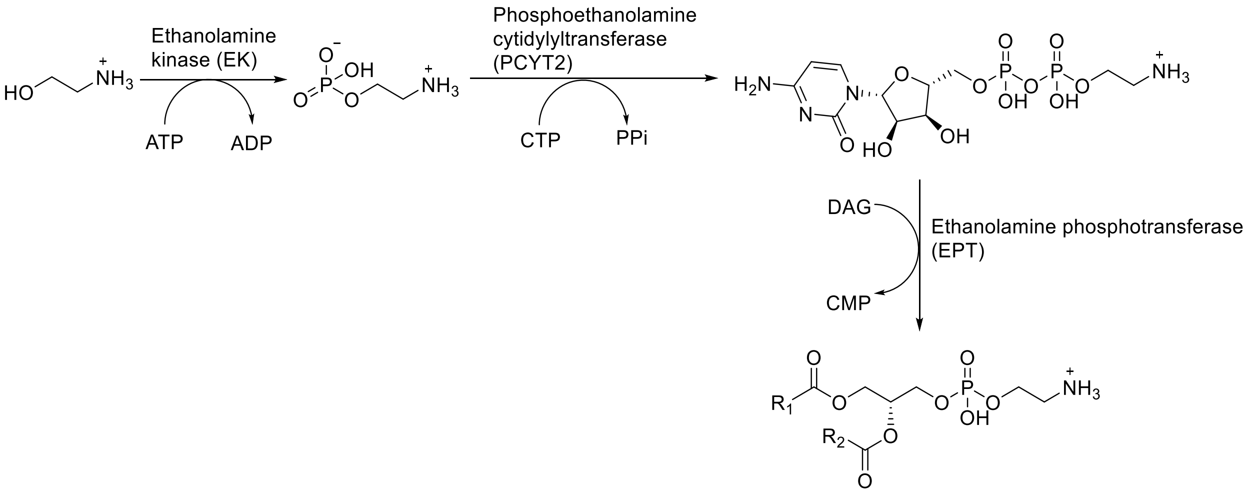


**Figure S1:** Kennedy pathway for PE biosynthesis

III. **Synthetic schemes and procedures**

Ethanolamine analogs **6**, **15**, **16**, and **18** were purchased from Sigma Aldrich and used without purification. Compounds **9**, **10**, and the dyes, azido fluorescein and alkynyl rhodamine were synthesized using previously established procedures in our laboratory[1].

**Scheme S1.** Synthetic scheme for **1**.

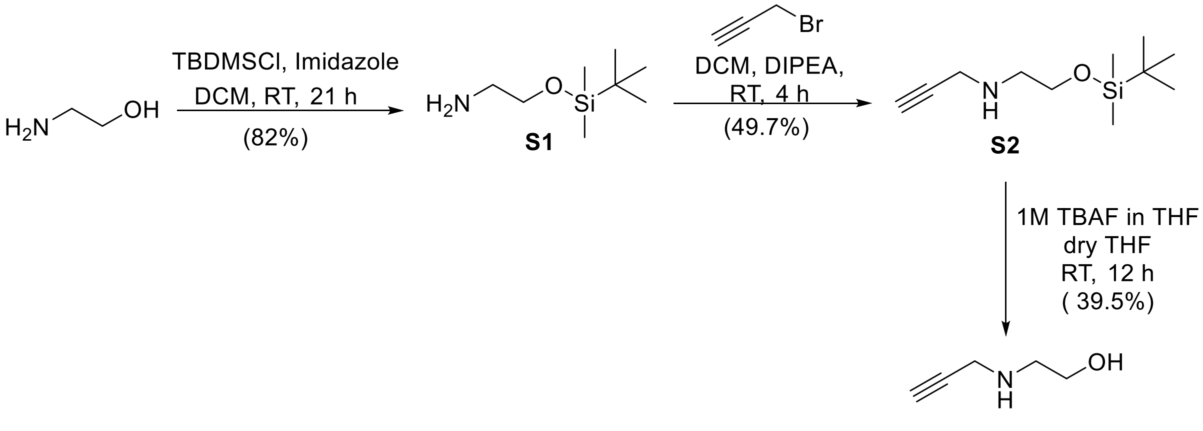


**2-(prop-2-yn-1-ylamino)ethan-1-ol (1)**

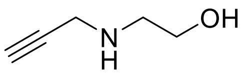

**1**

**S1** and **S2** were synthesized according to a previously reported procedure[1]. To a round-bottomed flask containing **S2** (860 mg, 4.03 mmol, 1 eq.) in anhydrous tetrahydrofuran (20 mL), tetrabutylammonium fluoride (6.05 mL of a 1 M solution in tetrahydrofuran, 6.05 mmol, 1.5 eq.) was added and the resultant reaction mixture was stirred at room temperature for 18 h. After removing the solvent under reduced pressure, the residue was subjected to silica gel column chromatography (10% methanol in ethyl acetate) to obtain **1** as a yellow liquid (212 mg, 39.5%).

$^{1}H$ NMR (500 MHz, $CDCl_3$): δ 3.69 (t, J = 5.12 Hz, 2H), 3.47 (d, J = 2.42 Hz, 2H), 3.25 (bs, 2H), 2.88 (t, J = 5.12 Hz, 2H), 2.25 (t, J = 2.42 Hz, 1H).

$^{13}C$ NMR (125 MHz, $CDCl_3$): δ 81.26, 72.23, 60.62, 49.98, 37.71.

HRMS (ESI): $C_5H_9NO$; Calculated mass, $[M+H]^+$100.0757; Observed mass, $[M+H]^+$ 100. 0769.

**Scheme S2.** Synthetic scheme for **2**.

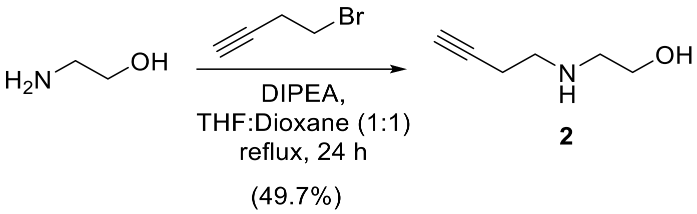


**2-(but-3-yn-1-ylamino)ethan-1-ol (2)**

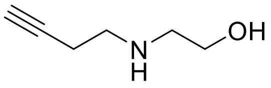


**2**

To a round-bottomed flask containing ethanolamine (229 mg, 3.75 mmol, 1 eq.) in tetrahydrofuran (10 mL) and dioxane (10 mL), *N,N*-diisopropylethylamine (0.72 mL, 4.12 mmol, 1.1 eq.) was added and cooled to 0 °C. Propargyl bromide (352 μL of a 80% (*w/w*) solution, 3.75 mmol, 1 eq.) was added dropwise to this mixture. After stirring at 0 °C for 30 min., the mixture was warmed to room temperature and refluxed for 24 h. The resulting solution was washed with saturated sodium bicarbonate solution (1 × 100 mL) and dichloromethane (2 × 50 mL). The organic layer was then dried by adding anhydrous sodium sulfate, concentrated under reduced pressure, and the crude residue was subjected to silica gel column chromatography (1% triethylamine in ethyl acetate) to obtain **2** as a yellow-coloured oil (198 mg, 49.7%).

$^{1}H$ NMR (400 MHz, $CDCl_3$): δ 3.65 (t, J = 5.18 Hz, 2H), 2.83-2.80 (m, 4H), 2.41 (td, J = 6.50, 2.59 Hz, 2H), 2.01 (t, J = 2.59 Hz, 1H).

$^{13}C$ NMR (125 MHz, CDCl3) δ 80.86, 70.50, 59.42, 50.23, 46.80, 18.25.

HRMS (ESI): $C_4H_{11}NO^+$; Calculated mass, $[M+H]^+$ 114.0913; Observed mass, $[M+H]^+$ 114.0919.

**Scheme S3.** Synthetic scheme for **3**.

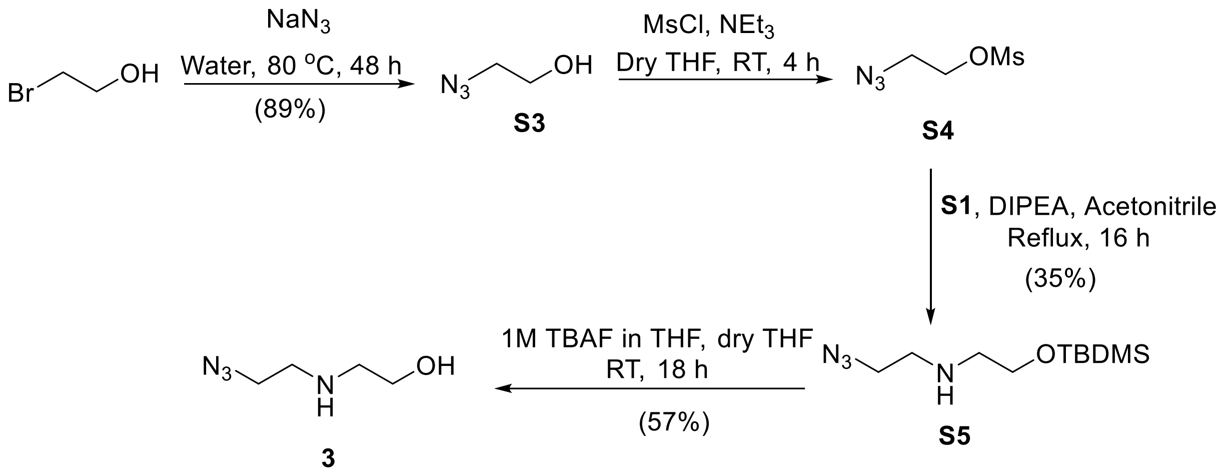


**2-((2-azidoethyl)amino)ethan-1-ol (3)**

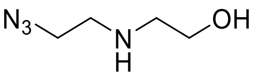

**3**

**S3, S4**, and **S5** were synthesized according to previously reported procedures[1]. In a round-bottomed flask containing solution of **S5** (1.0 g, 4.09 mmol, 1 eq.) in anhydrous tetrahydrofuran (20 mL), tetrabutylammonium fluoride (6.13 mL of a 1 M solution in tetrahydrofuran, 6.13 mmol, 1.5 eq.) was added and the resulting reaction mixture was stirred at room temperature for 18 h. After removing the solvent under reduced pressure, the residue was subjected to silica gel column chromatography (10% methanol in ethyl acetate containing 1% triethylamine) to obtain **3** as a yellow liquid (301 mg, 57%).

$^1$H NMR (500 MHz, $CDCl_3$): δ 3.66 (t, J = 5.16 Hz, 2H), 3.44 (t, J = 5.66 Hz, 2H), 2.85 (t, J = 5.67 Hz, 2H), 2.82 (t, J = 5.16 Hz, 2H), 1.92 (s, 2H).

$^{13}$C NMR (125 MHz, $CDCl_3$): δ 60.92, 51.39, 50.88, 48.23.

HRMS (ESI): $C_4H_{10}N_4O$; Calculated mass, $[M+H]^+$ 131.0927; Observed mass, $[M+H]^+$ 131.0923.

**Scheme S4:** Synthetic scheme for **4** and **12**.

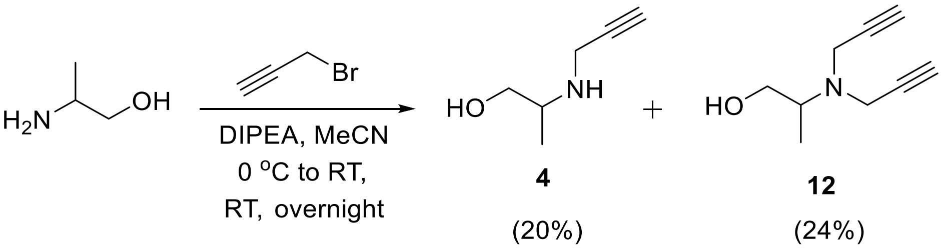

**2-(prop-2-yn-1-ylamino)propan-1-ol (4), 2-(di(prop-2-yn-1-yl)amino)propan-1-ol (12)**

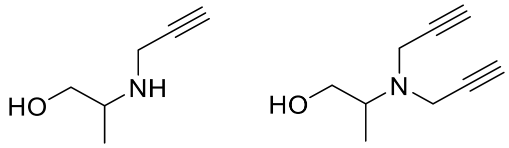


**4** **12**

To a round-bottomed flask containing 2-aminopropan-1-ol (212 μL, 2.6 mmol, 1 eq.) in acetonitrile (2 mL), *N,N*-diisopropylethylamine (462 μL, 6.65 mmol, 1 eq.) was added, and after cooling this solution to 0 °C, propargyl bromide (295 μL of a 80% (*w/w*) solution in toluene, 6.65 mmol, 1 eq.) was added dropwise. After stirring the mixture at 0 ºC for 30 min, it was warmed up to room temperature and stirred overnight. The resulting solution was washed with saturated sodium bicarbonate solution (1 × 60 mL) and dichloromethane (3 × 20 mL). The organic layer was then dried by adding anhydrous sodium sulfate, concentrated under reduced pressure and the crude residue was subjected to silica gel column chromatography (1% triethylamine in ethyl acetate) to give **4** (monosubstituted) as a yellow oil (61.5 mg, 20%) and **12** (disubstituted) as a yellow oil (97.9 mg, 24.3%).

**4**: $^{1}$H NMR (500 MHz, $CDCl_3$): δ 3.62 (dd, J = 10.79, 3.85 Hz, 1H), 3.46 (qd, J = 17.11, 2.43 Hz, 2H), 3.32 (dd, J = 10.80, 6.50 Hz, 1H), 3.03-2.97 (m, 1H), 2.22 (t, J = 2.43 Hz, 1H), 1.99 (bs, 1H), 1.93 (bs, 1H), 1.07 (d, J = 6.46 Hz, 3H).

$^{13}$C NMR (125 MHz, $CDCl_3$): δ 82.19, 71.57, 65.65, 53.15, 35.75, 16.92(s).

HRMS (ESI): $C_6H_{11}NO$; Calculated mass, $[M+H]^+$ 114.0913; Observed mass, $[M+H]^+$114.0937.

**12**: $^{1}$H NMR (500 MHz, $CDCl_3$): δ 3.53 (d, J = 2.38 Hz, 4H), 3.52-3.47 (m, 1H), 3.38 (ddd, J = 11.10, 8.52, 2.57 Hz, 1H), 3.13-3.07 (m, 1H), 2.69 (dd, J = 8.22, 2.74 Hz, 1H), 2.24 (t, J = 2.31 Hz, 2H), 1.08 (d, J = 6.73 Hz, 3H).

$^{13}$C NMR (125 MHz, $CDCl_3$): δ 80.14, 73.01, 63.60, 58.23, 38.91, 12.23.

HRMS (ESI): $C_9H_{13}NO$; Calculated mass, $[M+H]^+$ 152.1070; Observed mass, $[M+H]^+$ 152.1085.

**Scheme S5.** Synthetic scheme for **5** and **13**

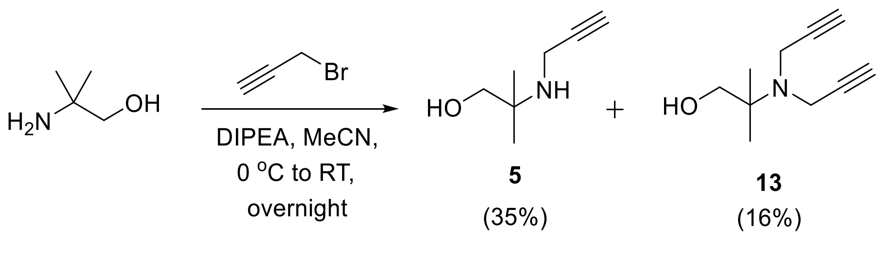

**2-methyl-2-(prop-2-yn-1-ylamino)propan-1-ol (5), 2-(di(prop-2-yn-1-yl)amino)-2-methylpropan-1-ol (13)**

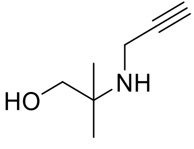


**5**

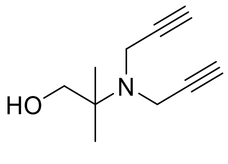


**13**

To a round-bottomed flask containing 2-amino-2-methylpropan-1-ol (400 mg, 4.48 mmol, 1 eq.) in acetonitrile (4 mL), *N,N*-diisopropylethylamine (780 μL, 4.48 mmol, 1 eq.) was added and after cooling this solution to 0 °C, propargyl bromide (499 μL of a 80% (*w/w)* solution in toluene, 4.48 mmol, 1 eq.) was added dropwise. After stirring the mixture at 0 °C for 30 min, it was warmed up to room temperature and stirred overnight. The resulting solution was washed with saturated sodium bicarbonate solution (1 × 60 mL) and dichloromethane (2 × 30 mL). The organic layer was then dried by adding anhydrous sodium sulfate, concentrated under reduced pressure and the crude residue was subjected to silica gel column chromatography (1% triethylamine in ethyl acetate) to obtain **5** (monosubstituted) as a yellow oil (199.6 mg, 34.9%) and **13** (disubstituted) as a yellow oil (119 mg, 16%).

**5**: $^{1}$H NMR (500 MHz, $CDCl_3$): δ 3.37 (d, J = 2.48 Hz, 2H), 3.34 (s, 2H), 2.21 (t, J = 2.48 Hz, 1H), 1.10 (s, 6H).

$^{13}$C NMR (125 MHz, $CDCl_3$): δ 83.51, 71.01, 68.78, 54.31, 31.67, 23.97.

HRMS (ESI): $C_7H_{13}NO$; Calculated mass, $[M+H]^+$ 128.1070; Observed mass,$[M+H]^+$ 128.1072.

**13**: $^{1}$H NMR (500 MHz, $CDCl_3$): δ 3.62 (d, J = 2.30 Hz, 4H), 3.30 (d, J = 5.16 Hz, 2H), 2.68 (t, J = 5.46 Hz, 1H), 2.22 (t, J = 2.06 Hz, 2H), 1.19 (s, 6H).

$^{13}$C NMR (125 MHz, $CDCl_3$): δ 81.12, 72.69, 68.35, 58.70, 36.30, 22.04.

HRMS (ESI): $C_{10}H_{15}NO$; Calculated mass, $[M+H]^+$ 166.1226; Observed mass, $[M+H]^+$ 166.1230.

**Scheme S6.** Synthetic scheme for **7**.

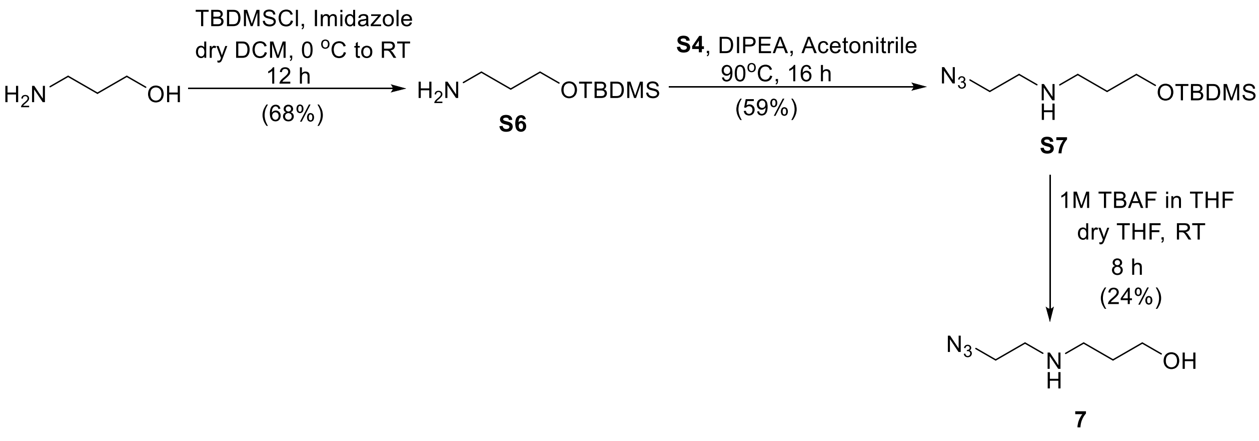

**3-((tert-butyldimethylsilyl)oxy)propan-1-amine (S6)**

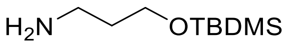

**S6**

A round-bottomed flask containing 3-amino-1-propanol (1 g, 13.3 mmol, 1 eq) and imidazole (0.996 g, 14.63 mmol, 1.1 eq) was subjected to high vacuum and then dissolved in anhydrous dichloromethane (20 mL) under nitrogen atmosphere and cooled to 0 ºC. To this mixture, a solution of tert-butyldimethylsilyl chloride (2.2 g, 14.63 mmol, 1.1 eq) in anhydrous dichloromethane (20 mL) at 0 ºC was added and stirred at room temperature for 12 h. The reaction was quenched with saturated bicarbonate (1 × 100 mL) and washed with saturated ammonium chloride (1 × 100 mL) and dichloromethane (3 × 20 mL). The organic layer was dried by adding anhydrous sodium sulfate and concentrated to afford **S6** as a pale-yellow oil (1.7 g, 68%).

$^1$H NMR (500 MHz, $CDCl_3$) δ 3.69 (t, *J* = 6.0 Hz, 2H), 2.79 (t, *J* = 6.7 Hz, 2H), 1.86 – 1.45 (m, 2H), 0.88 (s, 9H), 0.04 (s, 6H).

$^{13}$C NMR (125 MHz, $CDCl_3$) δ 61.25, 39.45, 36.42, 25.94, 25.70, 18.30, -5.36.

HRMS (ESI): $C_9H_{23}NOSi$; Calculated mass, $[M+H]^+$ 190.1622; Observed mass, $[M+H]^+$ 190.1622.

**N-(2-azidoethyl)-3-((tert-butyldimethylsilyl)oxy)propan-1-amine (S7)**

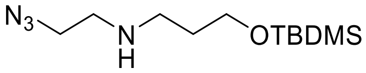

**S7**

In a round-bottomed flask, azido ethanol (452 mg, 5.2 mmol, 1 eq) was taken and subjected to high vacuum followed by addition of anhydrous triethylamine (1.05 g, 10.4 mmol, 2 eq) and anhydrous tetrahydrofuran (50 mL) and cooled to 0 ºC, followed by slow addition of methane sulfonyl chloride (1.19 g, 10.4 mmol, 2 eq) under nitrogen atmosphere. After 30 min, the mixture was allowed to warm to room temperature and stirred for 14 h. The reaction mixture was washed with 1 N sodium hydroxide (1 × 60 mL) and dichloromethane (4 × 20 mL). The organic layer was dried by adding anhydrous sodium sulfate, concentrated, and added to a solution of **S6** (1 g ,5.2 mmol, 1 eq) and *N,N*-diisopropylethylamine (0.672 g, 5.2 mmol, 1 eq) in acetonitrile (20 mL) and refluxed for 16 h at 90 ºC. The resulting reaction mixture was washed with saturated bicarbonate (1 × 80 mL) and dichloromethane (4 × 20 mL). The product was purified via silica gel column chromatography (4% methanol in dichloromethane) to afford **S7** as a yellow viscous oil (800 mg, 59%).

$^1$H NMR (500 MHz, $CDCl_3$) δ 3.75 (t, *J* = 5.9 Hz, 2H), 3.52 (t, *J* = 5.3 Hz, 2H), 2.84 (dt, *J* = 13.2, 6.3 Hz, 4H), 1.84 – 1.71 (m, 2H), 0.92 (s, 9H), 0.09 (s, 6H).

$^{13}$C NMR (125 MHz, $CDCl_3$) δ 61.78, 51.21, 48.52, 47.10, 29.70 (s), 25.92, 18.31, -5.38.

HRMS (ESI): $C_{11}H_{26}N_4OSi$; Calculated mass, $[M+H]^+$ 259.1949; Observed mass, $[M+H]^+$ 259.1939.

**3-((2-azidoethyl)amino)propan-1-ol (7)**

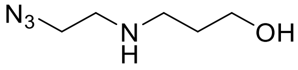

**7**

A round-bottomed flask charged with **S7** (530 mg, 2 mmol, 1 eq) was subjected to high vacuum and dissolved in dry tetrahydrofuran (20 mL), followed by addition of 1 M TBAF in tetrahydrofuran (0.784 g, 3 mmol, 1.5 eq) and stirred at room temperature for 8 h. The volatiles were removed under reduced pressure and purified via silica gel column chromatography (7% methanol in dichloromethane) to afford **7** as a yellow viscous liquid (73 mg, 24%).

$^1$H NMR (500 MHz, $CDCl_3$) δ 3.81 (t, *J* = 5.4 Hz, 2H), 3.47 (t, 2H), 2.90 (t, *J* = 5.8 Hz, 2H), 2.82 (t, 2H), 1.85 – 1.63 (m, 2H).

$^{13}$C NMR (125 MHz, $CDCl_3$) δ 63.86, 51.05, 49.27, 48.25, 30.72.

HRMS (ESI): $C_5H_{12}N_4O$; Calculated mass, $[M+Na]^+$ 167.0903; Observed mass, $[M+Na]^+$167.0904.

**Scheme S7.** Synthetic scheme for **8**.

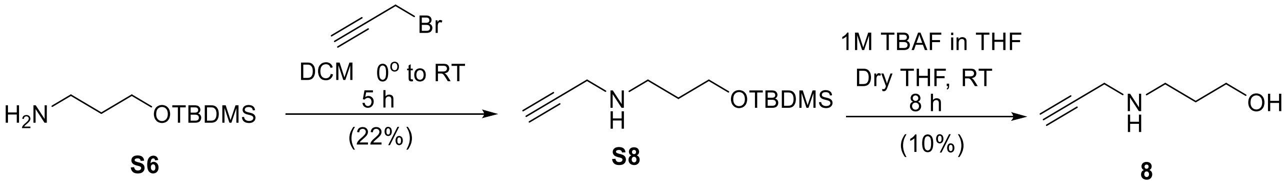


**N-(3-((tert-butyldimethylsilyl)oxy)propyl)prop-2-yn-1-amine (S8)**

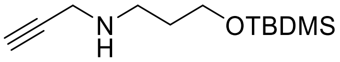

**S8**

To a round-bottomed flask containing **S6** (1 g, 5.28 mmol, 1 eq) and *N,N*-diisopropylethylamine (0.341 g, 2.64 mmol, 0.5 eq), dry dichloromethane (50 mL) was added and cooled to 0 °C. A solution of propargyl bromide (392 µL 80% (*w/w)* solution in toluene, 2.64 mmol, 0.5 eq) in dichloromethane (30 mL) was added slowly using dropping funnel at 0 °C and stirred for 30 min. The mixture was then allowed to warm up to room temperature and stirred for 5 h. Subsequently, the solution was washed with saturated sodium bicarbonate (1 × 70 mL) and dichloromethane (2 × 30 mL) and the organic layer was dried by adding anhydrous sodium sulfate, concentrated and purified via silica gel column chromatography (4% methanol in dichloromethane) to obtain **S8** as a yellow liquid (269 mg, 22%).

$^1$H NMR (500 MHz, $CDCl_3$) δ 3.70 (t, 2H), 3.42 (d, 2H), 2.79 (t, 2H), 2.20 (t, 1H), 1.74 – 1.69 (m, 2H), 0.89 (s, 9H), 0.05 (s, 6H).

$^{13}C$ NMR (125 MHz, $CDCl_3$) δ 82.09, 71.31, 61.76, 46.13 (s), 38.20, 32.5, 29.70, 25.94, 18.28, -5.36.

HRMS (ESI): $C_{12}H_{25}NOSi$; Calculated mass, $[M+H]^+$ 228.1778; Observed mass, $[M+H]^+$ 228.1781.

**3-(prop-2-yn-1-ylamino)propan-1-ol (8)**

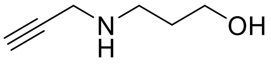

**8**

A round-bottomed flask charged with **S8** (269 mg, 1.18 mmol, 1 eq) was subjected to high vacuum and dissolved in dry tetrahydrofuran (15 mL), followed by addition of 1 M TBAF in tetrahydrofuran (462 mg, 1.77 mmol, 1.5 eq) under nitrogen atmosphere and stirred at room temperature for 8 h. The volatiles were removed under reduced pressure and purified via silica gel column chromatography (6% methanol and dichloromethane) to afford **8** as a yellow viscous liquid (13 mg, 10%).

$^{1}H$ NMR (500 MHz, $CDCl_3$) δ 3.83 (t, $J$ = 5.3 Hz, 2H), 3.53 (d, 2H), 3.03 (t, $J$ = 5.7 Hz, 2H), 2.29 (t, 1H), 1.83 – 1.76 (m, 2H).

$^{13}C$ NMR (125 MHz, $CDCl_3$) δ 80.10, 72.83, 63.47, 47.89, 37.70, 30.18.

HRMS (ESI): $C_6H_{11}NO$; Calculated mass, $[M+H]^+$ 114.0919; Observed mass, $[M+H]^+$114.0914.

**Scheme S8.** Synthetic scheme for **11**.

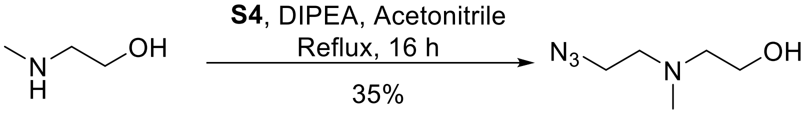


**11**

**2-((2-azidoethyl)(methyl)amino)ethan-1-ol (11)**

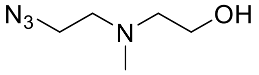

**11**

A round-bottomed flask containing azido ethanolamine (1.63 g, 18.78 mmol, 1 eq.) was subjected to high vacuum, flushed with nitrogen gas and to this flask, dry tetrahydrofuran (80 mL) and dry triethylamine (7.85 mL, 56.33 mmol, 3 eq.) were added. The flask was cooled to 0 °C and methanesulfonyl chloride (4.36 mL, 56.33 mmol, 3 eq.) was added dropwise under nitrogen atmosphere. The resulting reaction mixture was stirred for 2 h at 0 °C and then washed with 1 N NaOH (1 × 60 mL) and ethyl acetate (2 × 30 mL). The organic layer was dried by adding anhydrous sodium sulfate and concentrated under reduced pressure. This concentrated mesylate (**S4**) was added to a solution of *N*-methyl ethanolamine (750 μL, 9.38 mmol, 1 eq.) and *N,N*-diisopropylethylamine (1.96 mL, 11.26 mmol, 1.2 eq.) in acetonitrile (40 mL), and the resultant solution was stirred for 16 h at 80 °C. Subsequently, the solvent was evaporated under reduced pressure, and washed with 1N NaOH solution (1 × 50 mL) and ethyl acetate (3 × 30 mL). The organic layers were pooled, dried by adding anhydrous sodium sulfate, concentrated and subjected to silica gel column chromatography (1% triethylamine in ethyl acetate) to afford **11** as a yellow oil (467 mg, 35%).

$^1$H NMR (400MHz, $CDCl_3$): δ 3.61 (t, J = 5.27 Hz, 2H), 3.36 (t, J = 5.95 Hz, 2H), 2.68 (t, J = 5.97 Hz, 2H), 2.60 (t, J = 5.27 Hz, 2H), 2.32 (s, 3H).

$^{13}$C NMR (125 MHz, $CDCl_3$) δ 59.23, 58.71, 56.79, 49.07, 41.75.

HRMS (ESI): $C_5H_{12}N_4O^+$; Calculated mass, $[M+H]^+$ 145.1089 Observed mass, $[M+H]^+$ 145.1084.

**Scheme S9.** Synthetic scheme for **14**.

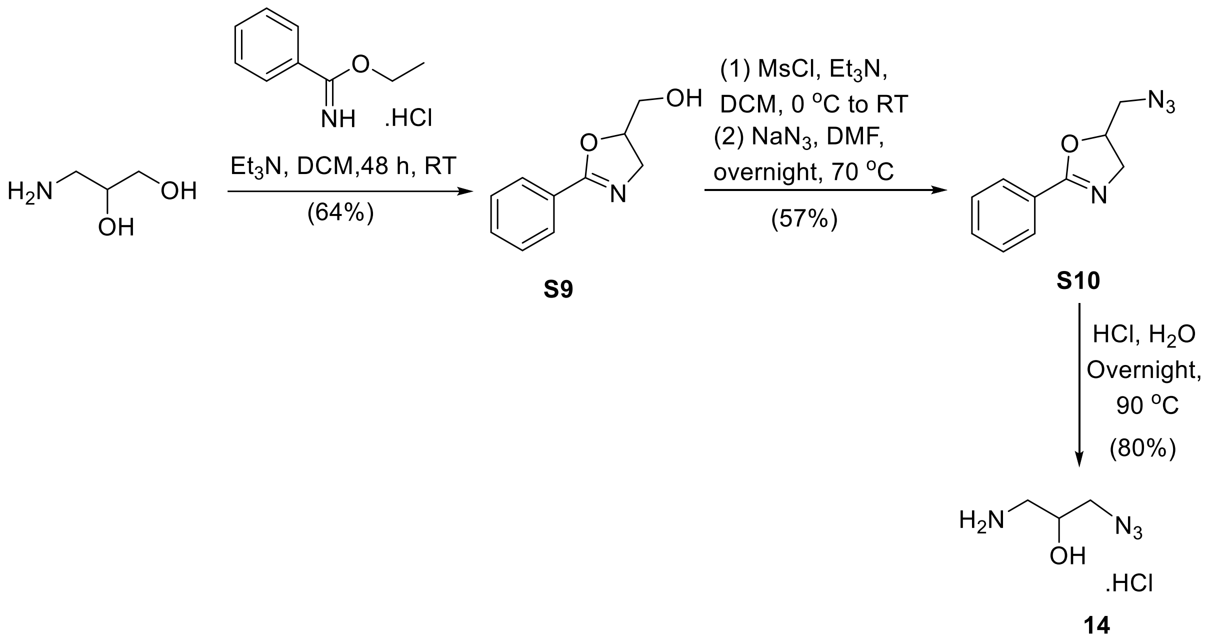


**(2-phenyl-4,5-dihydrooxazol-5-yl)methanol (S9)**

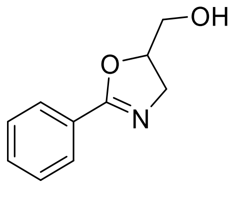


**S9**

A round-bottomed flask charged with ethyl benzimidate hydrochloride (500 mg, 2.69 mmol, 1.0 eq.) was subjected to high-vacuum and then purged with nitrogen. Subsequently, anhydrous dichloromethane (7 mL) was added, and to the resultant turbid white solution, dry triethylamine (432 μL, 3.10 mmol, 1.15 eq.) was added drop-wise and the mixture was stirred for 30 min at room temperature. Subsequently, 3-amino-1,2-propanediol (306 mg, 3.36 mmol, 1.25 eq.) was added and the resultant reaction mixture was stirred for 48 h at room temperature. The solvent was removed under reduced pressure and the residue was subjected to silica gel column chromatography (ethyl acetate) to obtain **S9** as a colourless oil (384 mg, 64%).

$^1H$ NMR (500 MHz, $CDCl_3$): δ 7.94 (d, J = 7.32 Hz, 2H), 7.48 (t, J = 7.38 Hz, 1H), 7.41 (t, J = 7.56 Hz, 2H), 4.85-4.80 (m, 1H), 4.10 (dd, J = 14.71, 9.93 Hz, 1H), 3.86- 3.81 (m, 2H), 3.74 (d, J = 4.76 Hz, 1H), 2.09 (s, 1H).

$^{13}C$ NMR (125 MHz, $CDCl_3$): δ 164.17, 131.58, 128.48, 128.29, 127.59, 80.28, 64.25, 56.42.

HRMS (ESI): $C_{10}H_{11}NO_2$; Calculated mass, $[M+H]^+$ 178.0863; Observed mass, $[M+H]^+$ 178.0858.

**5-(azidomethyl)-2-phenyl-4,5-dihydrooxazole (S10)**

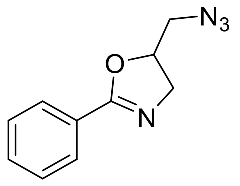


**S10**

A round-bottomed flask was charged with phenyl oxazole methanol (2.0 g, 11.28 mmol, 1.0 eq.), subjected to high-vacuum and then purged with nitrogen and dissolved in anhydrous dichloromethane (79 mL), followed by addition of triethylamine (3.0 mL, 22.57 mmol, 2.0 eq.) and resultant mixture was cooled to 0 °C, and methane sulfonyl chloride (1.75 mL, 22.57 mmol, 2 eq.) was added. The resulting mixture was stirred for 3 h at room temperature. The reaction was quenched by adding a 0.1 M HCl solution (23 mL) and resulting solution was washed with dichloromethane (2 × 30 mL). The organic phases were combined and washed with saturated sodium hydrogen carbonate solution (23 mL). The organic layers were combined, dried by adding anhydrous sodium sulphate and concentrated under reduced pressure. The resulting crude residue was used in the next step of synthesis. A round bottomed flask charged with the crude mesylate of phenyl oxazole ethanol was dissolved in *N,N*- Dimethylformamide (40 mL) and added sodium azide (3.66 g, 56.43 mmol, 5.0 eq.) and the mixture was stirred at 70 °C for 24 h. After removing *N,N*-Dimethylformamide under reduced pressure, the residue was diluted with water (1 × 160 mL) and the resulting water layer was washed with diethyl ether (8 × 50 mL). The organic layers were combined and dried by adding anhydrous sodium sulphate, concentrated and subjected to silica gel column chromatography (33% ethyl acetate in hexane) to afford colourless oil **S10** (1.32 g, 57%).

$^1H$ NMR (500 MHz, $CDCl_3$): δ 7.95 (d, J = 7.54 Hz, 2H), 7.49 (t, J = 7.37 Hz, 1H), 7.42 (t, J = 7.62 Hz, 2H), 4.92-4.87 (m, 1H), 4.15 (dd, J = 14.91, 9.83 Hz, 1H), 3.82 (dd, J = 14.92, 6.91 Hz, 1H), 3.49-3.43 (m, 2H).

$^{13}C$ NMR (125 MHz, $CDCl_3$): δ 163.88, 131.70, 128.56, 128.35, 127.37, 78.35, 57.87, 54.13.

HRMS (ESI): $C_{10}H_{10}N_4O$; Calculated mass, $[M+H]^+$ 203.0927; Observed mass, $[M+H]^+$ 203.0912.

**1-amino-3-azidopropan-2-ol hydrogen chloride (14)**

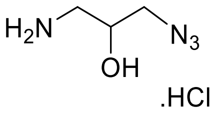


**14**

To a round-bottomed flask containing **S10** (100 mg, 0.5 mmol, 1eq.) was dissolved in 4 N HCl in water (5 mL). This reaction mixture stirred overnight at 90 °C. Subsequently, the reaction mixture was allowed to cool to room temperature and then washed with dichloromethane (5 × 20 mL). The water layer was concentrated under reduced pressure to yield **14** as a colourless oil (61 mg, 80%).

$^1$H NMR (500 MHz, DMSO-$d_6$): δ 8.15 (s, 3H), 5.90 (bs, 1H), 3.95-3.80 (m, 1H), 3.38 (dd, J = 12.78, 4.13 Hz, 1H), 3.29 (dd, J = 12.78, 6.21 Hz, 1H), 2.89-2.85 (m, 1H), 2.75-2.71 (m, 1H). $^{13}$C NMR (125 MHz, DMSO-$d_6$): δ 66.69, 53.56, 41.88.

HRMS (ESI): $C_3H_9N_4O^+$; Calculated mass, $[M+H]^+$ 117.0771; Observed mass, $[M+H]^+$ 117.0783.

**Scheme S10.** Synthetic scheme for **17**.

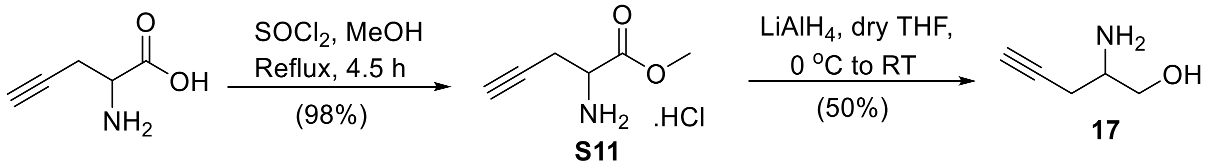


**Methyl 2-aminopent-4-ynoate hydrogen chloride (S11)**

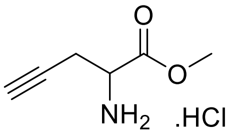


**S11**

In a round-bottomed flask, DL-propargyl glycine (390 mg, 3.45 mmol, 1 eq.) was dissolved in anhydrous methanol (6.5 mL) and cooled on an ice bath. Thionyl chloride (2.5 mL, 34.5 mmol, 10 eq.) was added drop-wise to the above solution under nitrogen atmosphere. This ice cooled reaction mixture was then allowed to attain room temperature and refluxed for 4.5 h under nitrogen atmosphere. After concentrating the reaction mixture under reduced pressure, the residue was subjected to silica gel column chromatography (10% methanol in dichloromethane) to obtain a pale white solid **S11** (552 mg, 98%).

$^1$H NMR (500 MHz, $D_2O$): δ 4.41 (t, J = 5.4 Hz, 1H), 3.90 (s, 3H), 3.06-2.94 (m, 2H), 2.61 (t, J = 2.6 Hz, 1H).

$^{13}$C NMR (125 MHz, DMSO) δ 171.13, 78.94, 75.26, 53.30, 52.00, 22.13.

HRMS (ESI): $C_6H_9NO_2^+$; Calculated mass, $[M]^+$ 128.0706; Observed mass, $[M]^+$ 128.0714.

**2-aminopent-4-yn-1-ol (17)**

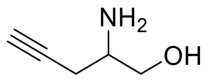

**17**

To a two-neck round-bottomed flask flushed with nitrogen gas was added **S11** (550 mg, 4.32 mmol, 1 eq.) and dry tetrahydrofuran (70 mL), forming a suspension that was stirred on an ice bath for 10 min. Then, $LiAlH_4$ (345 mg, 9.08 mmol, 2.1 eq.) was added in small portions under nitrogen at 0 °C. This resultant mixture was allowed to stir at 0 °C for 10 min and then at room temperature for 6.5 h. After stirring was completed, the reaction mixture was cooled on an ice bath and unreacted $LiAlH_4$ was quenched by adding a saturated solution of potassium sodium tartarate (32 mL). The resultant reaction mixture was allowed to stir at room temperature for 8 h and then washed with ethyl acetate (5 × 100 mL). The organic phases were combined and solvent was removed under reduced pressure. The obtained residue was subjected to silica gel column chromatography (10% methanol in chloroform) to obtain beige coloured solid **17** (167 mg, 50%).

$^1$H NMR (500 MHz, $D_2O$): δ 3.51 (ddd, *J* = 39.6, 11.3, 5.9 Hz, 2H), 2.95 (p, *J* = 5.9 Hz, 1H), 2.39 – 2.23 (m, 2H).

$^{13}$C NMR (125 MHz, DMSO) δ 82.46, 72.22, 65.05, 51.96, 23.45.

HRMS (ESI): $C_5H_9NO^+$; Calculated mass $[M+H]^+$ 100.0762, Observed mass, $[M+H]^+$ 100.0716.

**Scheme S11.** Synthetic scheme for **19**.

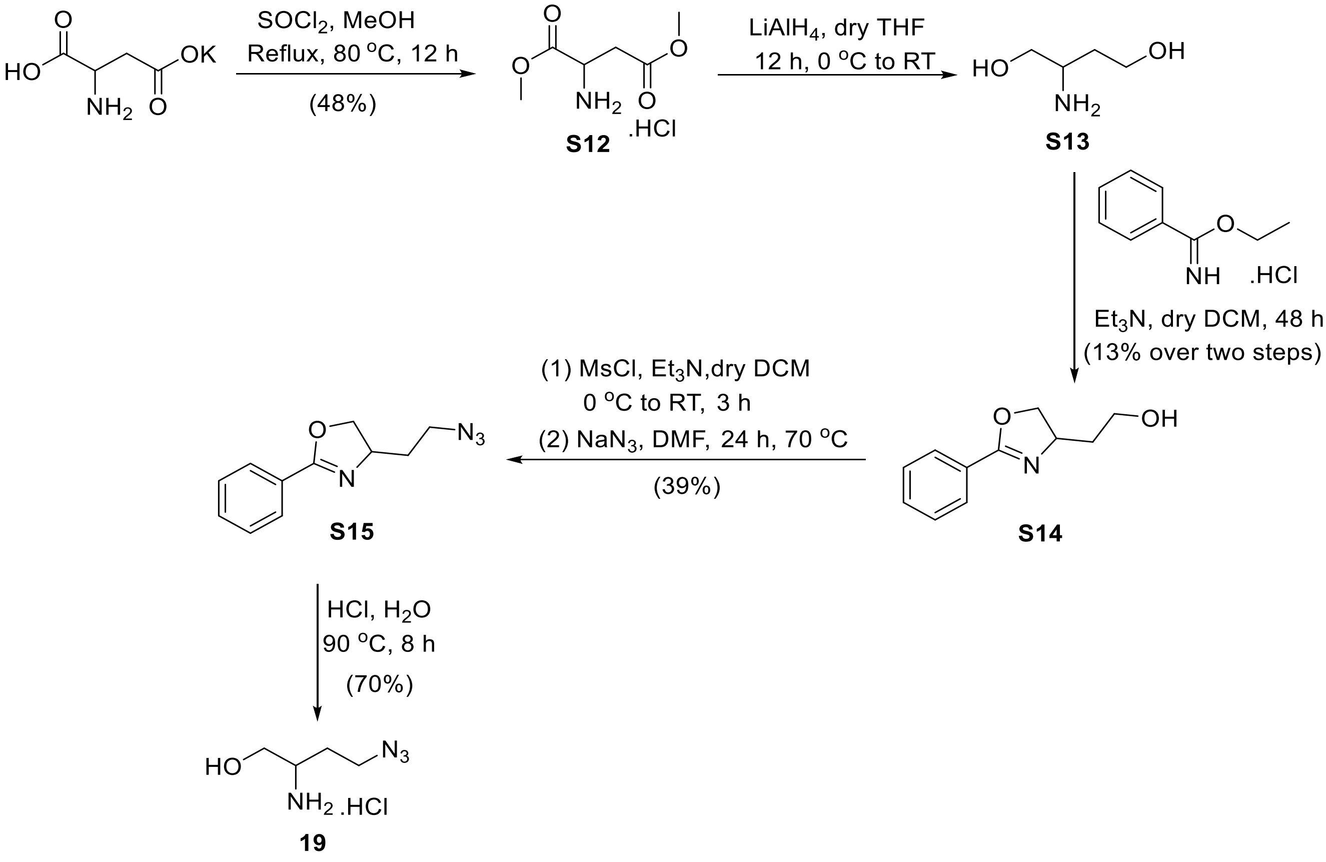

**Dimethyl aspartate hydrogen chloride (S12)**

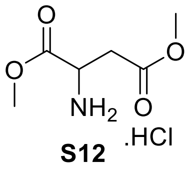

**S12**

The synthesis was performed by following a reported protocol with minor changes[2]. L-aspartic acid potassium salt (4.0 g, 23.4 mmol, 1 eq.) was added to a round-bottomed flask and dissolved in anhydrous methanol (44 mL). The resultant solution was cooled on an ice bath and thionyl chloride (17 mL, 234.0 mmol. 10 eq.) was added drop-wise. The reaction mixture was then allowed to attain room temperature and refluxed overnight. Subsequently, the solvent was removed under reduced pressure and the residue was subjected to silica gel column chromatography (10% methanol in dichloromethane) to obtain a pale yellow solid **S12** (2.15 g, 48%).

$^1$H NMR (500 MHz, DMSO-$d_6$): δ 8.79 (s, 2H), 4.32 (s, 1H), 3.72 (s, 3H), 3.64 (s, 3H), 3.08-2.97 (m, 2H).

$^{13}$C NMR (125 MHz, DMSO-$d_6$): δ 169.55, 168.65, 53.03, 52.15, 48.39, 33.97.

HRMS (ESI): $C_6H_{12}NO_4^+$; Calculated mass, $[M+H]^+$ 162.0761; Observed mass, $[M+H]^+$ 162.0758.

**2-aminobutane-1,4-diol (S13)**

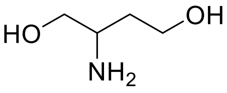

**S13**

The synthesis was performed by following a reported protocol with minor changes[2]. A two-neck round-bottomed flask was charged with **S12** (3.0 g, 18.61 mmol, 1 eq.) and anhydrous tetrahydrofuran (100 mL) was added, forming a turbid solution which was cooled to 0 ºC. $LiAlH_4$ (2.97 g, 78.18 mmol, 4.2 eq.) was added in small portions under nitrogen at 0 °C and the resultant mixture was allowed to stir at 0 °C for 30 min. Subsequently, the mixture was allowed to attain room temperature and stirred for 12 h. Then, the reaction mixture was cooled on an ice bath and unreacted $LiAlH_4$ was quenched by adding deionised water (5 mL), 1 M NaOH (2.4 mL) and allowed to stir at room temperature for 1 h. The water was removed by adding anhydrous sodium sulphate, and the resultant mixture was subjected to vacuum filtration. The solvent in the filtrate was removed under reduced pressure to yield compound **S13** in a crude form as a pale-yellow viscous liquid (716 mg). This crude **S13** was used for the next step without further purification.

**2-(2-phenyl-4,5-dihydrooxazol-4-yl)ethan-1-ol (S14)**

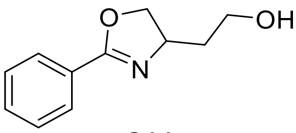

**S14**

A flame dried two-neck round-bottomed flask charged with ethyl benzimidate hydrochloride (636 mg, 3.42 mmol, 1.0 eq.) was subjected to high-vacuum and then purged with nitrogen. Subsequently, dry dichloromethane (9 mL) was added to form a turbid white solution followed by slow addition of dry triethylamine (503 µL, 3.94 mmol, 1.15 eq.) and resulting mixture was stirred for 30 min. at room temperature. Subsequently, 2-amino-1,4-butanediol (450 mg, 4.28 mmol, 1.25 eq.) was added and the resultant reaction mixture was stirred for 48 h at room temperature. After removing the solvent under reduced pressure, the residue was subjected to silica gel column chromatography (50% ethyl acetate in hexane to pure ethyl acetate) to obtain colourless oil **S14** (387 mg, 13%).

$^{1}$H NMR (500 MHz, $CDCl_3$): δ 7.92 (d, J = 7.5 Hz, 2H), 7.49 (t, J = 7.4 Hz, 1H), 7.41 (t, J = 7.6 Hz, 2H), 4.59 (t, J = 8.9 Hz, 1H), 4.44 (qd, J = 9.1, 5.0 Hz, 1H), 4.05 (t, J = 8.3 Hz, 1H), 4.00-3.93 (m, 1H), 3.91-3.85 (m, 1H), 2.84 (br, s, 1H), 1.97-1.82 (m, 2H).

$^{13}$C NMR (125 MHz, $CDCl_3$) δ 164.14, 131.75, 128.51, 128.45, 127.40, 73.25, 66.60, 61.97, 38.33.

HRMS (ESI): $C_{11}H_{13}NO_2$. Calculated mass, $[M+H]^+$ 192.1019; Observed mass, $[M+H]^+$ 192.1005.

**4-(2-azidoethyl)-2-phenyl-4,5-dihydrooxazole (S15**)

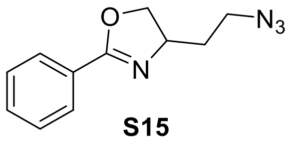


**S15**

A two-neck round-bottomed flask charged with **S14** (836 mg, 4.37 mmol, 1.0 eq.) was subjected to high-vacuum and then purged with nitrogen. Anhydrous dichloromethane (30 mL) was then added to dissolve the compound. To this flask, anhydrous triethylamine (1.22 mL, 8.74 mmol, 2.0 eq.) was added drop-wise via a syringe and the resultant mixture was cooled to 0 °C. After 10 min, methane sulfonyl chloride (677 µL, 8.74 mmol, 2 eq.) was added drop-wise at 0 °C. The resulting mixture was allowed to attain room temperature and stirred for 4 h. The reaction was quenched by adding a 0.1 M HCl solution (9 mL) and the resulting solution was washed with dichloromethane (3 × 30 mL). The organic phases were combined and washed with saturated sodium hydrogen carbonate solution (1 × 5 mL), dried by adding anhydrous sodium sulphate, and concentrated under reduced pressure. A round-bottomed flask (50 mL) charged with this crude mesylate of phenyl oxazole ethanol was dissolved in *N,N*-dimethylformamide (15 mL), and sodium azide (1.42 g, 21.85 mmol, 5.0 eq.) was added. The resultant solution was stirred at 70 °C for 24 h. Subsequently, *N,N*-dimethyl formamide was removed under reduced pressure, the residue was diluted with water (65 mL) and washed with diethyl ether (6 × 30 mL). After removing the organic solvent under reduced pressure, the residue was subjected to silica gel column chromatography (20% ethyl acetate in hexane) to obtain colourless oil **S15** (371 mg, 39%).

$^{1}$H NMR (500 MHz, $CDCl_3$): δ 7.94 (d, J = 8.0 Hz, 2H), 7.48 (t, J = 7.4 Hz, 1H), 7.41 (t, J = 7.6 Hz, 2H), 4.54 (t, J = 8.9 Hz, 1H), 4.43-4.34 (m, 1H), 4.07 (t, J = 8.0 Hz, 1H), 3.55 (t, J = 6.9 Hz, 2H), 1.98-1.85 (m, 2H).

$^{13}$C NMR (125 MHz, $CDCl_3$): δ 164.21, 131.59, 128.48, 128.42, 127.70, 72.58, 64.37, 48.76, 35.29.

HRMS (ESI): $C_{11}H_{12}N_4O$; Calculated mass, $[M+H]^+$ 217.1084; Observed mass, $[M+H]^+$ 217.1080.

**2-amino-4-azidobutan-1-ol hydrogen chloride (19)**

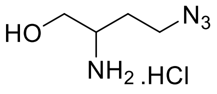

**19**

Oxazoline ring hydrolysis was performed as reported previously[3]. **S15** (371 mg, 1.71 mmol, 1eq.) was dissolved in 4 N HCl in water (17.5 mL, 10 mL per mmol 19) and then heated at 90 °C for 8 h. Subsequently, the reaction mixture was allowed to cool to room temperature and washed with dichloromethane (10 × 30 mL) and diethyl ether (10 x 30 mL). The resulting water layer was washed with ethyl acetate (5 × 30 mL) followed by dichloromethane (5 × 30 mL) and then concentrated under reduced pressure to afford **19** as a brown oil (200 mg, 70%).

$^1$H NMR (500 MHz, DMSO-$d_6$): δ 8.11 (s, 3H), 5.35 (s, 1H), 3.61-3.58 (m, 1H), 3.50 (m, 3H), 3.12 (t, J = 4.77 Hz, 1H), 1.80 (q, J = 6.94 Hz, 2H).

$^{13}$C NMR (125 MHz, DMSO-$d_6$): δ 60.40, 50.16, 46.91, 28.31.

HRMS (ESI): $C_4H_{11}N_4O^+$; Calculated mass, $[M+H]^+$ 131.0927; Observed mass, $[M+H]^+$ 131.0931.

### IV. Antibodies used in the study

**Table S1:** Antibodies used for western blot analysis

| Protein name | Manufacturer | Catalog no. | Dilution used |
|---|---|---|---|
| COX IV | Cell signalling technology | 4850 | 1:7000 |
| VDAC | Abcam | ab154856 | 1:4000 |
| GM130 | Abcam | ab52649 | 1:5000 |
| Na+/K+ ATPase | Abcam | ab76020 | 1:25000 |
| Calnexin | Cell signalling technology | C5C9 | 1:2000 |
| β-actin | Sigma | A2066 | 1:10000 |
| Nup62 | Abcam | ab96134 | 1:4000 |
| VPS13A | Merck | HPA021662 | 1:1000 |
| Goat anti-rabbit IgG HRP | Abcam | ab205718 | 1:5000 |

### V. Lipidomics procedures

**Compound administration and lipid extraction from HEK293/HeLa cells:** For all lipidomics experiments, 3 million cells in 10 mL of complete medium were seeded into one 10 cm sterile culture dishes (for each treatment) and incubated for 24 h in a $CO_2$ incubator. The compounds were administered as per the desired final concentrations from 65 mM stock solutions in PBS and metabolic labelling was allowed to occur over the desired time period in a $CO_2$ incubator. To extract lipids from labelled cells, the adhered cells were washed with ice-cold PBS (2 × 5 mL), resuspended in ice-cold PBS (5 mL), pooled in a centrifuge tube and centrifuged at 1,000 g for 5 min at room temperature. The supernatant was then aspirated off and the lipids were extracted from the resultant pellet using modified Bligh Dyer method. Briefly, the cell pellet was treated with a mixture of chloroform and methanol (1 mL of a 1:2 *v*/*v* solution), vortexed for 1 min followed by addition of chloroform (0.5 mL) and 1 M NaCl (0.5 mL), and again vortexed for 1 min. Phase separation was achieved by centrifuging the mixture at 3,300 g at room temperature for 30 min. The lower lipid-enriched organic layer was separated and dried using a benchtop vacuum concentrator (Eppendorf concentrator plus) at room temperature. For compound parameter optimization via direct infusion method, the dried lipid samples were redissolved in chloroform (0.5 mL) and 360 µL of this lipid stock solution was added to a mixture of chloroform/methanol/300 mM ammonium acetate in water such that the final composition was 360/798/42 (*v*/*v*/*v*) and the final volume of the lipid mixture was 1.2 mL. For all LC-MS/MS analysis, the dried lipids were dissolved in a mixture of methanol and chloroform (100 µL of a 2:1 *v*/*v* solution). For all control experiments, the procedure described above was followed without administration of any ethanolamine derivatives.

**HPLC method details:**

Lipids extracted from cells were fractionated on a 2.6 µm Kinetex HILIC column (I.D. 100 × 2.1 mm, 100 Å, Phenomenex). The elution protocol used is summarized below:

Mobile Phase A: 95:5 Acetonitrile: Water (containing 10 mM ammonium acetate)

Mobile Phase B: 1:1 Acetonitrile: Water (containing 10 mM ammonium acetate)

Mobile Phase C: 97:3 Acetonitrile: Water (containing 10 mM ammonium acetate)

Gradient:

0 to 1 min: 100% C

1 to 20 min: 100% C to 100% A

20 to 32 min: 100% A to 30% B and 70% A

32 to 39 min: 30% B and 70% A to 100% A

39 min to 40 min: 100% A to 100% C

40 min to 54 min: 100% C (for column re-equilibration)

Flow rate: 0.5 mL/min

Column oven temperature: 40 °C

Sample volume: 10 µL of lipid solution extracted from cells.

**Mass spectrometry details:** Lipids eluted from the HILIC column were analyzed and detected on a Sciex 4500 QTRAP mass spectrometer equipped with an ESI source heated to 400 °C. The scan speed was maintained at 200 Da/sec. The electrospray capillary was held at +5500 volts, curtain gas was set at 35 (arbitrary units), CAD gas was maintained at the medium level and the two ion source gases GS1 and GS2 were set at 50 and 60 (arbitrary units) respectively.

**Scan modes:** The neutral loss (NL) mode was used for detecting all the non-natural or native ethanolamine lipids, as well as all methylated versions of the labelled lipids except the ones that are quaternary amines (PC lipid derivatives). To detect the PC derivatives of ethanolamine analogs, precursor ion scan (PIS) scan was performed. Multiple reaction monitoring (MRM) scans were performed to quantitate the major lipid species in D4-ethanolamine and compound **19** administered cells. All MRM transitions employed in this study are depicted below in **Table S2**.

**Table S2: MRM Q1/Q3 for lipidomics experiments performed in this study.**

| **Lipid Species** | **Q1 mass (Da)** | **Q3 mass (Da)** |
|---|---|---|
| PE standard (37:4) | 754.54 | 613.52 |
| 34:1 D4-Eth labelled PE species | 722.56 | 577.52 |
| 36:2D4-Eth labelled PE species | 748.58 | 603.53 |
| 32:1 D4-Eth labelled PE species | 694.53 | 549.49 |
| 38:5 D4-Eth labelled PE species | 770.60 | 625.36 |
| 36:4 D4-Eth labelled PE species | 744.60 | 599.56 |
| 34:8 D4-Eth labelled PE species | 708.60 | 563.56 |
| 34:2 D4-Eth labelled PE species | 720.60 | 575.56 |
| 34:1 **19**-labelled PE species | 787.57 | 577.52 |
| 36:2 **19**-labelled PE species | 813.58 | 603.53 |
| 32:1 **19-** labelled PE species | 759.54 | 549.49 |

**Mitochondrial lipidomics**: A total of 80 µg of mitochondrial protein (refer Section X) was subjected to lipid extraction. Prior to extraction, 50 pmol of IS PE (17:0/20:4) was spiked into each sample to enable normalization. Lipids were extracted using the Bligh–Dyer method following protocol described above. Extracted lipid samples were analyzed using MRM scanning. Quantification was performed by calculating the area under the curve for each lipid species. The obtained values were first normalized to the spiked PE lipid standard to account for extraction efficiency. To further correct for differences in mitochondrial loading, the normalized lipid intensities were subsequently adjusted using VDAC protein levels determined by densitometric analysis from western blotting, which served as a marker for mitochondrial abundance. Following normalization, data were represented as bar graphs. Statistical significance was determined using multiple unpaired t-tests.

**Whole cell lipidomics procedure employed in VPS13A experiments:** Following VPS13A knockdown, lipids were extracted from the cells. Briefly, adherent cells were washed twice with 5 mL of ice-cold PBS, harvested in 5 mL of ice-cold PBS, transferred to a centrifuge tube, and pelleted by centrifugation at 1,000 × g for 5 min at room temperature. The supernatant was carefully removed, and an aliquot was collected for protein quantification using the

Bradford assay. The remaining cell suspension was centrifuged again, and 50 pmol of the internal standard PE (17:0/20:4) was added to the resulting cell pellet. Lipid extraction was then performed using the Bligh–Dyer method as described in Section V. For lipid quantification, MRM scans were performed and the area under the curve for each lipid species was first normalized to the internal standard PE and subsequently normalized to the corresponding protein concentration. Statistical significance was determined using multiple unpaired t-tests.

**Compound parameters:** The compound parameters (declustering potential, entrance potential, collision energy, and collision cell exit potential) were optimized by ramping each of them against the intensity by the direct infusion method. Native ethanolamine lipids were detected by employing a NL of 141 Da (*m/z* for phosphoethanolamine) and the compound parameters were optimized separately for each ethanolamine analogs.

**Lipidomics data processing using the LipidView software:** To identify ethanolamine lipids labelled with D4 and non-natural headgroups, the LipidView software (Sciex, version 1.2) was utilized to process the LC/MS data. This involved manual modifications to the software's database. Peaks were adjusted for isotopic overlap and subjected to processing with a mass tolerance of 0.5 Da, a minimum intensity threshold of 0.1%, and a minimum signal-to-noise ratio of 10. Subsequently, the phospholipid species obtained were validated against the Lipid Maps structure database (LMSD). Species not present in the LMSD were excluded from further analysis.

### VI. Imaging procedures

**Fixed cell imaging:** Glass-bottom dishes (35 mm) were coated with 200 µl of 0.001% poly-L-lysine for 15 min to promote cell adhesion. Dishes were subsequently washed three times with 200 µl of 1X Phosphate-Buffered Saline (PBS). HeLa cells were seeded at a density of 0.2 × $10^6$ cells/mL in 1 mL of complete medium and incubated for 12 h at 37°C in a humidified atmosphere containing 5% $CO_2$. After a 12 h incubation in a $CO_2$ incubator at 37 °C, 31.7 µL of 65 mM stock solutions of ethanolamine analogs in PBS was administered into each well (Final concentration: 2 mM). The same volume of PBS devoid of ethanolamine analogs was added in the control wells. After a further 24 h incubation in a $CO_2$ incubator at 37 °C, cells were washed with PBS (3 × 500 µL) at room temperature and fixed by incubating them in formaldehyde (200 µL of a 3.7% *w*/*v* solution in PBS) for 30 min at 37 °C. The fixed cells were washed with TBS (3 × 500 µL) and then incubated in a click reaction cocktail (100 µL) for 30 min at room temperature under dark. The click reaction cocktail was made in 0.1 M Tris-HCl buffer at pH 8.5 and contained 1 mM $CuSO_4.5H_2O$ (added from a stock of 500 mM in water), 100 mM L-ascorbic acid (added from a stock of 500 mM in water), and 20 µM 5-azidofluorescein (added from a stock of 10 mM in DMSO) for imaging alkynyl ethanolamine analogs or 20 µM alkynyl rhodamine (added from a stock of 10 mM in DMSO) for imaging cells treated with azido ethanolamine analogs. After completion of this 30 mins long incubation, the cells were washed with TBS (3 × 500 µL), 0.5 M NaCl (3 × 500 µL), and then again with TBS (3 × 500 µL). Finally, 250 µL TBS was added to each well, and the cells were imaged using the Olympus FV3000 confocal imaging system. All images were acquired by using a 100XO objective either under the 488 nm channel (Laser: 1.0, HV: 600) for analogs **2**, **4**, **5**, **8**, and **12** or the 561 nm channel (Laser: 0.1, HV: 500) for analogs **3**, **7**, **14**, **19**, and the representative images are provided in **Figure 1E**. Each experiment was performed three times by administering the cells with the aforementioned ethanolamine analogs into three separate wells and performing the imaging experiment as described above.

**Live-cell imaging:**

**Whole cell 19-PE imaging with BCN-BODIPY:** Glass-bottom dishes (35 mm) were coated with 200 μl of 0.001% poly-L-lysine for 15 min to promote cell adhesion. Dishes were subsequently washed three times with 200 μl of 1X Phosphate-Buffered Saline (PBS). HeLa cells were seeded at a density of 0.2 × $10^6$ cells/mL in 1 mL of complete medium and incubated for 12 h at 37°C in a humidified atmosphere containing 5% $CO_2$. Following the initial incubation, the medium was replaced with 1 mL of complete medium containing the desired concentration of compound **19** (diluted from a 1 mM stock of **19** in PBS for 10/100 μM administration experiments, and from a 65 mM stock for 200 μM). Cells were incubated for the desired labelling duration in 5% $CO_2$ incubator at 37°C. Subsequently, the cells were washed three times with 1 ml of complete medium, and the medium was replaced with 100 μL of 1 μM BCN-BODIPY in serum-free medium for whole cell imaging experiments. Cells were incubated for 15 min at 37°C to allow for strain-promoted click chemistry, and then washed three times with 500 μL of complete medium. An additional 500 μL of complete medium was added, and the cells were incubated for 30 min at 37°C in the $CO_2$ incubator to remove remaining BCN-BODIPY dye. Before imaging, the medium was replaced with 500 μL of imaging media (DMEM without phenol red, 10% FBS, 1% penicillin-streptomycin). Fluorescence imaging was performed using an Alexa Fluor 488 filter setting. Imaging parameters employed for various experiments are depicted in **Table S3**.

**Organellar PE imaging experiments (colocalization experiments):** Glass bottom dishes (35 mm) were seeded and administered with compound **19** and labelled with BCN-BODIPY as described above. Cells were stained with 100 μL of 1 μM ER tracker/50 nM lysotracker/1 μM Golgi tracker/100 nM mitotracker markers in serum-free medium for 30 min. Following three washes with 500 μL complete medium, an additional 500 μL of imaging medium was added, and the cells were imaged by employing the settings listed in **Table S3**. Colocalization analysis was done using ImageJ. For Pearson's correlation coefficient (PCC) analysis, Z-stacks were acquired spanning the entire cell volume (black to black). PCC values were calculated using the BIOP JACoP plugin in ImageJ. Huang's auto-thresholding method was applied to both channels for PCC analysis in the case of ER colocalization, whereas Otsu auto-thresholding was used for background subtraction in both channels for lysosomal and mitochondrial colocalization analyses. PCC values are presented as mean ± SD.

**Imaging PE externalization during apoptosis:** Glass-bottom dishes (35 mm) were first coated with 200 μL of 0.001% poly-L-lysine for 15 min to enhance cell adherence, followed by washing with 1× PBS (3 × 200 μL). HeLa cells were seeded at a density of 0.2 × $10^6$ cells/mL in 1 mL of complete medium and incubated for 12 h at 37 °C in a humidified atmosphere containing 5% $CO_2$. The cells were then treated with **19** (200 μM in complete medium) and incubated for 24 h under the same conditions. Following incubation, cells were washed with complete medium (3 × 1 mL) to remove excess compound. Apoptosis was induced to promote externalization of **19**-PE by incubating the cells with 200 μL of 1X PBS containing 6 μM calcium ionophore (A23187[4]) and 2 mM $CaCl_2$ for 45 min at 37 °C. After induction, cells were washed with complete medium (3 × 1 mL). For bioorthogonal labelling of externalized lipids, cells were incubated with 100 μL of 40 μM AF647-DBCO (prepared in serum-free DMEM) for 90 min at

4 °C to minimize endocytosis and preserve membrane surface localization. The cells were then washed with complete medium (3 × 0.5 mL) and further incubated in 0.5 mL complete medium at 4 °C for an additional 30 min. To validate PE externalization, cells were incubated with 100 µL of 45 µM duramycin-LC-biotin–streptavidin AF594 conjugate for 30 min at room temperature, followed by washing with complete medium (3 × 0.5 mL). Prior to imaging, cells were maintained in imaging medium and visualized using an Olympus FV3000 confocal microscope equipped with 594 nm and 647 nm laser lines and a 100X oil immersion objective. The cells were imaged by employing the settings listed in **Table S3**. Co-localization of AF647 (compound **19**-PE) and AF594 (duramycin) signals on the outer leaflet of the plasma membrane was analyzed, and line intensity profiles were generated using ImageJ. Line intensity profiles for apoptosis analysis were generated using the "Plot Profile" tool in the ImageJ analysis window.

**19-PE imaging in mitochondria using Cy5-DBCO:** Glass-bottom dishes (35 mm) were seeded as described above and after 12 h, the cells were treated with compound **19** (200 µM in 1 mL complete medium) and incubated for an additional 24 h under the same conditions. After incubation, cells were washed with complete medium (3 × 1 mL) to remove excess compound. For bioorthogonal labelling, cells were incubated with 100 µL of 200 nM Cy5-DBCO (prepared in serum-free DMEM) for 15 min at 37 °C. The cells were then washed with complete medium (3 × 0.5 mL) and subsequently treated with 400 nM Cy7-$N_3$ for 15 min at 37 °C to quench unreacted Cy5-DBCO. Subsequently, cells were washed again with complete medium (0.5 mL) for 15 min. Then, MT10 mitotracker (0.1 µM) was added and the cells were incubated in this dye for 30 min. Cells were then briefly incubated in 100 µL of wash medium (high [$K^+$] RPMI media containing 103.44 mM KCl, 5.33 mM NaCl, 50 µM CCCP, 20 µM valinomycin, and 10% FBS) for 1 min, after which the medium was removed and cells were washed with complete medium (3 × 0.5 mL) to ensure minimal background fluorescence. Prior to imaging, 250 µL of imaging medium was added to each dish. Fluorescence imaging was performed using an Alexa Fluor 640 laser line. The cells were imaged by employing the settings listed in **Table S3**. For imaging of VPS13A knockdown cells, cells were treated with 200 µM compound 19 for 24 h, followed by live-cell imaging using Cy5 DBCO following the above protocol.

**Microscope image acquisition settings used in this study:**

**Table S3: Microscope settings employed for different experiments**

| **Experiment** | **Dye** | **Organelle tracker** | **Laser (nm)** | **Laser power** | **HV value** | **Offset** |
|---|---|---|---|---|---|---|
| Figure 1 | Azidofluorescein | | 488 | 1 | 600 | 3 |
| | Rhodamine alkyne | - | 561 | 0.1 | 500 | 3 |
| Figure 2 | BCN-BODIPY | - | 488 | 0.1 | 600 | 3 |
| Figure 3 | AF647-DBCO | - | 640 | 3.0 | 600 | 3 |
| | Duramycin AF594 | PM | 561 | 2.0 | 600 | 3 |
| Figure 4 | BCN-BODIPY | - | 488 | 0.1 | 600 | 3 |
| | - | ER | 594 | 1.0 | 600 | 3 |
| | - | Golgi | 594 | 2.0 | 600 | 3 |
| | - | Lyso | 594 | 0.3 | 600 | 3 |
| | - | Mito | 594 | 1.2 | 600 | 3 |
| Figure 5 | Cy5-DBCO | - | 640 | 3.0 | 600 | 3 |
| | - | MT10 | 488 | 1.0 | 600 | 3 |

### VII. Procedure for TLC analysis followed by MS analysis on fluorescent spots

#### TLC analysis of fluorescently labelled 19-PE lipids (spot denoted by "*" in Figure 2B of the main text)

HeLa cells seeded at 0.3 million cells/mL in four 10 cm dishes were metabolically labelled with **19** (10 µM), and subjected to click chemistry with BCN-BODIPY (1 µM) for 15 min within a $CO_2$ incubator at 37 °C. Lipids were then extracted as per the protocol described in Section V and the dried lipid extracts were resuspended in 100 µL MeOH: $CHCl_3$ (2:1, *v*/*v*). Samples (5 µL each) were spotted onto silica gel 60 TLC plates using a Hamilton syringe, and TLC was performed using a solvent system composed of $CHCl_3$: MeOH: $H_2O$ (65:20:2, *v*/*v*/*v*). Fluorescent spots were visualized under 488 nm using a Bio-Rad ChemiDoc imaging system. The fluorescent spot corresponding to the **19**-labelled lipid conjugated to BCN-BODIPY was scraped from the TLC plate, resuspended in 500 µL LC/MS grade methanol, vortexed, and centrifuged at 8000g for 20 min. The supernatant was collected, passed through a syringe filter, and dried under vacuum in D-AL mode using a benchtop vacuum concentrator (Eppendorf Concentrator Plus) at room temperature.

For compound parameter optimisation, the same procedure was followed as mentioned in Section V. Mass spectrometry scans in the positive ion mode were performed for neutral loss (NL) of 20 Da (fragmentation depicted below in Figure S2) for the loss of HF using the following parameters: DP (58.77 V), EP (12.56 V), CE (42.10 V), and CXP (19.62 V) The LC conditions were identical to those employed for detecting **19**-PE lipids (Section V).

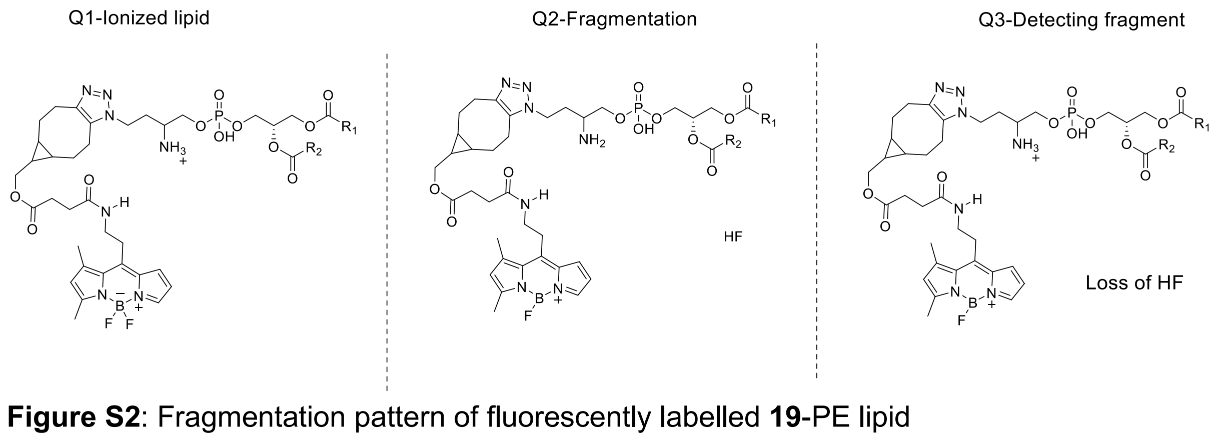


**Figure S2**: Fragmentation pattern of fluorescently labelled **19**-PE lipid

Expected masses for the three abundant **19**-PE lipid after click chemistry with BCN-BODIPY:

1. 34:1 **19-**labelled lipid conjugated to BCN-BODIPY

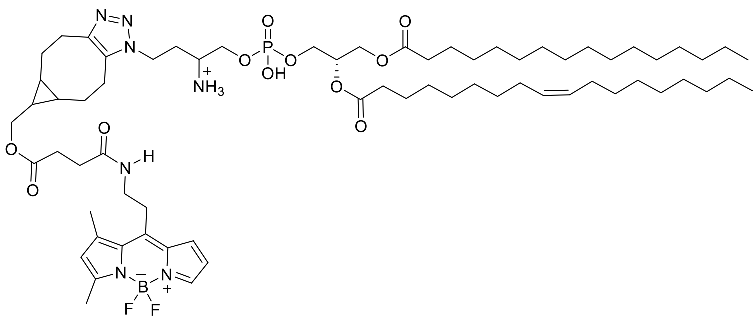


2. 36:2 **19**-labelled lipid conjugated to BCN-BODIPY

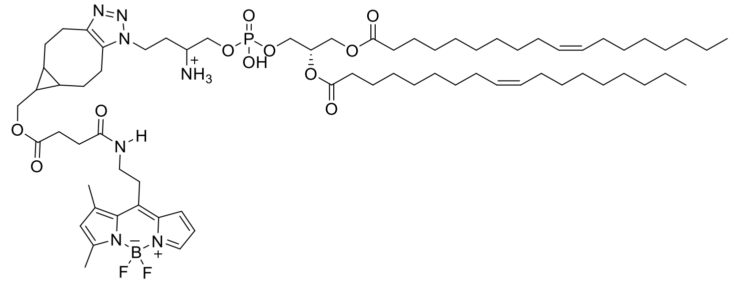


3. 32:1 **19-**labelled lipid conjugated to BCN-BODIPY

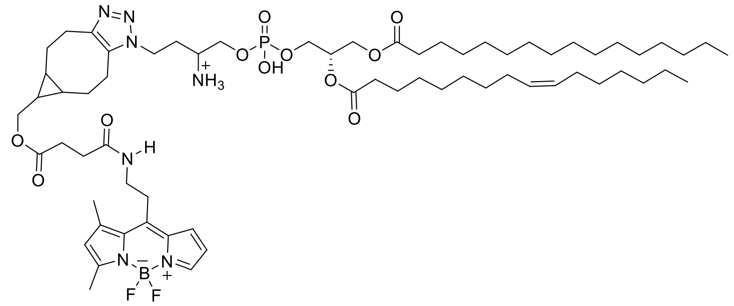


**TLC analysis of fluorescently labelled 19-PE lipids spot denoted by "#" in Figure 2B of the main text**

We isolated a minor spot depicted as "#" in TLC shown in Fig 2B and LC-MS analysis of the extracted spot revealed masses corresponding to fluorescently labelled carboxymethylated **19**-PE lipids.

Carboxymethylation has been reported previously on PE lipids and it involves a non-enzymatic reaction of PE with glucose depicted below[5,6].

**Scheme S12.** Carboxymethylation of PE lipids.

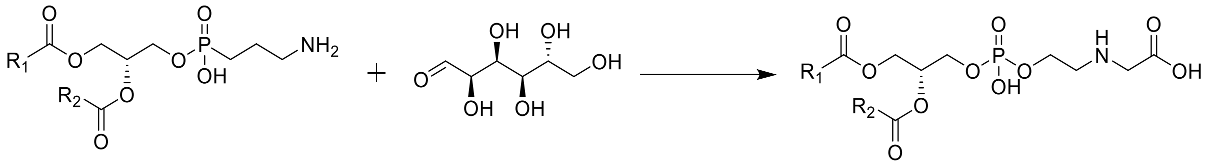

Expected masses of fluorescently labelled, carboxymethylated **19**-PE lipids.

1. 34:1 carboxymethyl **19**-labelled lipid conjugated to BCN- BCN-BODIPY

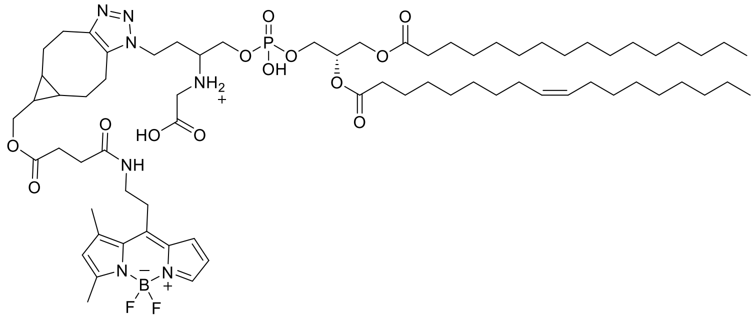

Exact mass: 1340.83

2. 36:2 carboxymethyl **19**-labelled lipid conjugated to BCN-BODIPY

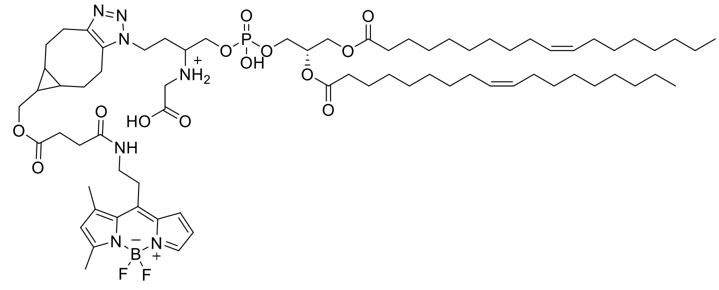

Exact mass: 1366.84

3. 32:1 carboxymethyl **19**-labelled lipid conjugated to BCN-BODIPY

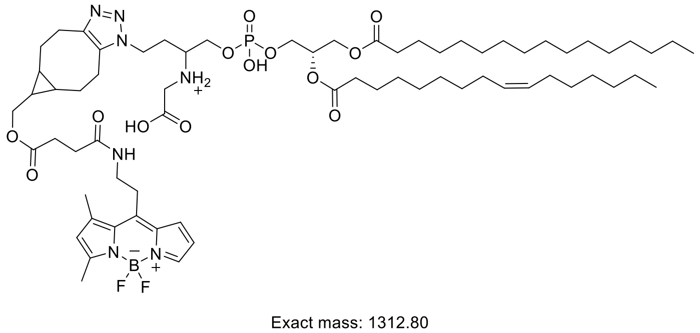

Exact mass: 1312.80

### VIII. Cytotoxicity evaluation of 19 using MTT assay

The cytotoxicity of **19** was assessed in HeLa cells using an MTT-based colorimetric assay, which relies on the reduction of yellow tetrazolium salt (MTT) to purple formazan crystals by metabolically active cells. HeLa cells were seeded in a 96-well plate at a density of 15,000 cells per well and incubated overnight at 37 °C in a humidified atmosphere containing 5% $CO_2$ to allow for cell adherence. The cells were then treated with varying concentrations (10 µM, 100 µM, 200 µM, 2 mM, and 4 mM) of **19** administered from a 65 mM stock prepared in complete medium, and the incubation was performed for 24 h in the incubator. Each concentration was tested in triplicate. Following the incubation, the culture medium was aspirated, and serum-free medium (50 µL) was added to each well to minimize interference from serum components. Subsequently, MTT solution (150 µL of 0.5 mg/mL stock prepared in serum-free DMEM) was added to each well, and the plate was protected from light and incubated at 37 °C for 4 h to allow for formazan crystal formation. After incubation, the MTT solution was carefully removed, and 150 µL of DMSO was added to each well to solubilize the formazan crystals by gentle rocking. Absorbance was measured at 570 nm using a microplate reader. The experimental setup included appropriate controls: blank wells containing medium and MTT reagent without cells to account for background absorbance, untreated cells serving as the negative control (100% viability), and cells treated with 2% Triton X-100 as a positive control. Cell viability (%) was calculated by subtracting the blank absorbance and normalizing the values of treated samples to the untreated control.

### IX. shRNA-mediated VPS13A knockdown in HeLa cells:

HeLa cells were seeded in 3 cm tissue culture dishes at a density of 0.3 × $10^6$ cells/mL and grown to 60–70% confluency in DMEM supplemented with 10% FBS and 1% penicillin–streptomycin at 37 °C in a humidified atmosphere containing 5% $CO_2$ for 12 h. Transfection was performed using polyethyleneimine (PEI). For each dish, the DNA–PEI transfection complex was prepared by diluting 2 µg of VPS13A shRNA plasmid DNA in 125 µL Opti-MEM (Solution A) and 10 µg PEI (from a 1 mg/mL stock) in 125 µL Opti-MEM (Solution B). Solution A was added to solution B, mixed gently by pipetting, and incubated at room temperature for 20 min to allow complex formation. The resulting 250 µL DNA–PEI complex was added dropwise to the cells to ensure even distribution, and the cells were incubated for 6 h at 37 °C in a 5% $CO_2$ incubator. Following incubation, the medium was replaced with fresh complete DMEM. Cells were further incubated for 72 h post-transfection, with medium replacement every 24 h. A control experiment was performed in parallel using a non-targeting (scrambled) shRNA plasmid following the same procedure. The VPS13A shRNA target sequence was: CCGGGCAGAGAAGAAGCTAAAGATTCTCGAGAATCTTTAGCTTCTTCTCTGCTTTTTG.

After 72 h of transfection, live-cell imaging experiments were performed using BCN-BODIPY and Cy5-DBCO as described above. For lipidomics experiments in VPS13A knockdown cells, cells were cultured in 10 cm dishes and processed following the same procedure described above. After 72 h, cells were treated with 100 µM D4-ethanolamine for 24 h followed by lipid extraction using Bligh-dyer method. For mitochondrial lipidomics, mitochondria were isolated as described below followed by lipid extraction.

To check the knockdown efficiency, cells were washed with 1X PBS (2 × 1 mL), scraped into 1 mL of 1X PBS, and pelleted by centrifugation at 1000g for 3 min at 4 °C. The supernatant was discarded, and the cell pellet was resuspended in 80 µL of 1x Roche protease inhibitor cocktail (1× PIC), followed by incubation on ice for 30 min. The lysates were then sonicated at 50% amplitude using 5 s ON/ 5 s OFF cycles for a total duration of 1 min 20 s. Protein concentration was determined using the Bradford assay followed by western blot analysis described below.

Equal amounts of protein (35 µg per sample) were resolved on a 4–15% gradient SDS-PAGE gel for VPS13A KD samples, followed by transfer onto a PVDF membrane at 120 V (≈45 mA) for 12 h at 4 °C. The membrane was washed with 0.05% PBST (3 × 10 min) and blocked with 5% BSA in 1X PBS for 1 h at room temperature. Subsequently, the membrane was incubated with primary antibody for 3 h at room temperature, followed by washing with 0.05% PBST (3 × 10 min). The membrane was then incubated with HRP-conjugated secondary antibody for 2 h at room temperature and washed again with PBST (3 × 10 min). Protein bands were detected using a ECL pierce western blot detection kit, and images were acquired using a Bio-Rad ChemiDoc imaging system. The details of all the antibodies used are shown in **Table S1**.

#### X. Mitochondrial isolation using percoll gradient

Mitochondria was isolated according to the previously published procedure[7]. Following transfection with VPS13A shRNA plasmid, cells were treated with 100 µM D4-ethanolamine for 24 h. Cells were washed with ice-cold DPBS (1X PBS supplemented with 1 mM $CaCl_2$ and 1 mM $MgCl_2$) and collected in 4 mL of ice-cold lysis buffer (5 mM HEPES, 0.5 mM EGTA, 250 mM mannitol, and 0.1% BSA, pH 7.4, containing protease inhibitors) by gentle scraping. The cell suspension was homogenized by sonication at 15% amplitude (3 pulses of 10 s each) and centrifuged at 600g for 10 min to remove nuclei and unbroken cells. The resulting supernatant was collected and subjected to centrifugation at 10,300g for 10 min to pellet crude mitochondria. The mitochondrial pellet was resuspended in 1 mL of lysis buffer and layered onto an 18% percoll solution in polycarbonate tubes, followed by centrifugation at 95,000g for 30 min using an SW40Ti rotor. Two distinct bands were obtained, corresponding to mitochondria-associated membranes (upper band) and purified mitochondria (lower band). The mitochondrial fraction was collected, washed twice by centrifugation at 10,000g for 10 min, and resuspended in mitochondrial resuspension buffer (5 mM HEPES, 0.5 mM EGTA, and 250 mM mannitol, pH 7.4, containing protease inhibitors) for downstream analyses. Protein concentration in all fractions was determined using the BCA assay. The purity of mitochondrial preparations was assessed by western blotting using established organelle-specific marker proteins. Lipids from isolated mitochondrial fractions were extracted using the Bligh–Dyer method and subjected to lipidomic analysis.

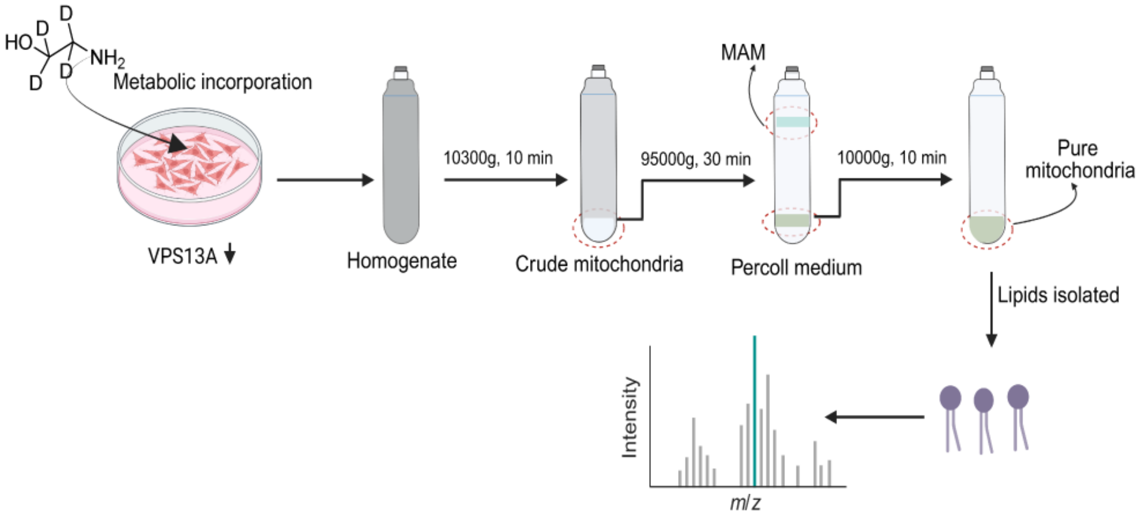


**Figure S3**: Workflow for mitochondrial fractionation

### XI. Lipidomics data

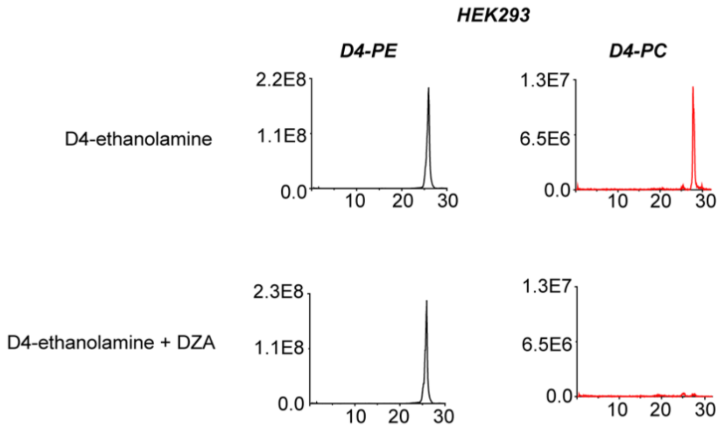


**Figure S4.** Results of LC/MS lipidomics experiments performed on lipids isolated from HEK293 cells subjected to the workflow depicted in **Figure 1B** (top) of the main text. The data demonstrates that both D4-PE and D4-PC lipids are biosynthesized (top panel), and whereas DZA has negligible effect on D4-PE formation, it potently inhibits D4-PC formation in HEK293 cells (bottom panel).

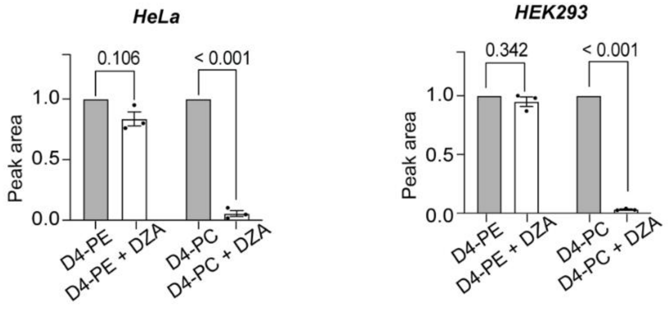


**Figure S5**. Bar graphs representing quantitative comparisons over three replicates for HeLa (left) and HEK293 (right) cells based on lipidomics results obtained from experiments performed according to the workflow depicted in **Figure 1B** (top) of the main text.

**Table S4.** Compound parameters used for the NL/PIS of lipids labelled with all the ethanolamine analogs studied:

| Analogs | Labelled structures | Scan mode NL/PIS (Da) | DP (volts) | EP (volts) | CE (volts) | CXP (volts) |
|---|---|---|---|---|---|---|
| | 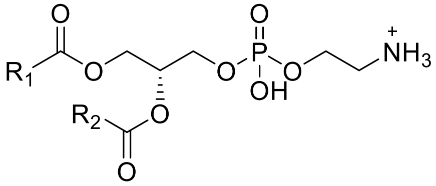 Native **PE** | NL of 141.01 | 125.05 | 10.26 | 27.03 | 23.03 |
| | 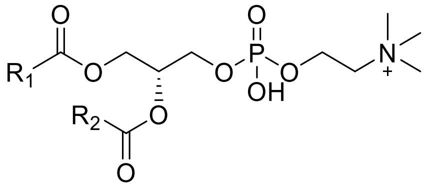 Native **PC** | PIS of 184.07 | 189.53 | 5.88 | 35.05 | 7.04 |
| **1.** | 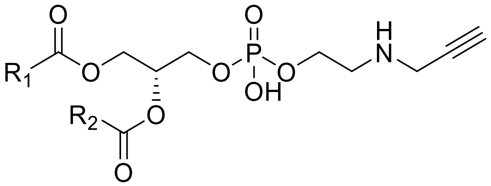 **1**-PE | NL of 179.03 | 120.81 | 10.76 | 31.05 | 24.01 |

| | | | | | | |
|---|---|---|---|---|---|---|
| | 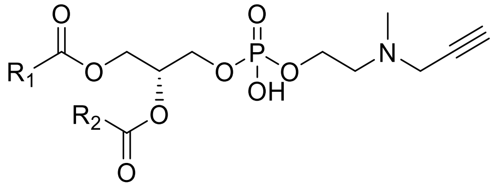 Me-**1**-PE | NL of 193.05 | 114 | 11.8 | 30.60 | 9.30 |
| | 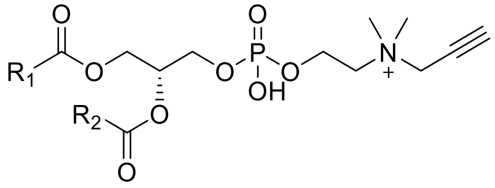 $Me_2$-**1**-PE (PC-derivative) | PIS of 208.1 | 183.60 | 6.10 | 36.70 | 8.40 |
| **2**. | 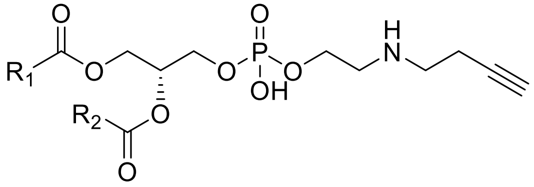 **2**-PE | NL of 193.05 | 50 | 9.90 | 32 | 11.60 |
| | 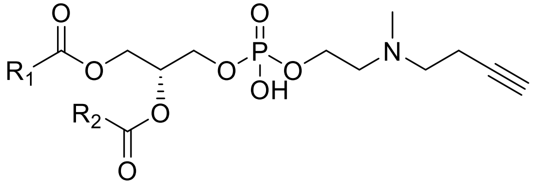 Me-**2**-PE | NL of 207.06 | 35 | 6.88 | 22.36 | 14.80 |

| | | | | | | |
|---|---|---|---|---|---|---|
| | 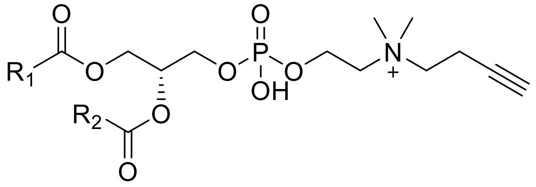 Me$_2$-**2**-PE (PC-derivative) | PIS of 222.08 | 136.32 | 8.98 | 33.58 | 16.77 |
| | 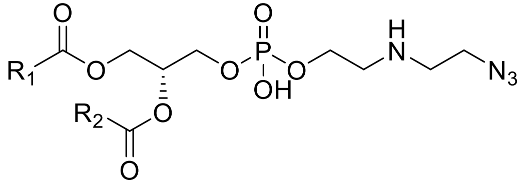 **3**-PE | NL 210.05 | 48.8 | 9.98 | 29.10 | 20.29 |
| **3**. | 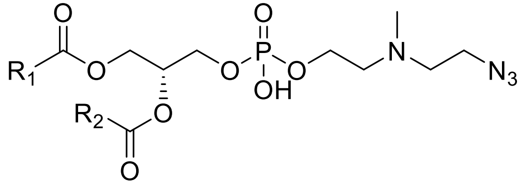 Me-**3**-PE | NL of 224.06 | 30.1 | 7.1 | 27.5 | 15.12 |
| | 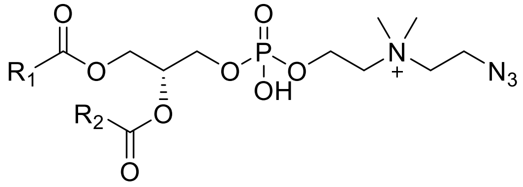 Me$_2$-**3**-PE (PC-derivative) | PIS of 239.09 | 69.27 | 7.76 | 38.20 | 12.90 |

| | | | | | | |
|---|---|---|---|---|---|---|
| **4**. | 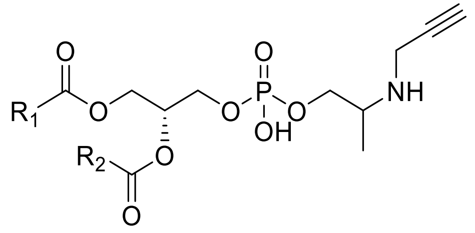 **4**-PE | NL of 193.05 | 47.13 | 7 | 28.85 | 11.41 |
| | 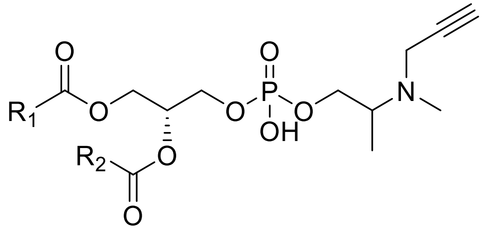 Me-**4**-PE | NL of 207.06 | 42.50 | 6.90 | 27.70 | 12.60 |
| | 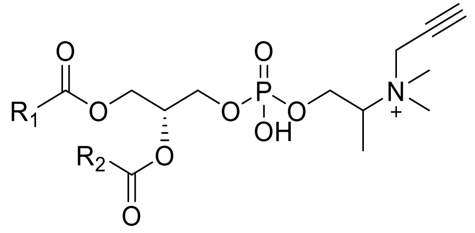 Me$_2$-**4**-PE (PC-derivative) | PIS of 222.08 | 22.2 | 6.9 | 34.5 | 8.08 |

| | | | | | | |
|---|---|---|---|---|---|---|
| | 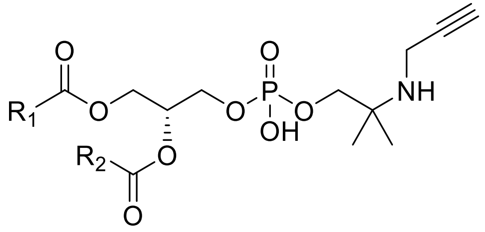 **5**-PE | NL of 207.06 | 150 | 9.1 | 41.9 | 22.9 |
| **5**. | 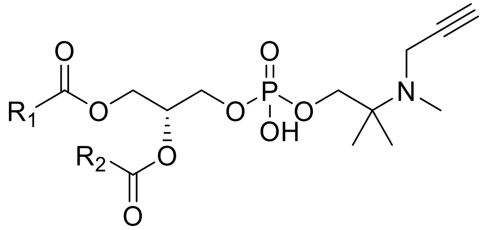 Me-**5**-PE | NL of 221.08 | 33.76 | 8.13 | 21.75 | 11.34 |
| | 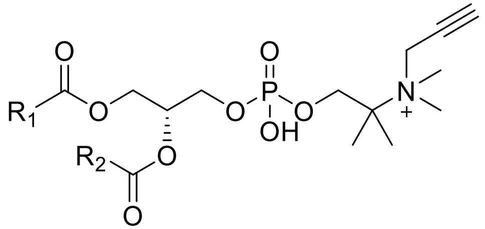 Me$_2$-**5**-PE (PC-derivative) | PIS of 236.1 | 43.25 | 9.80 | 36.10 | 17.4 |
| **6**. | 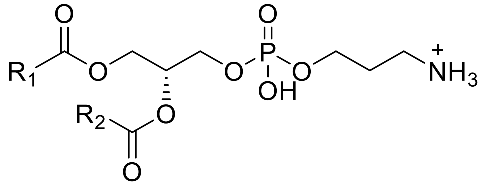 **6**-PE | NL of 155.03 | 102.5 | 13.1 | 30.26 | 22.70 |

| | | | | | | |
|---|---|---|---|---|---|---|
| | 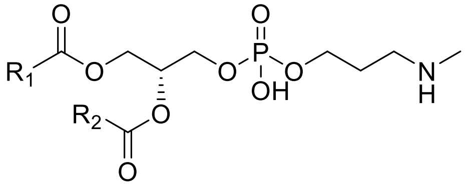<br>Me-**6**-PE | NL of 169.05 | 190.9 | 13.2 | 41.41 | 24.10 |
| | 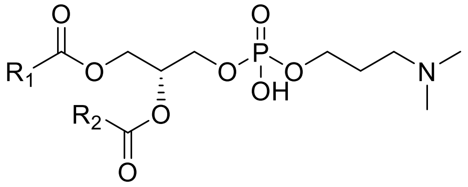<br>$Me_2$-**6**-PE | NL of 183.07 | 194.1 | 10.8 | 42.22 | 24.62 |
| | 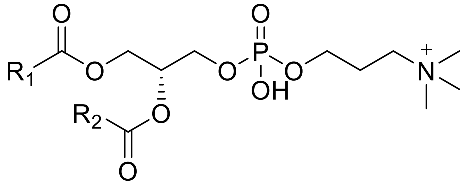<br>$Me_3$-**6**-PE<br>(PC-derivative) | PIS of 198.09 | 62.07 | 7.90 | 48.20 | 22.59 |
| **7**. | 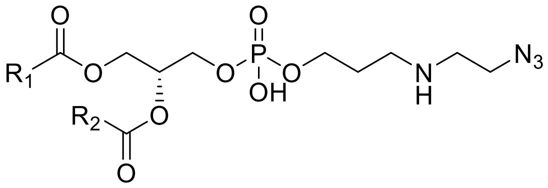<br>**7**-PE | NL of 224.06 | 177.03 | 14.22 | 48.13 | 24.95 |

| | | | | | | |
|---|---|---|---|---|---|---|
| | 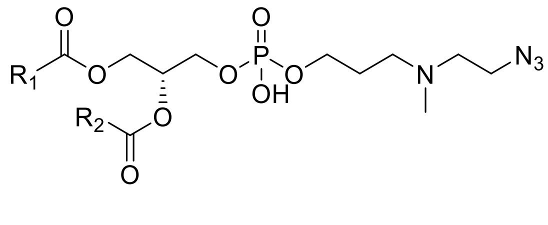 Me-**7**-PE | NL of 238.08 | 123.43 | 11.90 | 41.57 | 23.07 |
| | 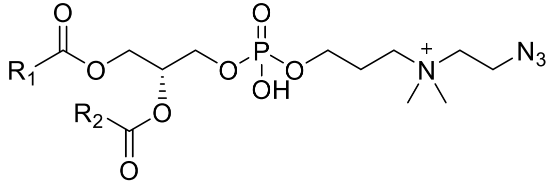 $Me_2$-**7**-PE (PC-derivative) | PIS of 253.10 | 43.30 | 11.80 | 29.40 | 15.09 |
| **8**. | 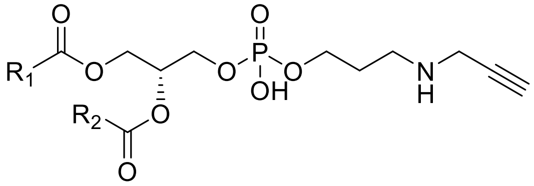 **8**-PE | NL of 193.05 | 83.72 | 6.83 | 23.80 | 13.80 |
| | 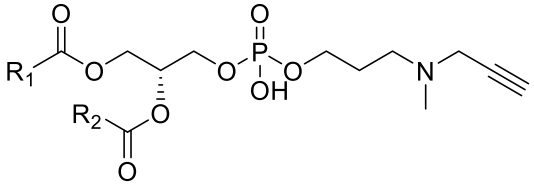 Me-**8**-PE | NL of 207.07 | 30.30 | 11.00 | 21.14 | 18.50 |

| | | | | | | |
|---|---|---|---|---|---|---|
| | 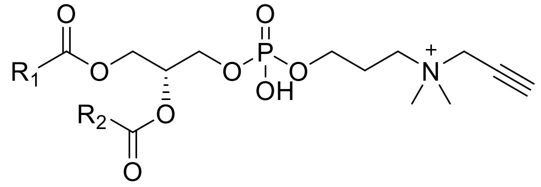 Me$_2$-**8**-PE | PIS of 222.09 | 43.60 | 10.17 | 29.13 | 9.35 |
| **9**. | 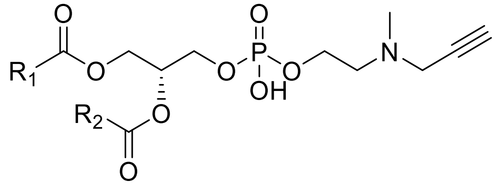 **9**-PE | NL of 193.05 | 114.00 | 11.80 | 30.60 | 9.30 |
| | 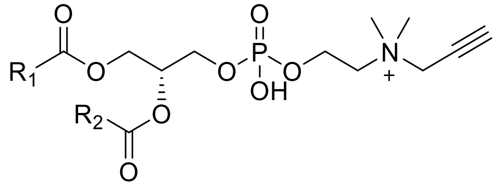 Me-**9**-PE (PC-derivative) | PIS of 208.10 | 183.6 | 6.10 | 36.70 | 8.40 |
| **10**. | 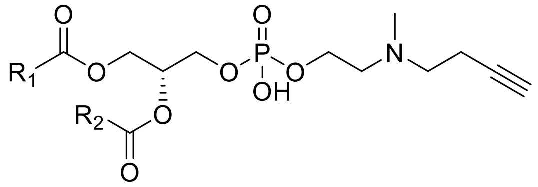 **10**-PE | NL of 207.06 | 151.50 | 9.92 | 41.40 | 9.53 |

| | | | | | | |
|---|---|---|---|---|---|---|
| | 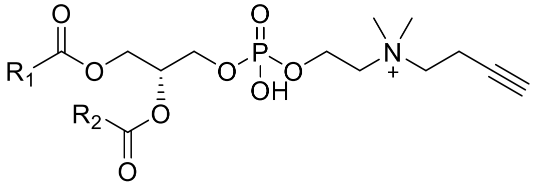 Me-**10**-PE (PC-derivative) | PIS of 222.08 | 191.40 | 10.85 | 39.27 | 14.96 |
| **11**. | 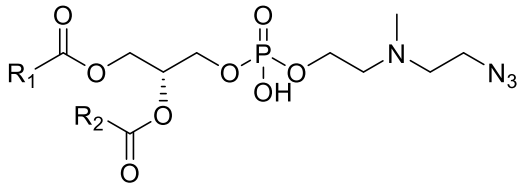 **11**-PE | NL of 224.06 | 174.55 | 10.07 | 32.04 | 8.98 |
| | 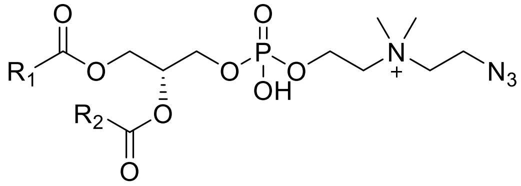 Me-**11**-PE (PC-derivative) | PIS of 239.09 | 129.76 | 5.96 | 26.41 | 10.51 |
| **12**. | 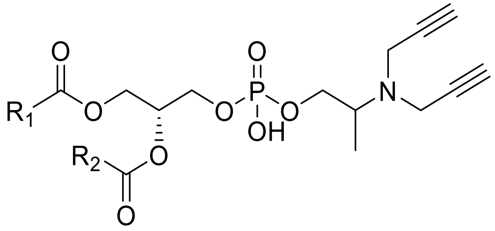 **12**-PE | NL of 231.06 | 39.41 | 11.03 | 21.01 | 15.06 |

| | | | | | | |
|---|---|---|---|---|---|---|
| | 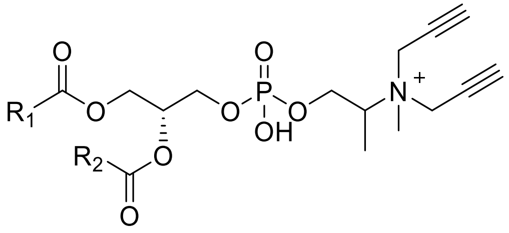<br>Me-**12**-PE<br>(PC-derivative) | PIS of 246.08 | 42.30 | 7.90 | 60.90 | 8.10 |
| **13**. | 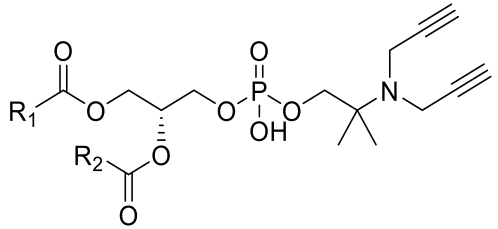<br>**13**-PE | NL of 245.08 | 120 | 9.15 | 34.35 | 9.27 |
| | 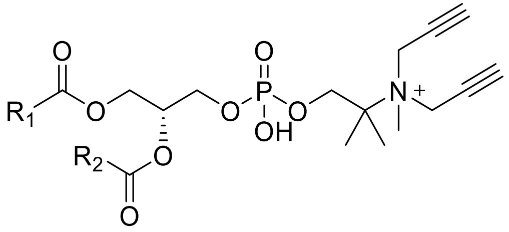<br>Me-**13**-PE<br>(PC-derivative) | PIS of 260.10 | 72.80 | 8.95 | 40.87 | 15.15 |

| | | | | | | |
|---|---|---|---|---|---|---|
| **14**. | 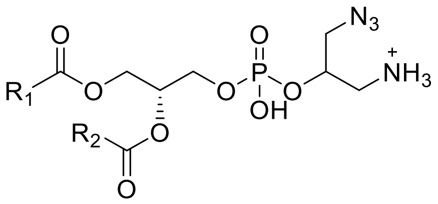 **14**-PE | NL 196.03 | 120 | 6 | 40 | 23.12 |
| | 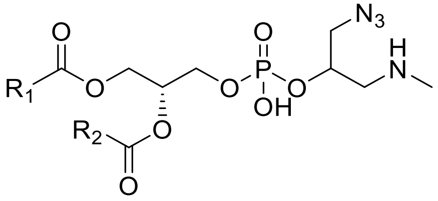 Me-**14**-PE | NL 210.05 | 45.49 | 9.78 | 29.64 | 11.57 |
| | 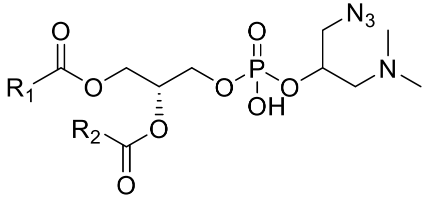 $Me_2$-**14**-PE | NL of 224.06 | 38.7 | 5.97 | 32.94 | 11.93 |
| | 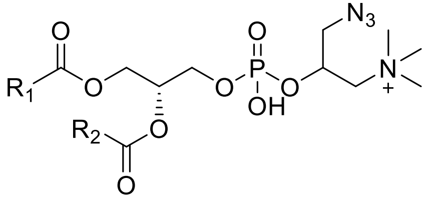 $Me_3$-**14**-PE (PC-derivative) | PIS of 239.09 | 30.5 | 8.98 | 38.96 | 8.82 |

| | | | | | | |
|---|---|---|---|---|---|---|
| **15**. | 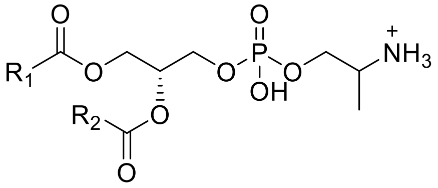 **15**-PE | NL of 155.03 | 130.4 | 10.68 | 21.35 | 22.70 |
| | 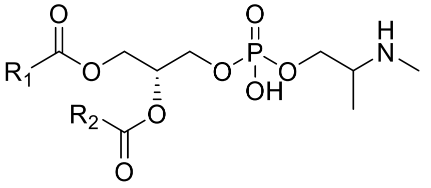 Me-**15**-PE | NL of 169.05 | 181.78 | 12.75 | 41.11 | 24.90 |
| | 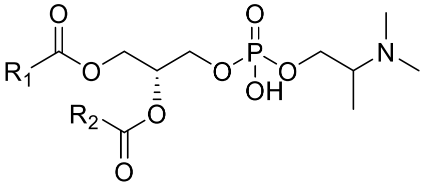 $Me_2$-1**5**-PE | NL of 183.07 | 184.13 | 11.64 | 41.60 | 20.14 |
| | 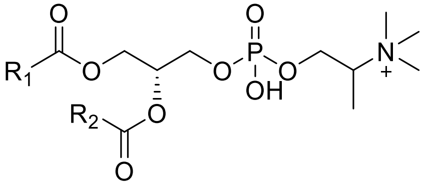 $Me_3$-**15**-PE (PC-derivative) | PIS of 198.09 | 94.4 | 11.10 | 35.60 | 12.90 |

| | | | | | | |
|---|---|---|---|---|---|---|
| **16**. | 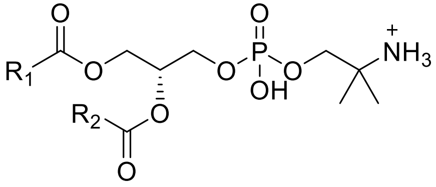 **16**-PE | NL of 169.05 | 152.6 | 5.35 | 29.29 | 20.79 |
| | 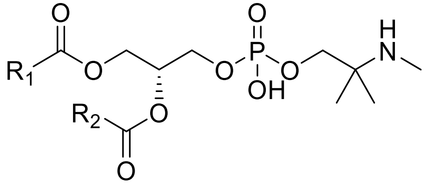 Me-**16**-PE | NL of 183.07 | 158.10 | 11.40 | 42.04 | 23.17 |
| | 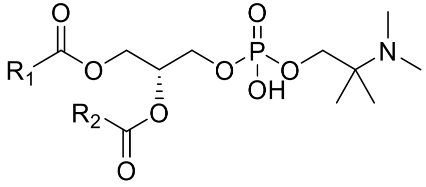 $Me_2$-**16**-PE | NL of 197.08 | 78.70 | 11.30 | 28.80 | 32.37 |
| | 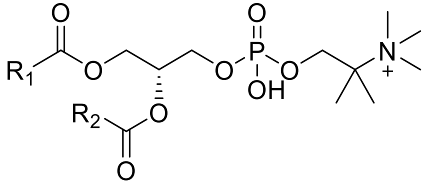 $Me_3$-**16**-PE (PC-derivative) | PIS of 212.10 | 87.90 | 10.10 | 81.29 | 17.09 |

| | | | | | | |
|---|---|---|---|---|---|---|
| **17**. | 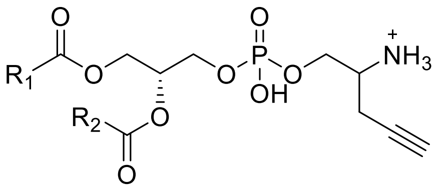<br>**17**-PE | NL of 179.03 | 120.8 | 10.76 | 31.05 | 24.01 |
| | 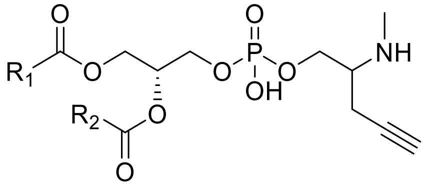<br>Me-**17**-PE | NL 193.05 | 45.49 | 9.78 | 29.64 | 11.57 |
| | 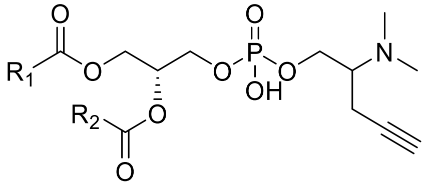<br>$Me_2$-**17**-PE | NL of 207.06 | 38.7 | 5.97 | 32.94 | 11.93 |
| | 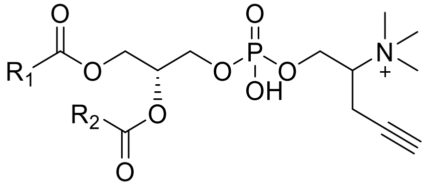<br>$Me_3$-**17**-PE<br>(PC-derivative) | PIS of 222.09 | 79.58 | 11.64 | 37.33 | 5.67 |

| | | | | | | |
|---|---|---|---|---|---|---|
| **18**. | 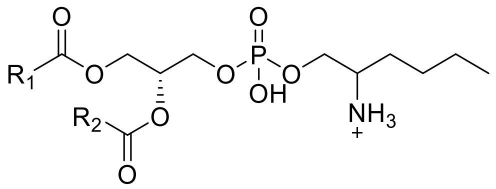 **18**-PE | NL of 197.08 | 134.26 | 6.88 | 31.80 | 21.44 |
| | 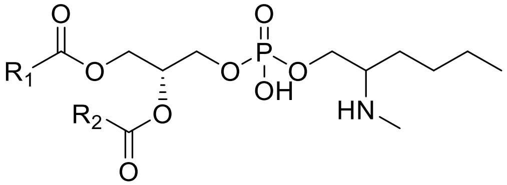 Me-**18**-PE | NL of 211.1 | 59.82 | 13.00 | 30.30 | 22.80 |
| | 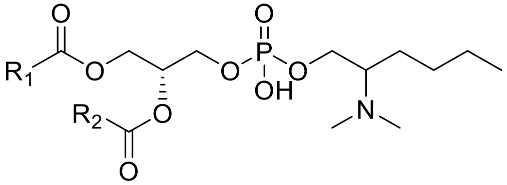 $Me_2$-**18**-PE | NL of 225.1 | 198.28 | 13.00 | 50.13 | 16.31 |
| | 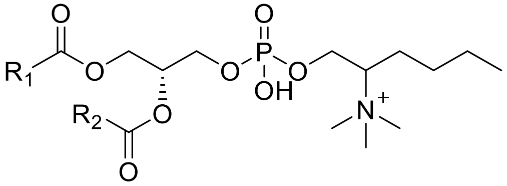 $Me_3$-**18**-PE (PC-derivative) | PIS of 240.14 | 29.70 | 13.10 | 36.30 | 29.23 |

| | | | | | | |
|---|---|---|---|---|---|---|
| **19**. | 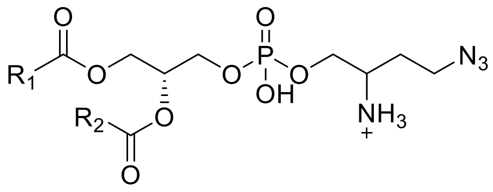<br>**19**-PE | NL of 210 | 140 | 13.59 | 31.34 | 22.90 |
| | 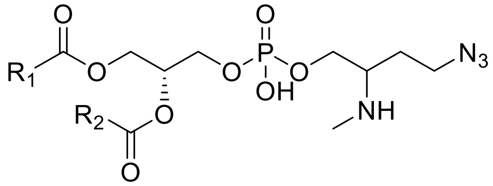<br>Me-**19**-PE | NL of 224.06 | 42.9 | 11.03 | 30.14 | 21.62 |
| | 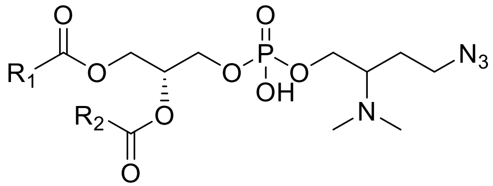<br>$Me_2$-**19**-PE | NL of 238.08 | 47 | 7.9 | 19.56 | 15.19 |
| | 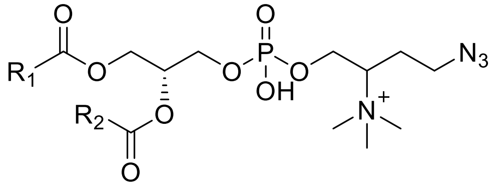<br>$Me_3$-**19**-PE<br>(PC-derivative) | PIS of 253.1 | 40 | 7.18 | 35.07 | 18.03 |

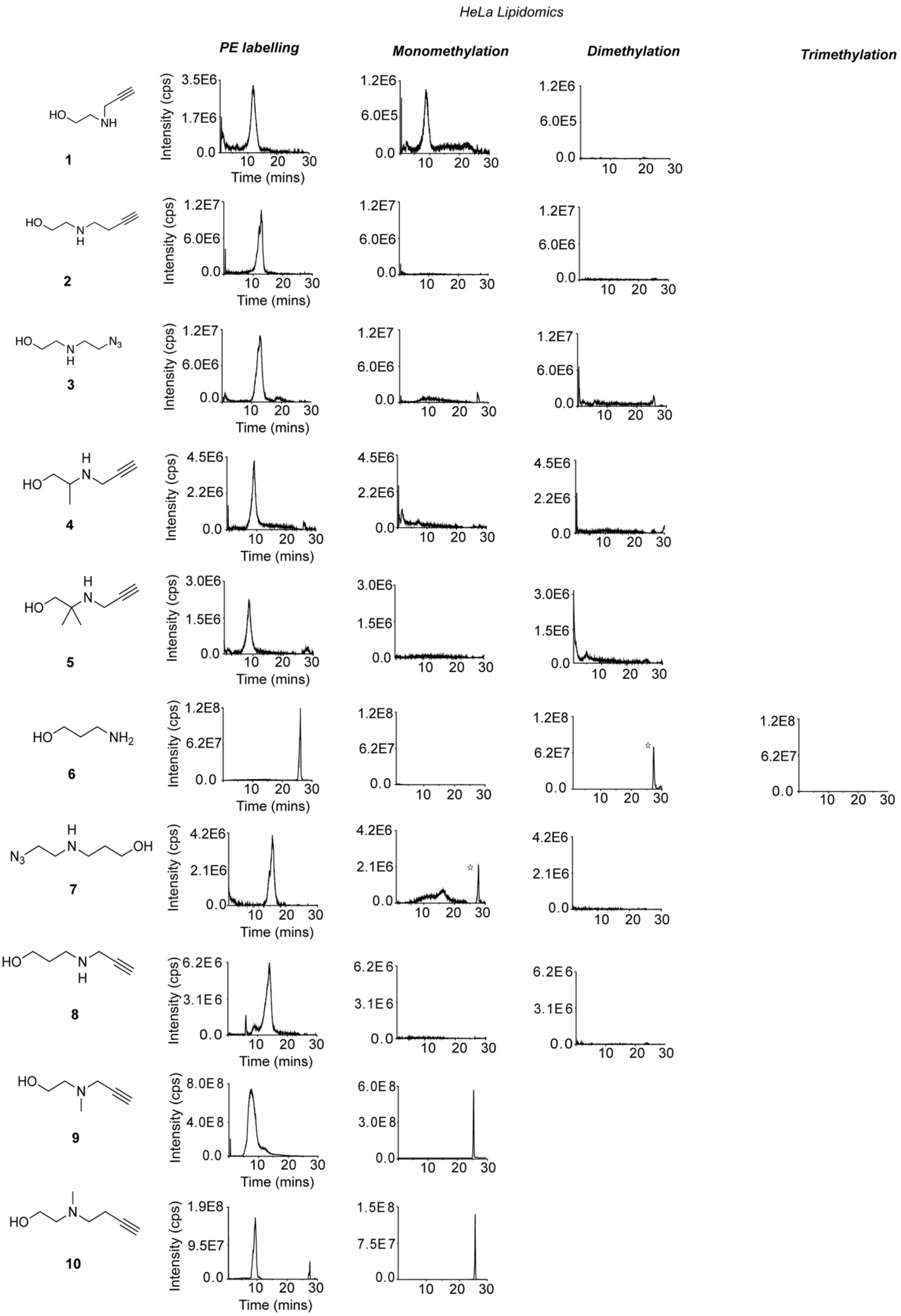
HeLa Lipidomics
PE labelling
Monomethylation
Dimethylation
Trimethylation
Intensity (cps)
Time (mins)
1
2
3
4
5
6
7
8
9
10

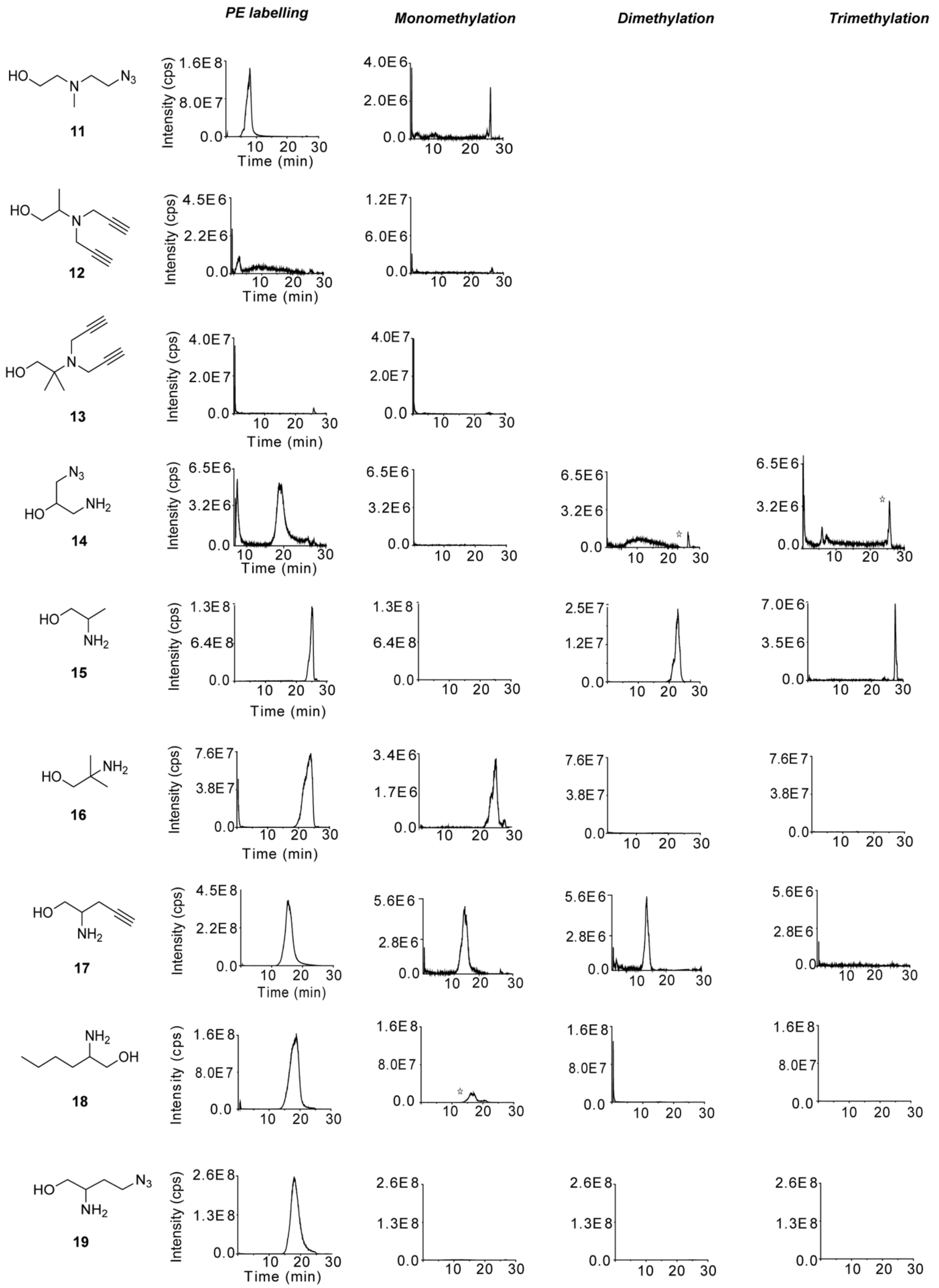


**Figure S6.** Characterization of metabolic labelling of mammalian ethanolamine-containing lipids in HeLa cells. The peaks marked with an asterisk (*) corresponds to a signal that is also observed in control cells not treated with the probe, indicating that it originates from an endogenous lipid species or background signal rather than probe-derived labelling.

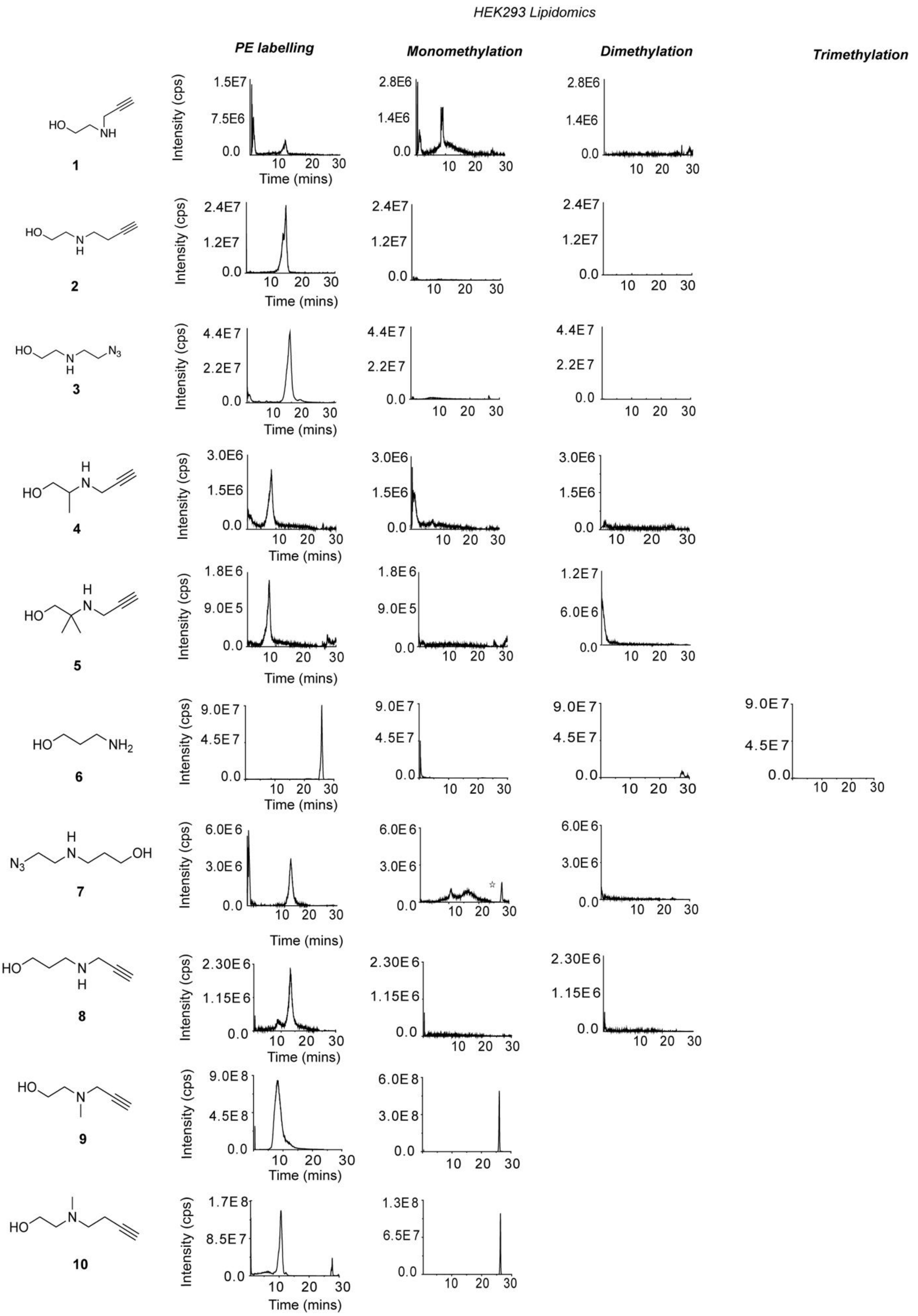
HEK293 Lipidomics
PE labelling
Monomethylation
Dimethylation
Trimethylation
Intensity (cps)
Time (mins)
1
2
3
4
5
6
7
8
9
10

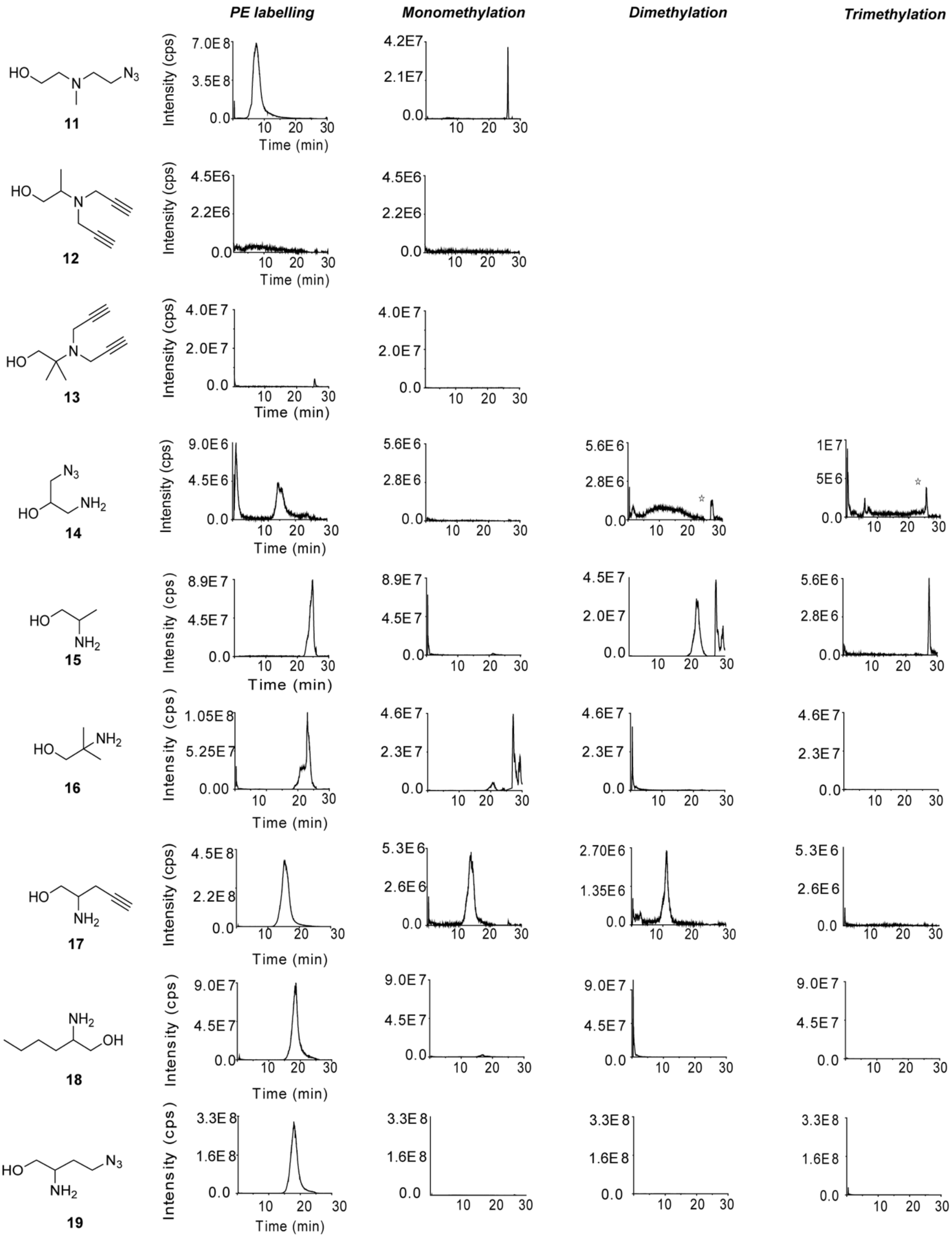


**Figure S7.** Characterization of metabolic labelling of mammalian ethanolamine-containing lipids in HEK293 cells. The peaks marked with an asterisk (*) corresponds to a signal that is also observed in control cells not treated with the probe, indicating that it originates from an endogenous lipid species or background signal rather than probe-derived labelling.

**LipidView analysis for ethanolamine analogs 2, 3, 4, 5, 6, 7, 8, 12, 14, 18, and 19 that exclusively label PE lipids in HeLa cells.**

The peaks for individual labelled PE lipid species have been normalized to the most intense peak intensities observed and the data are presented as mean ± SEM from three independent biological replicates.

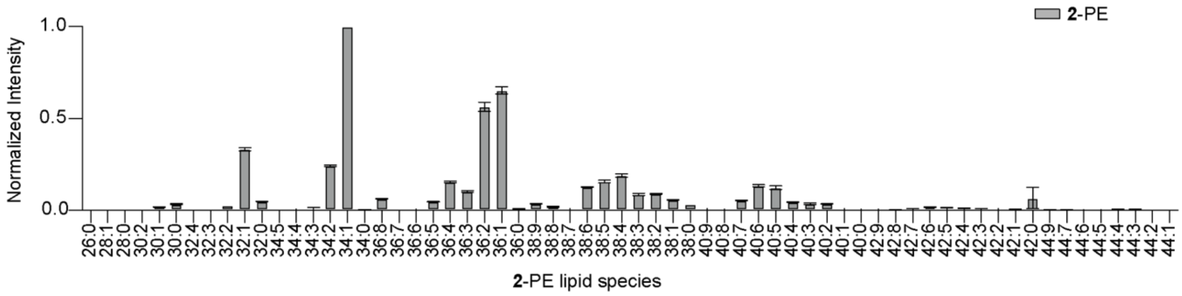


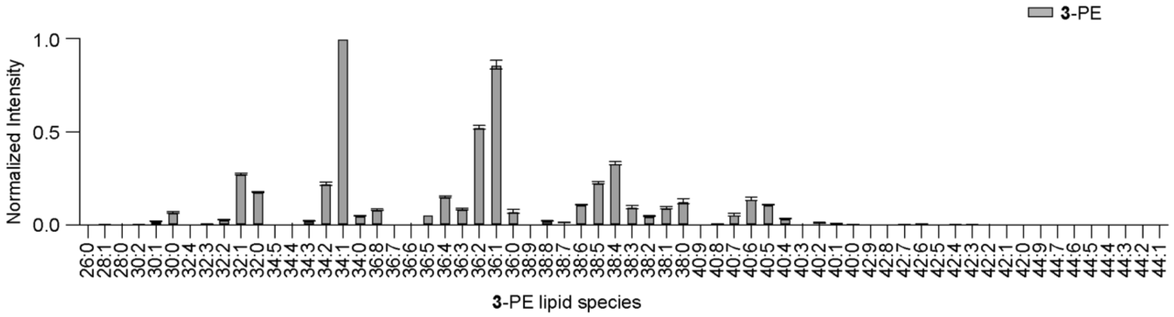


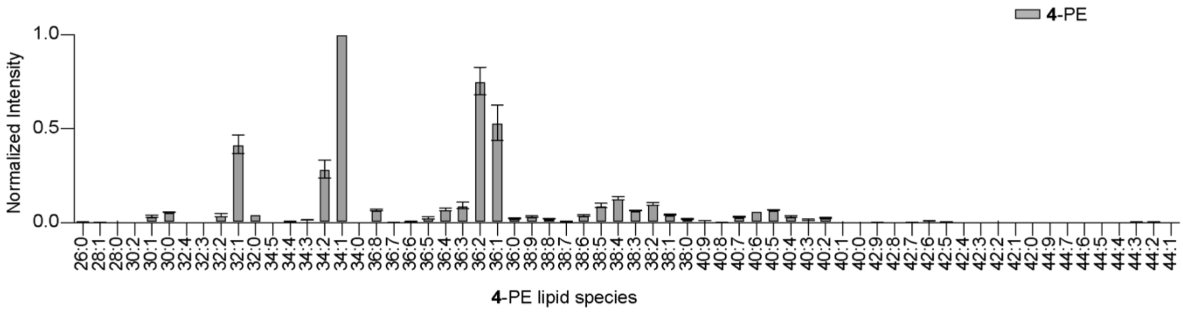


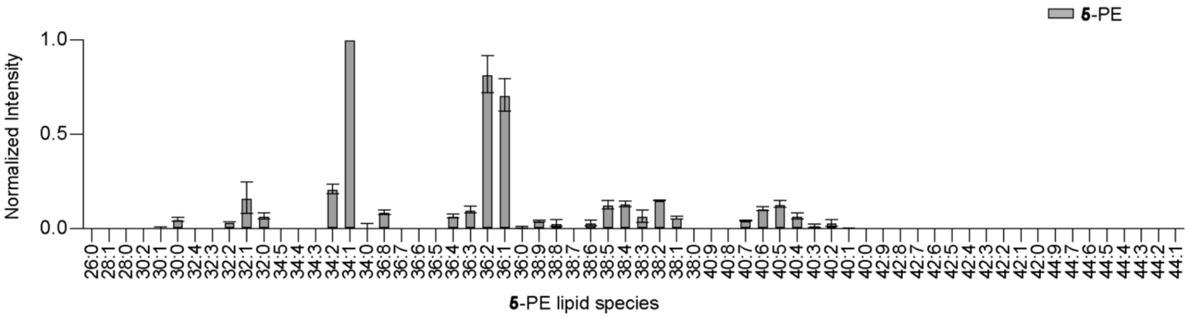

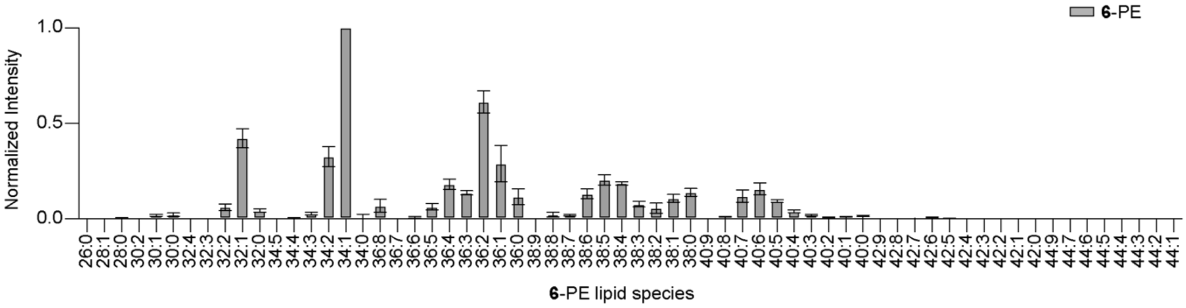
6-PE
Normalized Intensity
1.0
0.5
0.0
6-PE lipid species

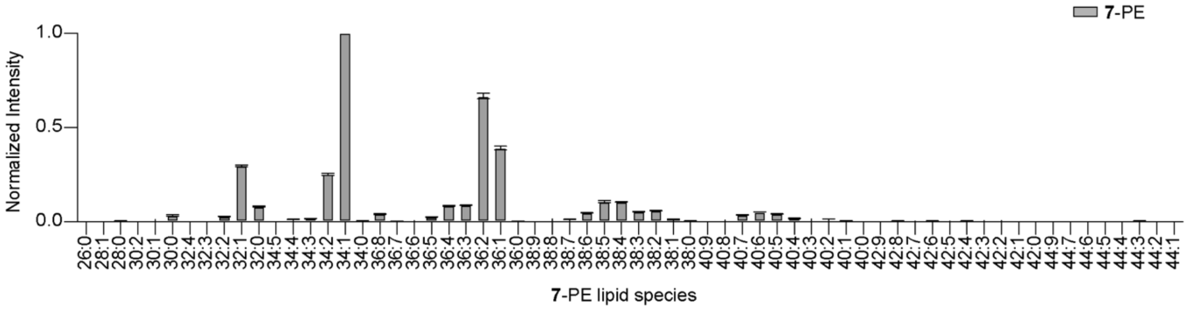
7-PE
Normalized Intensity
1.0
0.5
0.0
7-PE lipid species

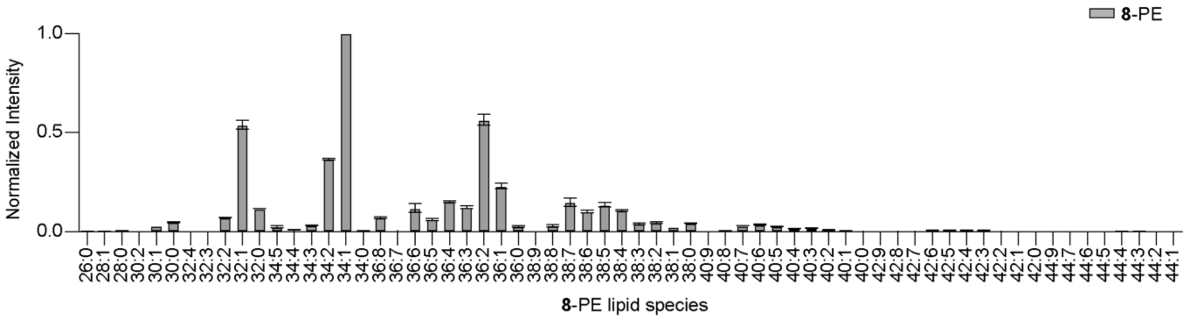
8-PE
Normalized Intensity
1.0
0.5
0.0
8-PE lipid species

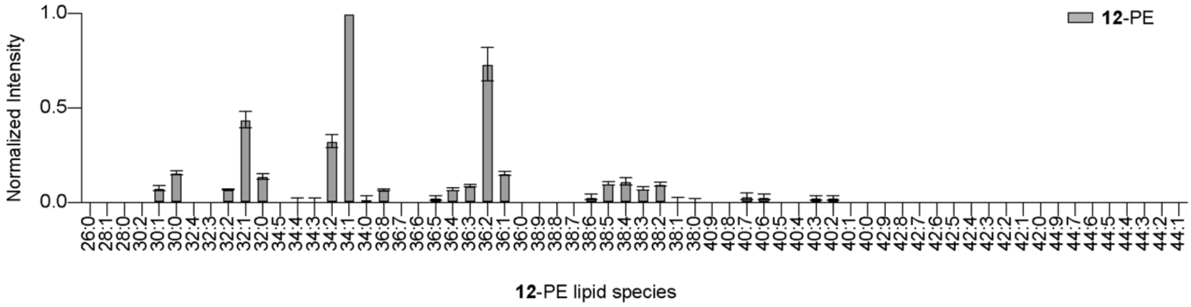
12-PE
Normalized Intensity
1.0
0.5
0.0
12-PE lipid species

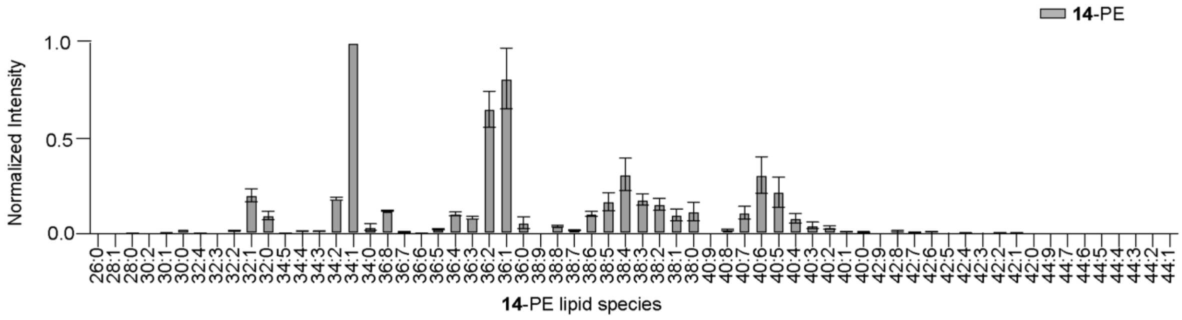
14-PE
Normalized Intensity
1.0
0.5
0.0
14-PE lipid species

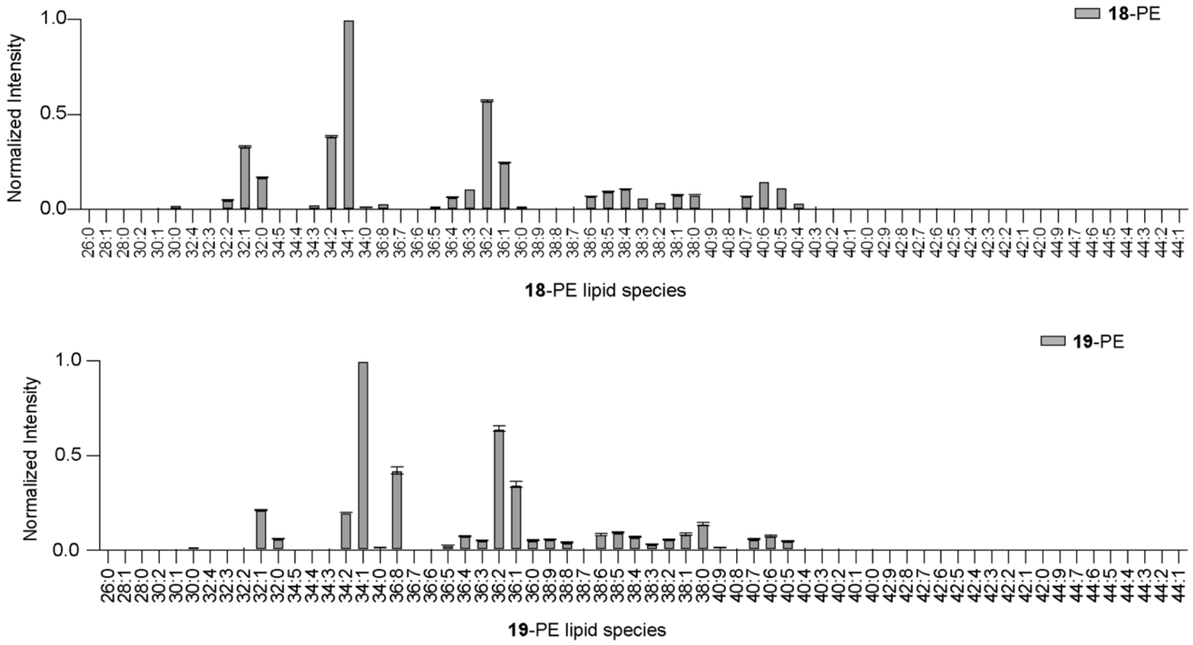


**Figure S8.** Lipidview analysis on analogs administered to HeLa cells.

**LipidView analysis for ethanolamine analogs 2, 3, 4, 5, 6, 7, 8, 12, 14, 18, and 19 that exclusively label PE lipids in HeK293 cells.**

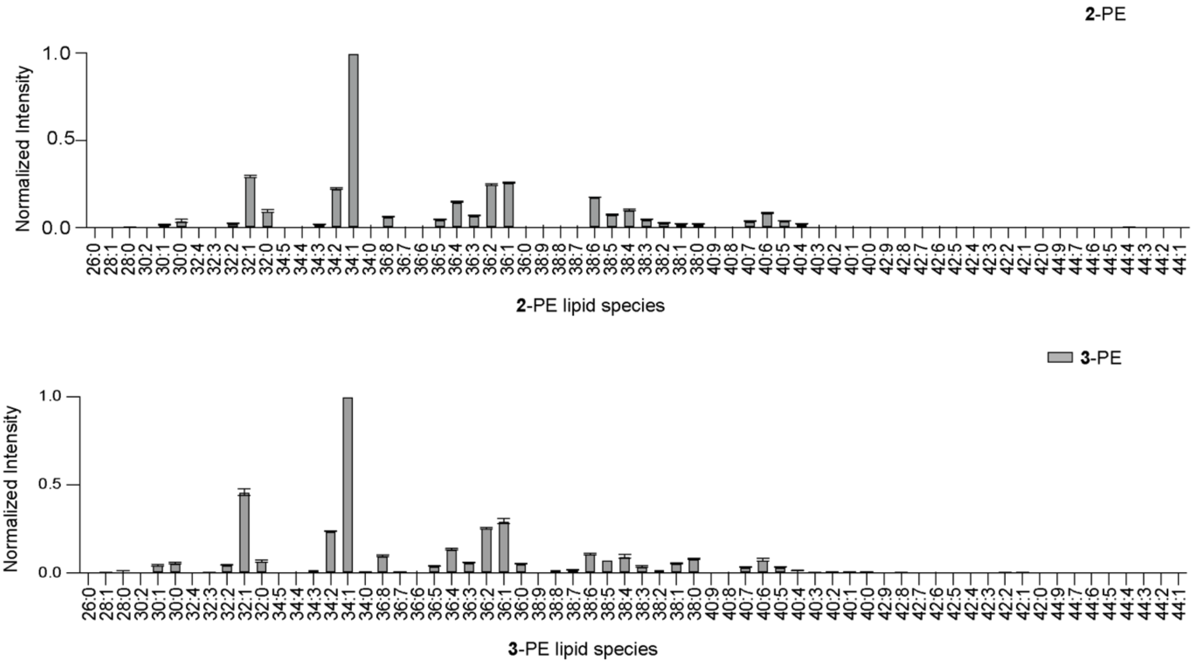

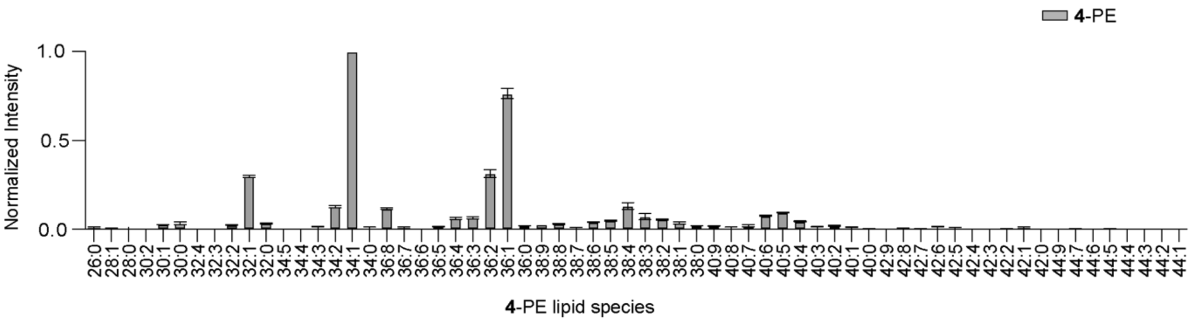
4-PE
Normalized Intensity
4-PE lipid species

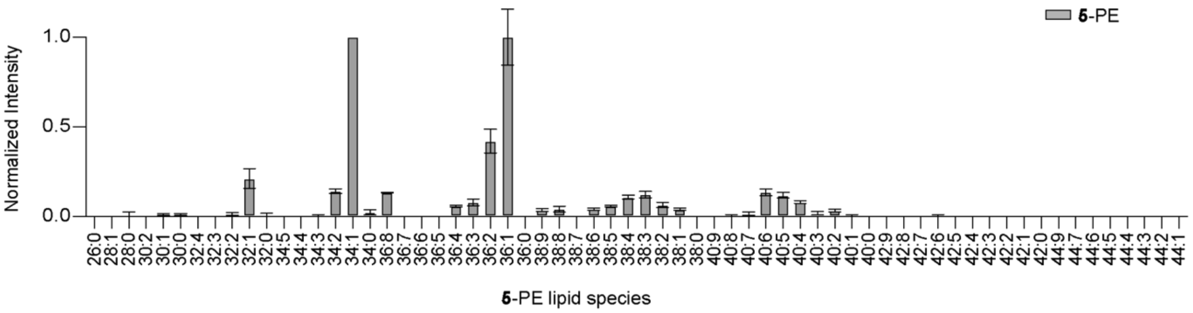
5-PE
Normalized Intensity
5-PE lipid species

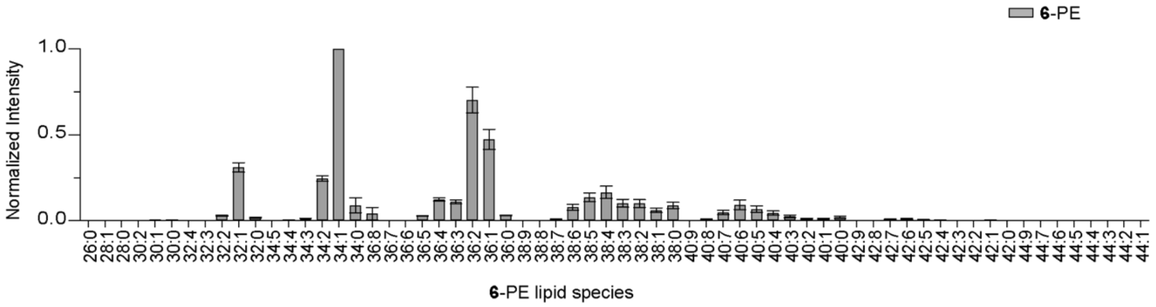
6-PE
Normalized Intensity
6-PE lipid species

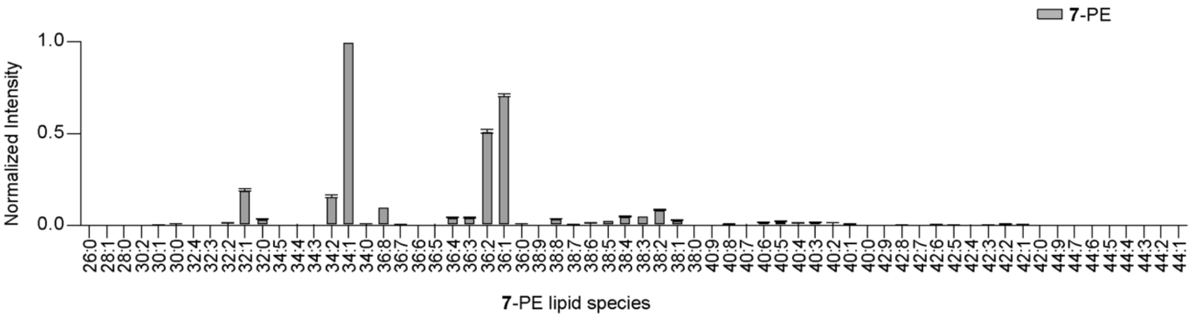
7-PE
Normalized Intensity
7-PE lipid species

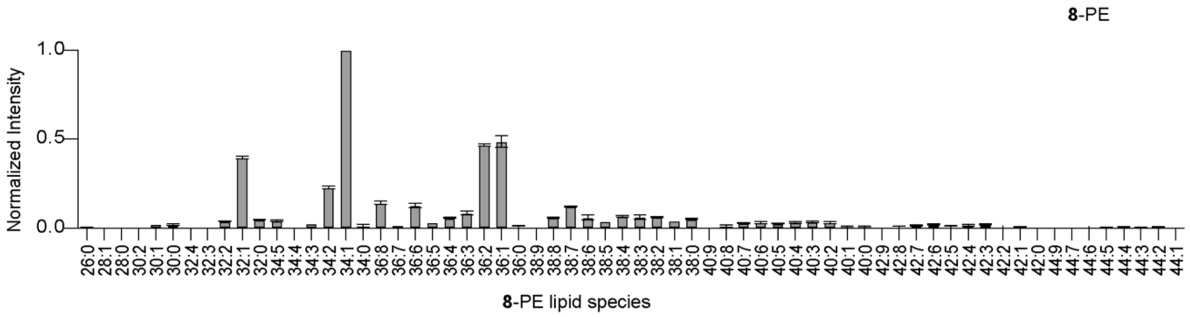
8-PE
Normalized Intensity
8-PE lipid species

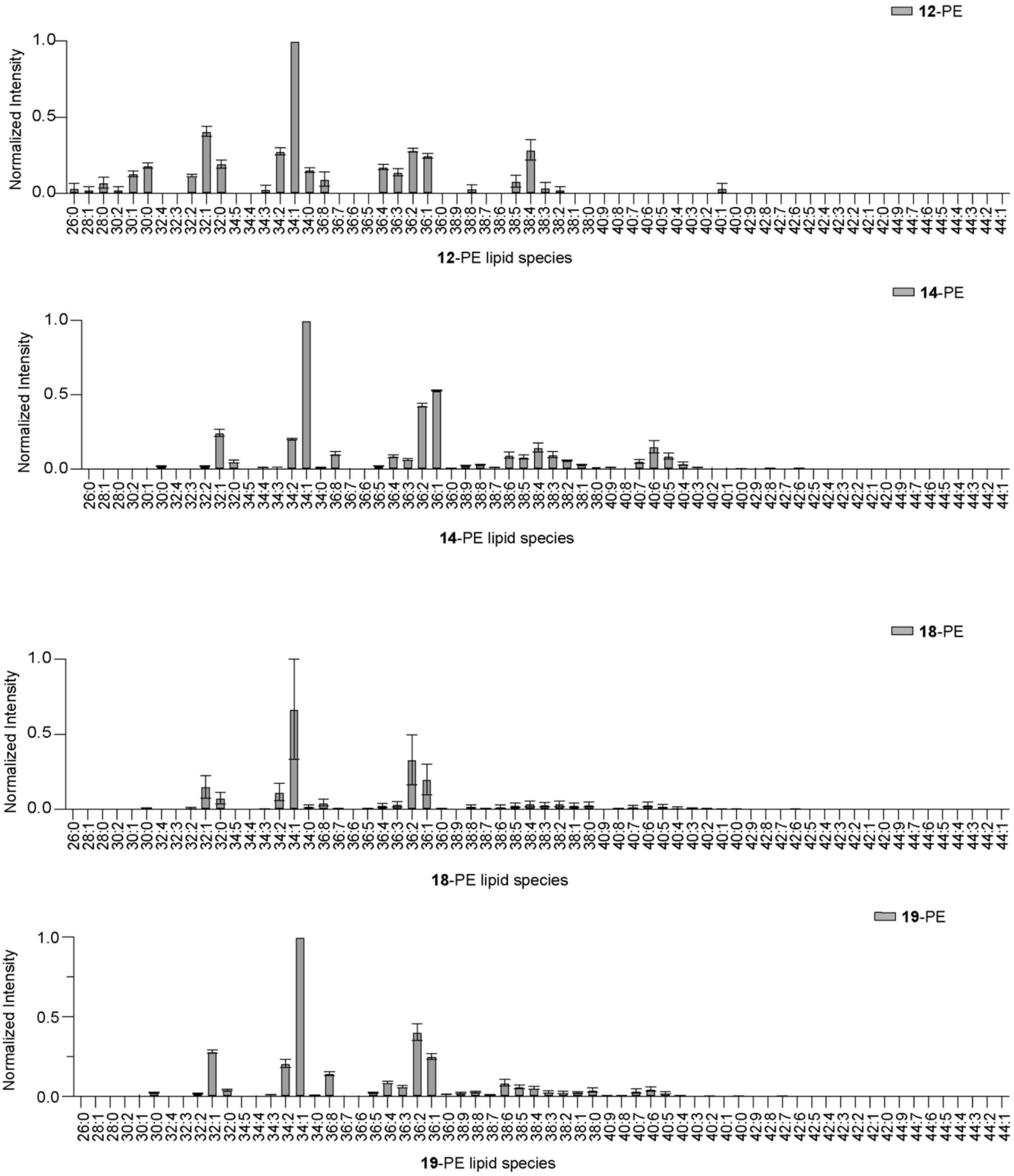


**Figure S9.** Lipidview analysis on analogs administered to HEK293 cells.

### XII. Cytotoxicity studied on compound 19

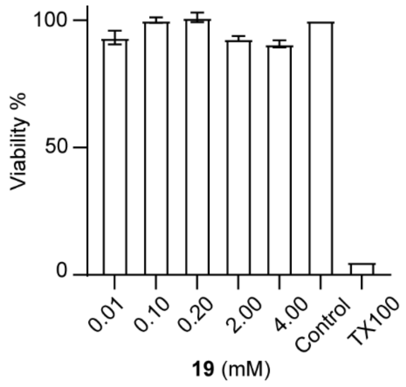


**Figure S10.** MTT assay on HeLa cells administered with compound **19**. The % viability observed at 0.01 mM to 4 mM concentrations of compound 19 with TritonX 100 (TX100) as a positive control.

### XIII. Data for MS analysis on fluorescent TLC spot denoted by "#" in Figure 2B of the main text

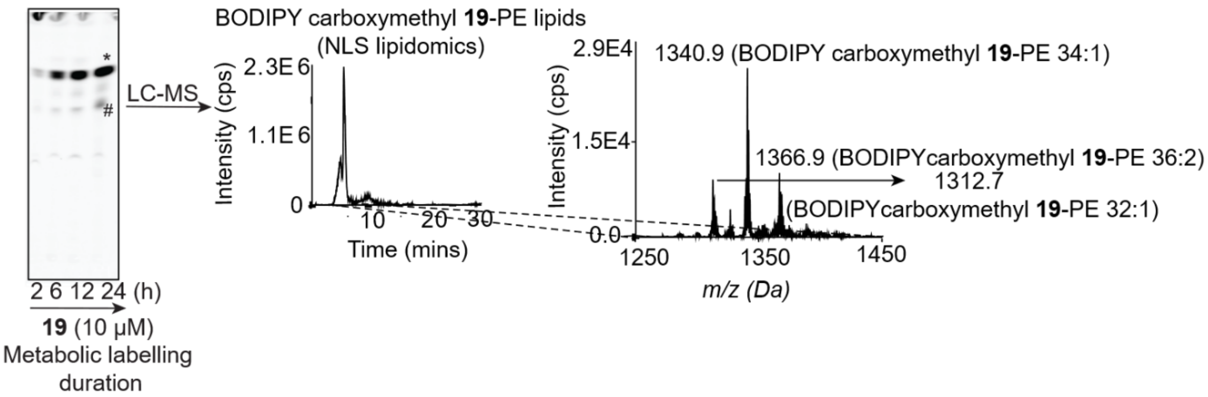


**Figure S11.** LC (middle panel) and MS spectra on the LC peak (right panel) obtained upon injection of the TLC spot "#" isolated from the TLC (left panel).

### XIV. Western blot on VPS13A knockdown cells

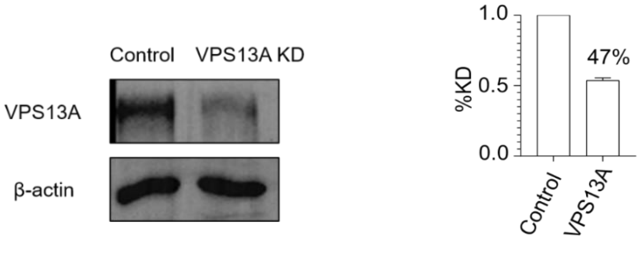


**Figure S12:** Left: Western blot showing VPS13A knockdown. Right: Knockdown measured via densitometry in three replicates.

**XV. Western blot analysis on purified mitochondrial fractions obtained by performing percoll gradient fractionation on VPS13KD and control HeLa cells**

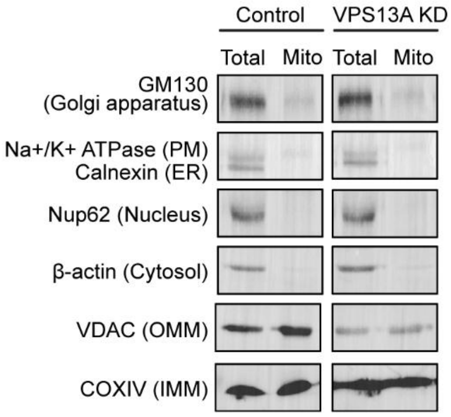


**Figure S13**: Western blot performed by using markers for different organellar proteins on the total and purified mitochondrial fractions. (PM: plasma membrane, ER: endoplasmic reticulum, OMM: outer mitochondrial membrane, IMM: inner mitochondrial membrane, Mito: mitochondria). The purified mitochondria sample lane does not depict any bands for proteins belonging to other organelles whereas bands for mitochondrial proteins, VDAC and COXIV are clearly visible.

### XVII. $^{1}$H and $^{13}$C NMR spectra

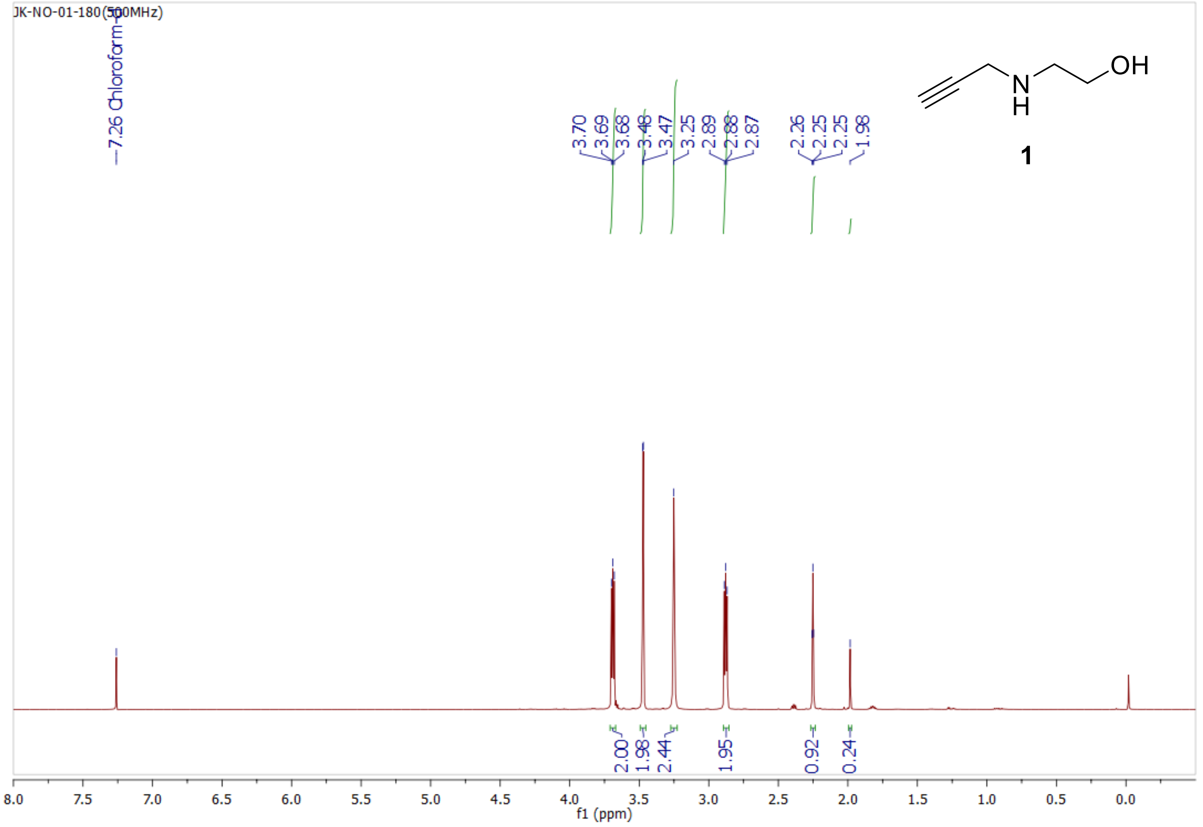


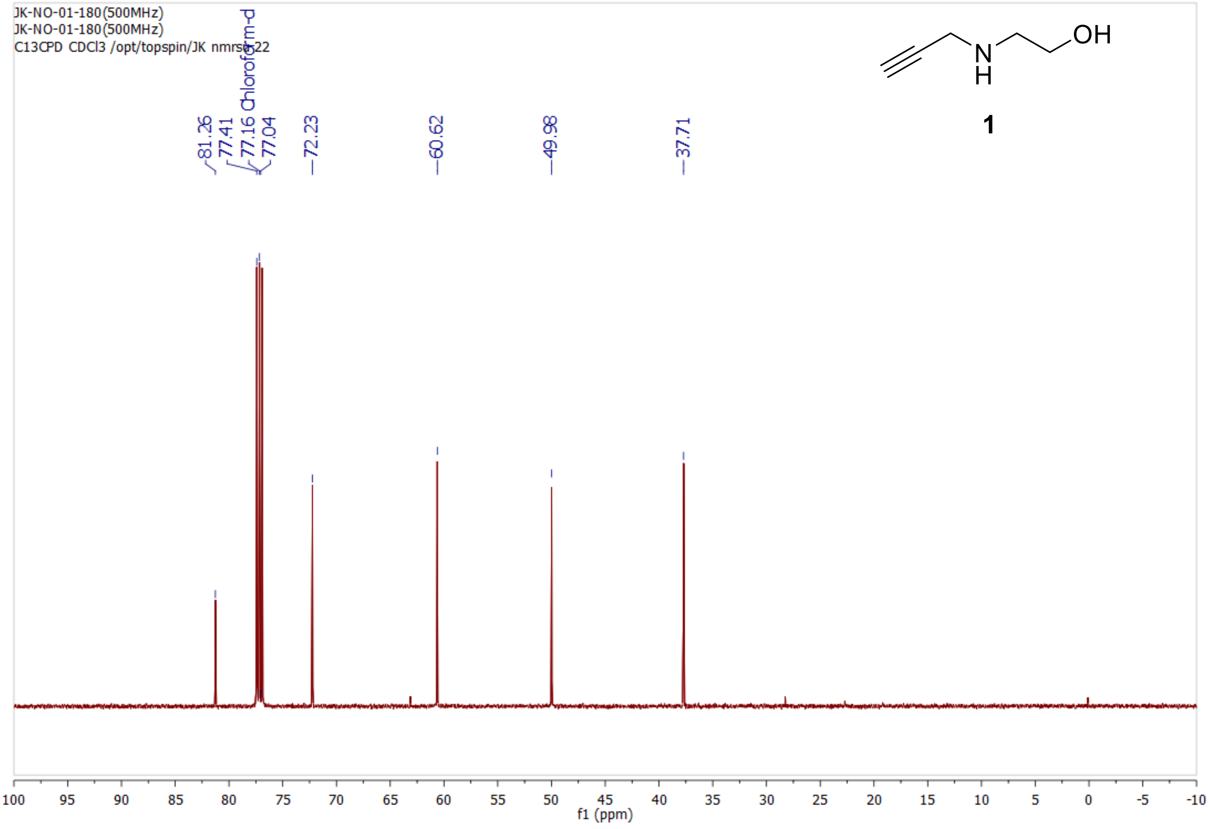

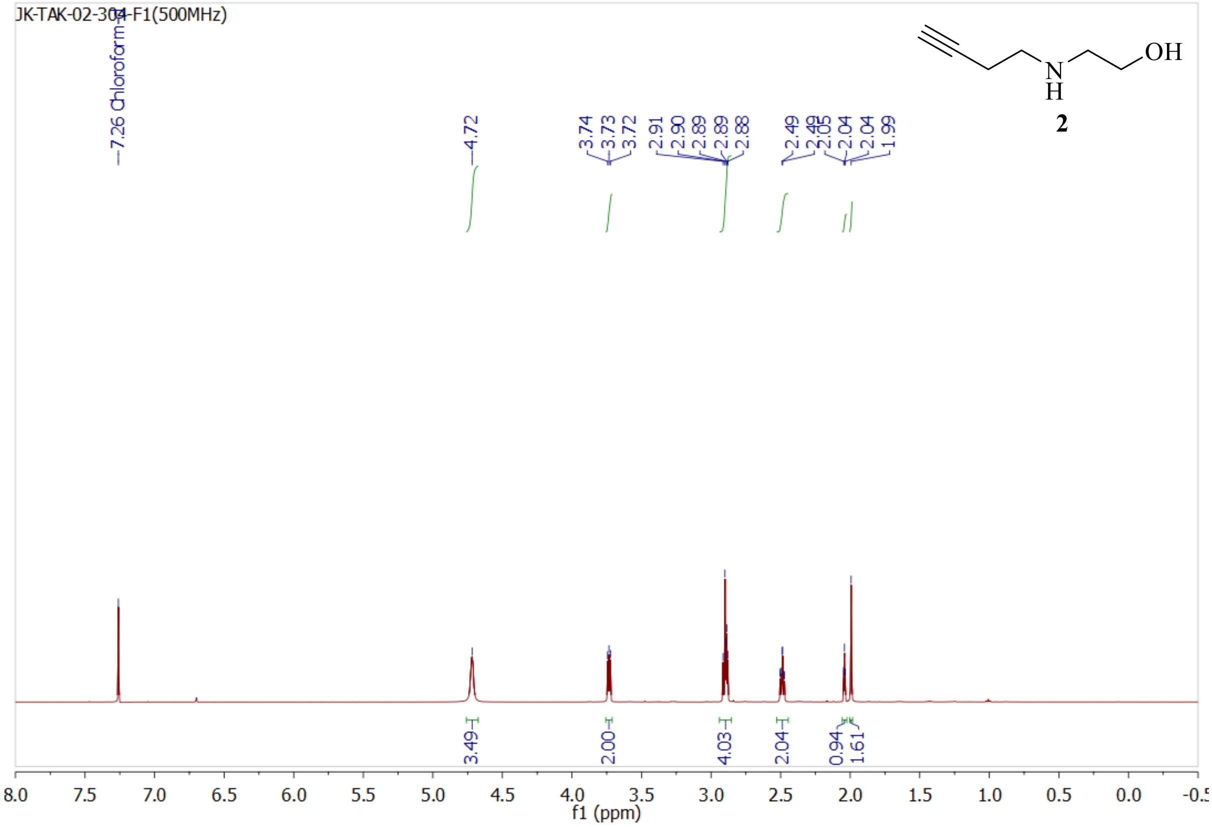
JK-TAK-02-304-F1(500MHz)
7.26 Chloroform-d
4.72
3.74
3.73
3.72
2.91
2.90
2.89
2.89
2.88
2.49
2.49
2.05
2.04
2.04
1.99
3.49
2.00
4.03
2.04
0.94
1.61
f1 (ppm)
N
H
OH
2

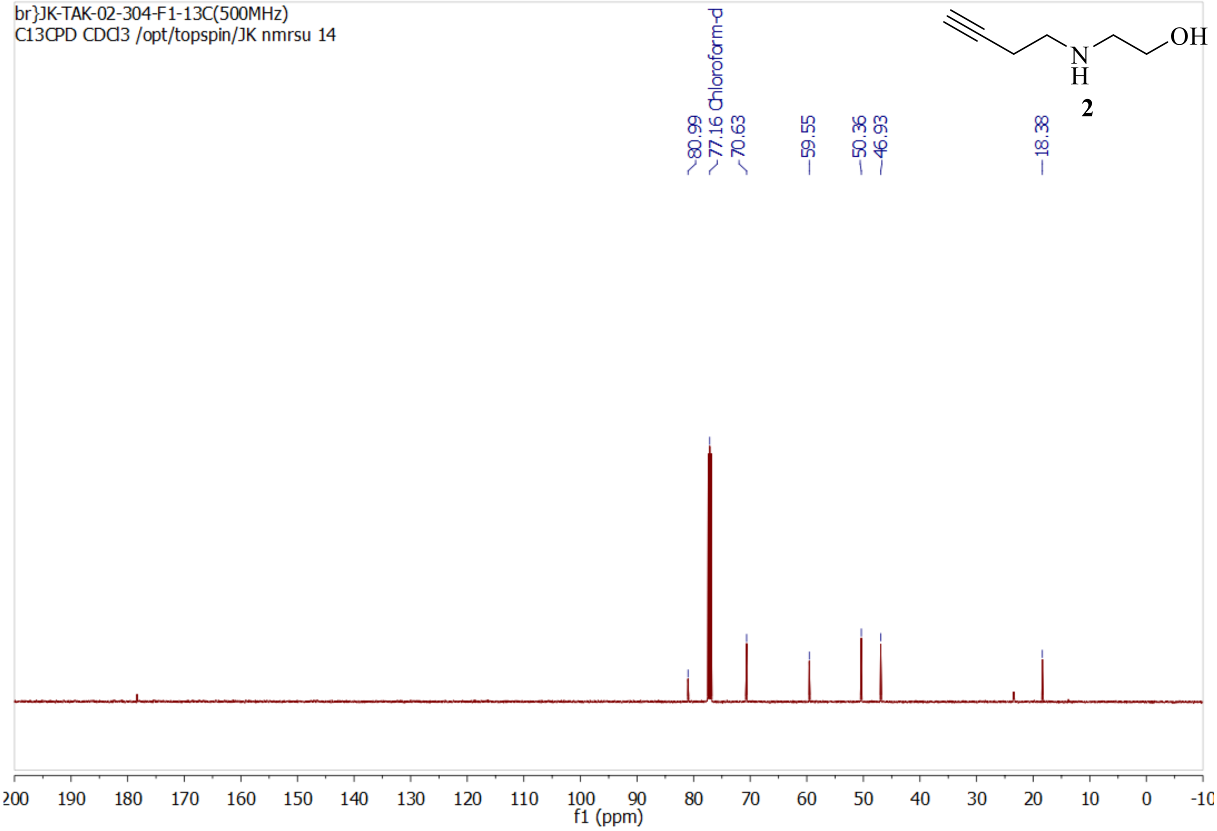
br}JK-TAK-02-304-F1-13C(500MHz)
C13CPD CDCl3 /opt/topspin/JK nmrsu 14
80.99
77.16 Chloroform-d
70.63
59.55
50.36
46.93
18.38
f1 (ppm)
N
H
OH
2

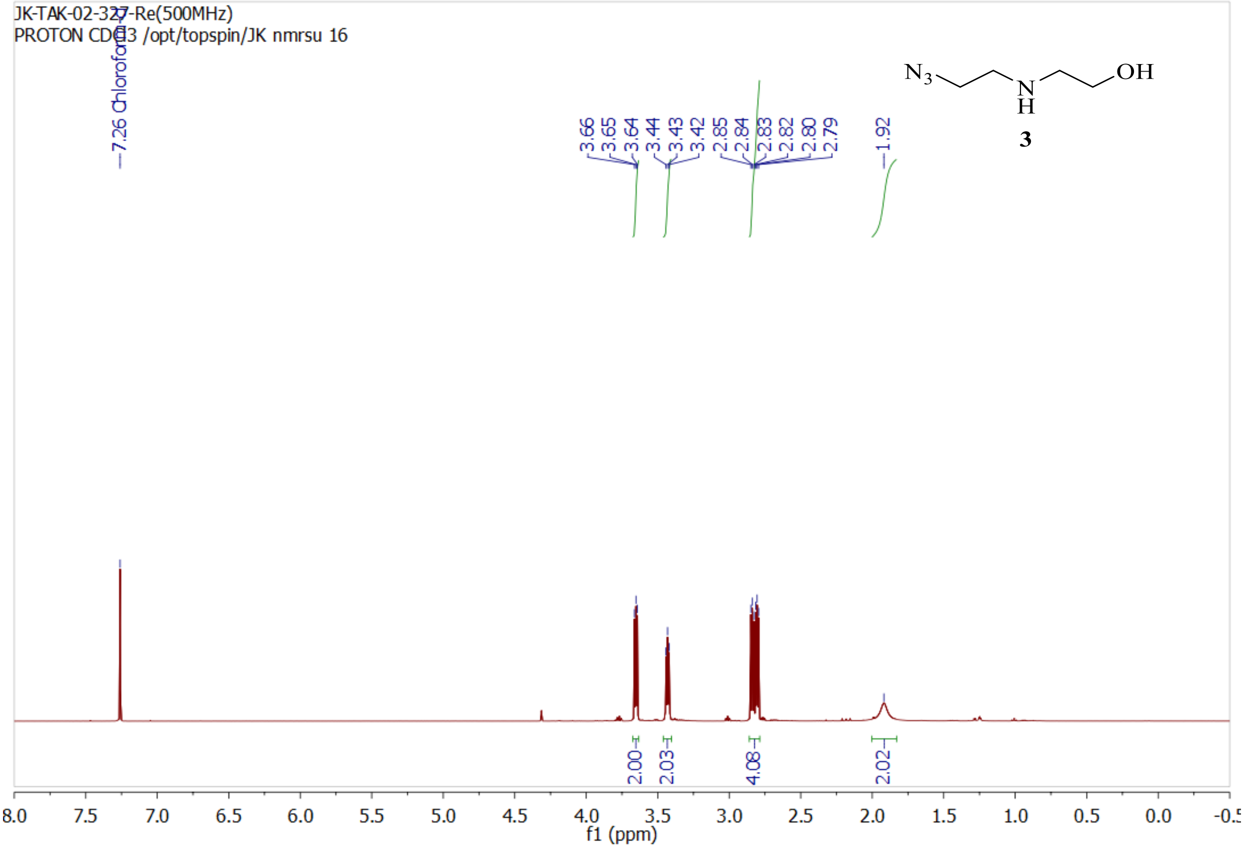
JK-TAK-02-327-Re(500MHz)
PROTON CDCl3 /opt/topspin/JK nmrsu 16
7.26 Chloroform-d
3.66
3.65
3.64
3.44
3.43
3.42
2.85
2.84
2.83
2.82
2.80
2.79
1.92
2.00
2.03
4.08
2.02
8.0 7.5 7.0 6.5 6.0 5.5 5.0 4.5 4.0 3.5 3.0 2.5 2.0 1.5 1.0 0.5 0.0 -0.5
f1 (ppm)
N3 N H OH
3

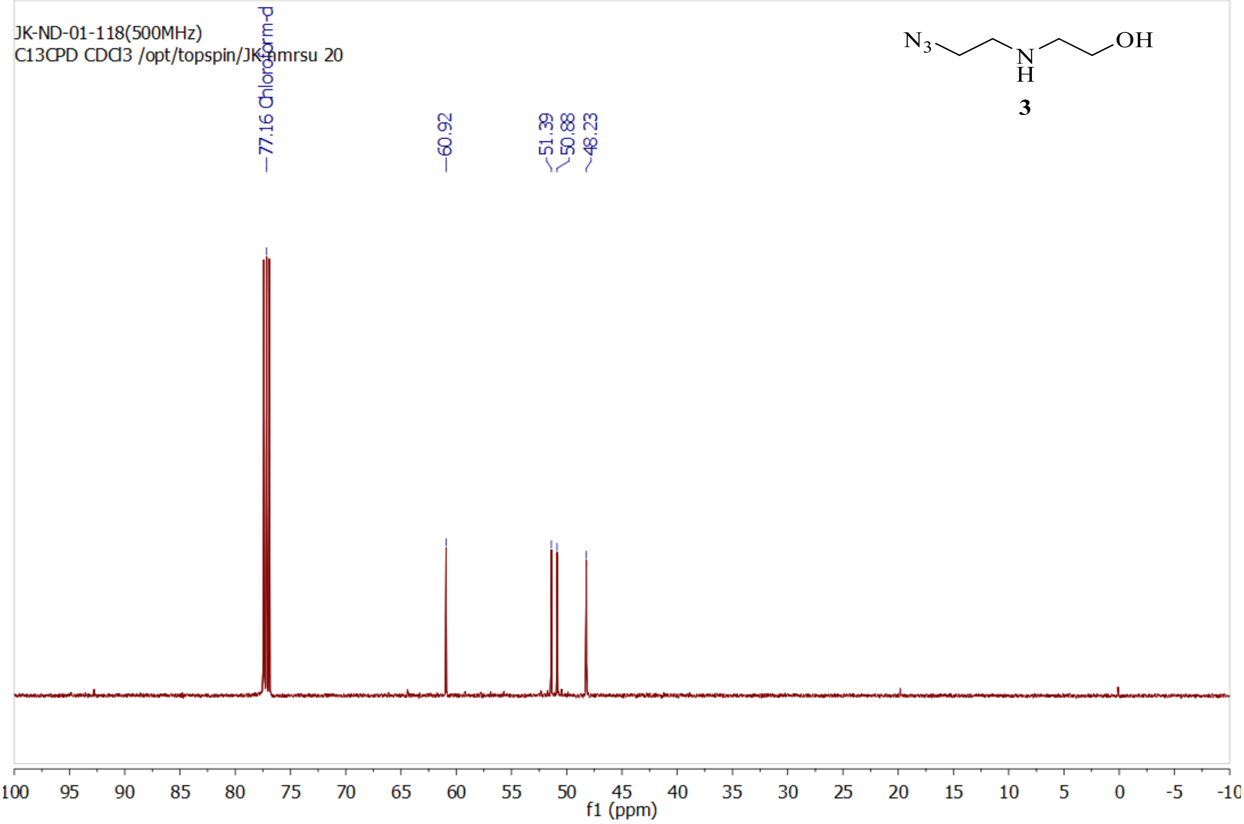
JK-ND-01-118(500MHz)
C13CPD CDCl3 /opt/topspin/JK nmrsu 20
77.16 Chloroform-d
60.92
51.39
50.88
48.23
100 95 90 85 80 75 70 65 60 55 50 45 40 35 30 25 20 15 10 5 0 -5 -10
f1 (ppm)
N3 N H OH
3

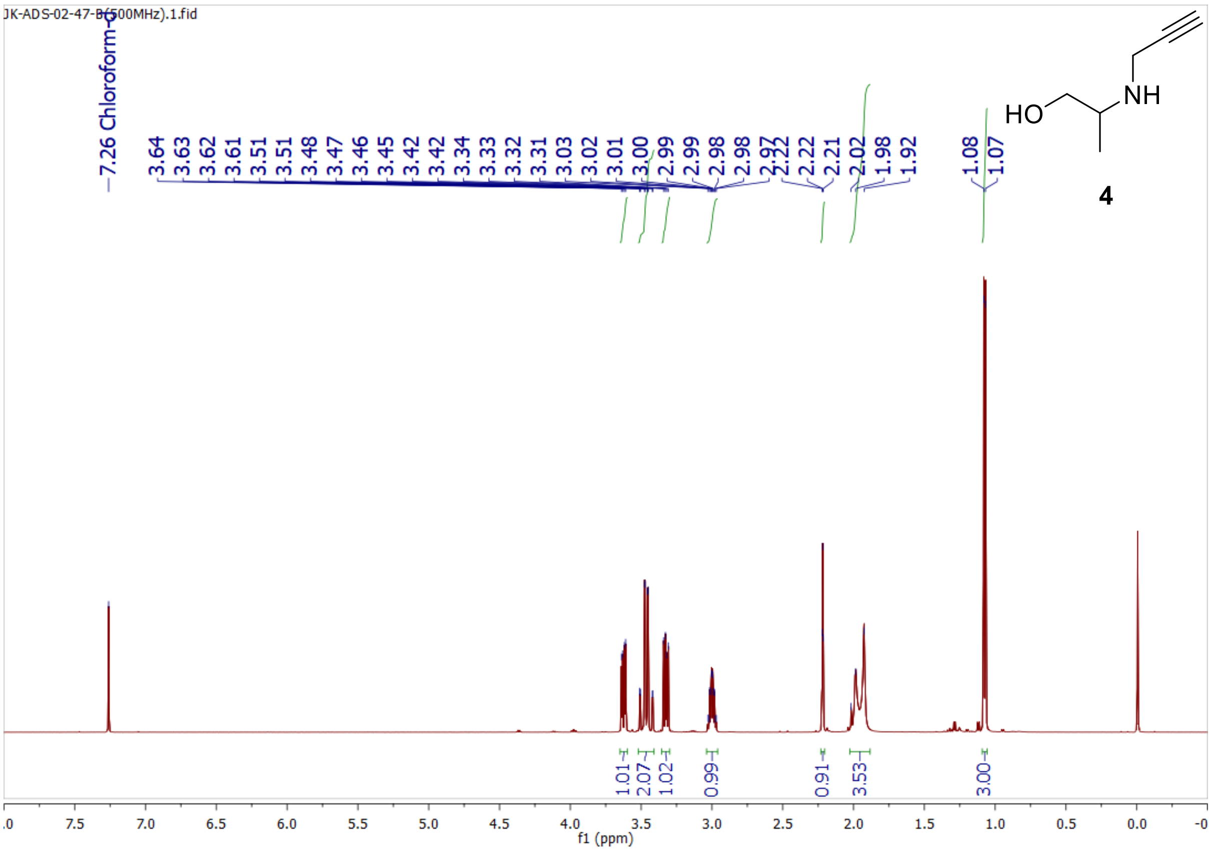
JK-ADS-02-47-B(500MHz).1.fid
7.26 Chloroform-d
3.64
3.63
3.62
3.61
3.51
3.51
3.48
3.47
3.46
3.45
3.42
3.42
3.34
3.33
3.32
3.31
3.03
3.02
3.01
3.00
2.99
2.99
2.98
2.98
2.97
2.22
2.22
2.21
2.02
1.98
1.92
1.08
1.07
HO
NH
4
1.01
2.07
1.02
0.99
0.91
3.53
3.00
f1 (ppm)

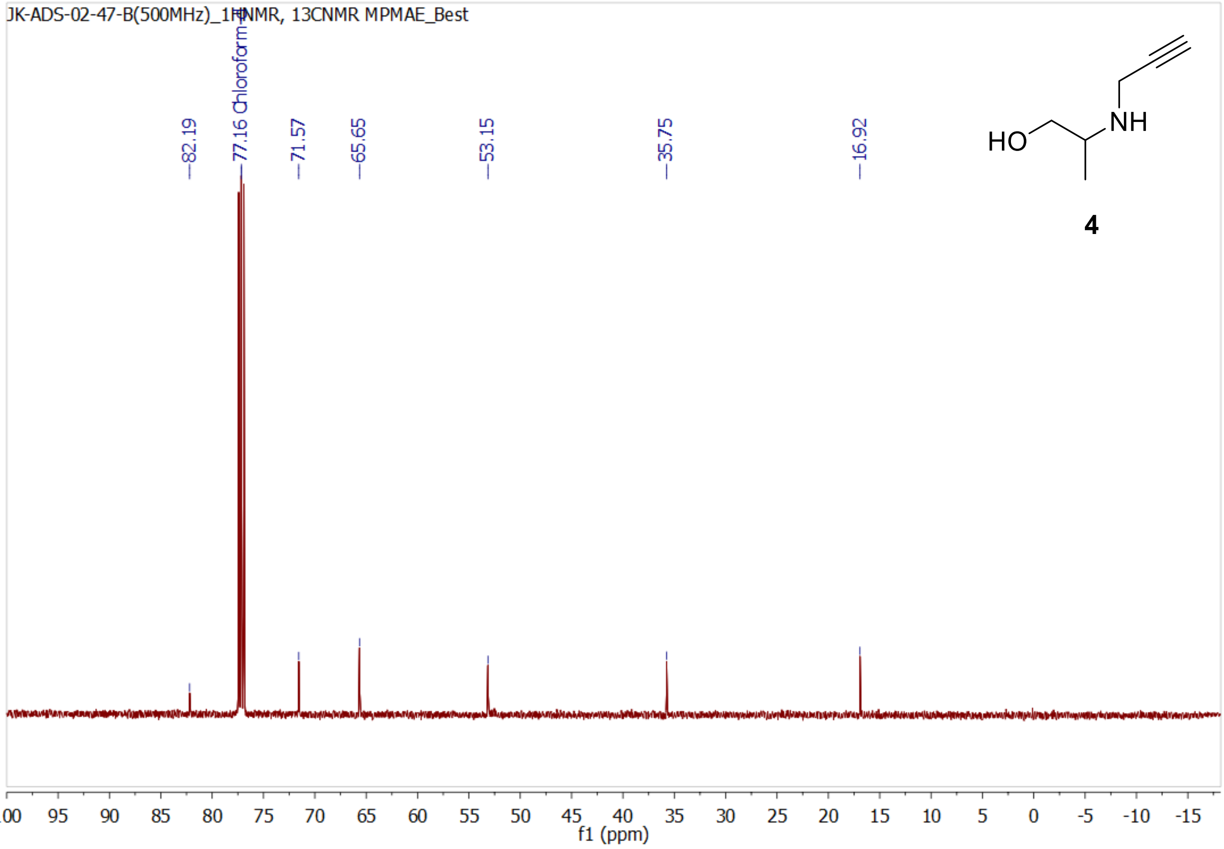
JK-ADS-02-47-B(500MHz)_1HNMR, 13CNMR MPMAE_Best
82.19
77.16 Chloroform-d
71.57
65.65
53.15
35.75
16.92
HO
NH
4
f1 (ppm)

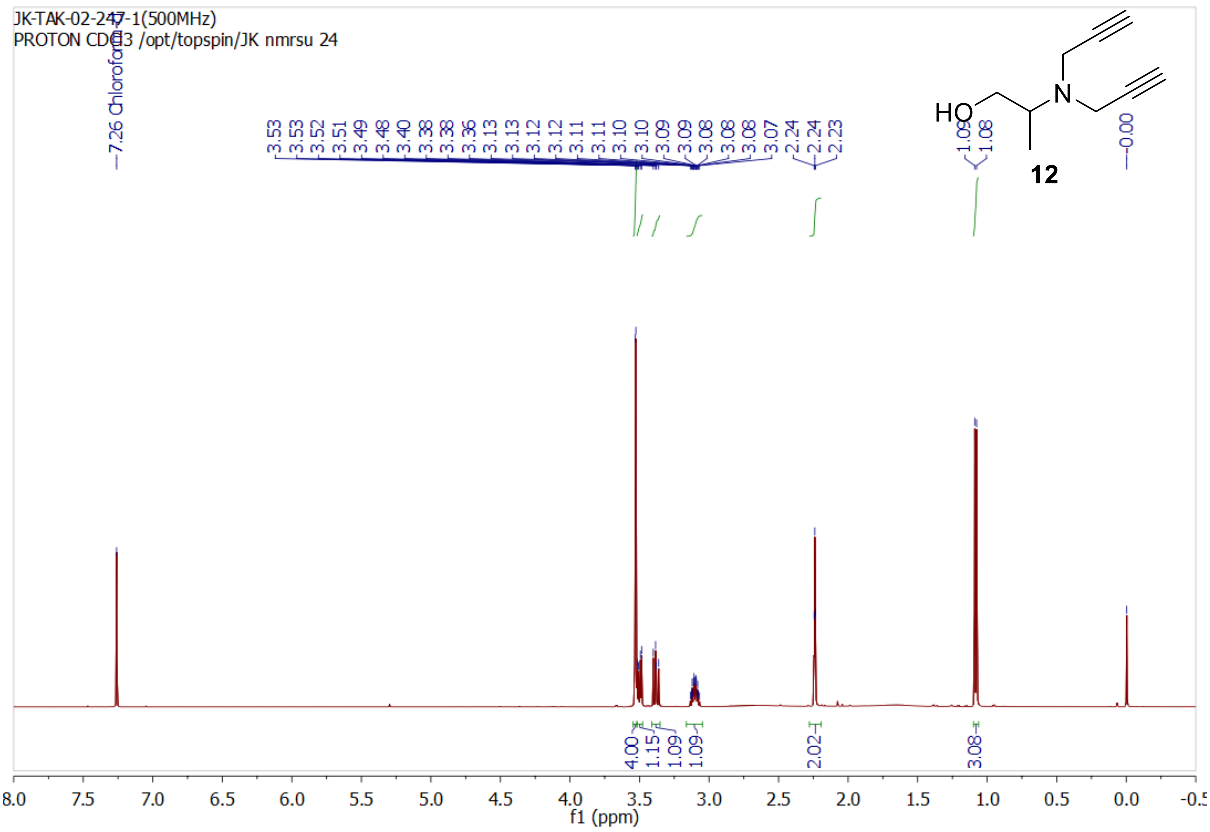
JK-TAK-02-247-1(500MHz)
PROTON CDCl3 /opt/topspin/JK nmrsu 24
7.26 Chloroform-d
3.53 3.53 3.52 3.51 3.49 3.48 3.40 3.38 3.38 3.36 3.13 3.13 3.12 3.12 3.11 3.11 3.10 3.10 3.09 3.09 3.08 3.08 3.08 3.07 2.24 2.24 2.23
1.09 1.08
0.00
HO
N
12
4.00 1.15 1.09 1.09 2.02 3.08
f1 (ppm)

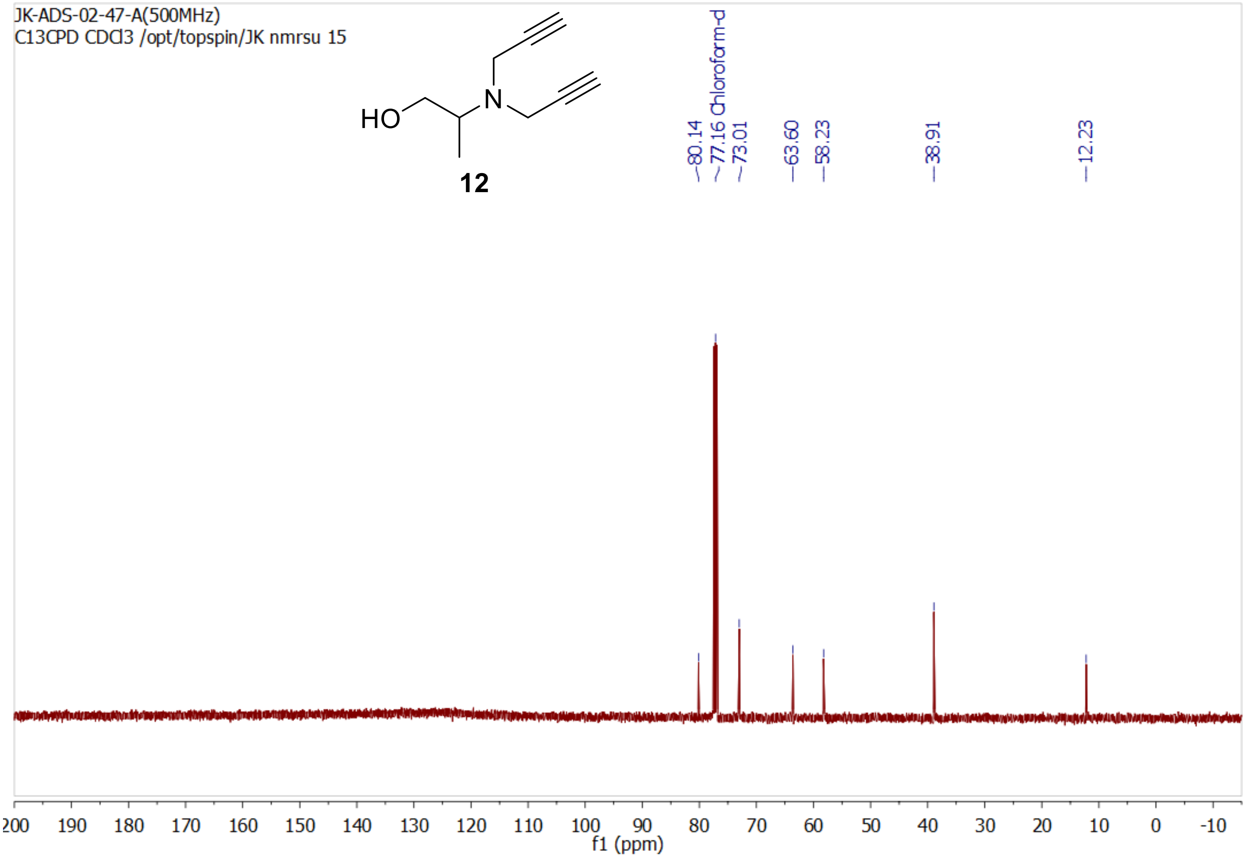
JK-ADS-02-47-A(500MHz)
C13CPD CDCl3 /opt/topspin/JK nmrsu 15
HO
N
12
80.14 77.16 Chloroform-d 73.01 63.60 58.23 38.91 12.23
f1 (ppm)

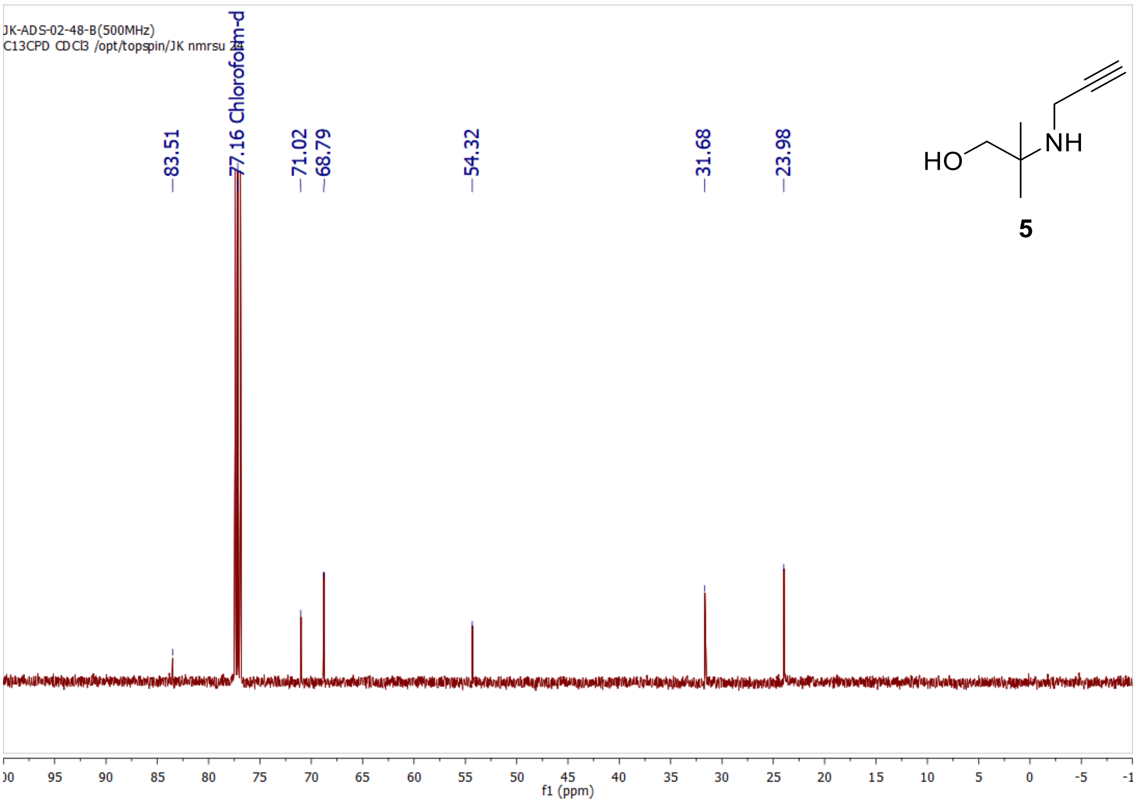
JK-ADS-02-48-B(500MHz)
C13CPD CDCl3 /opt/topspin/JK nmrsu 2
83.51
77.16 Chloroform-d
71.02
68.79
54.32
31.68
23.98
HO
NH
5
f1 (ppm)

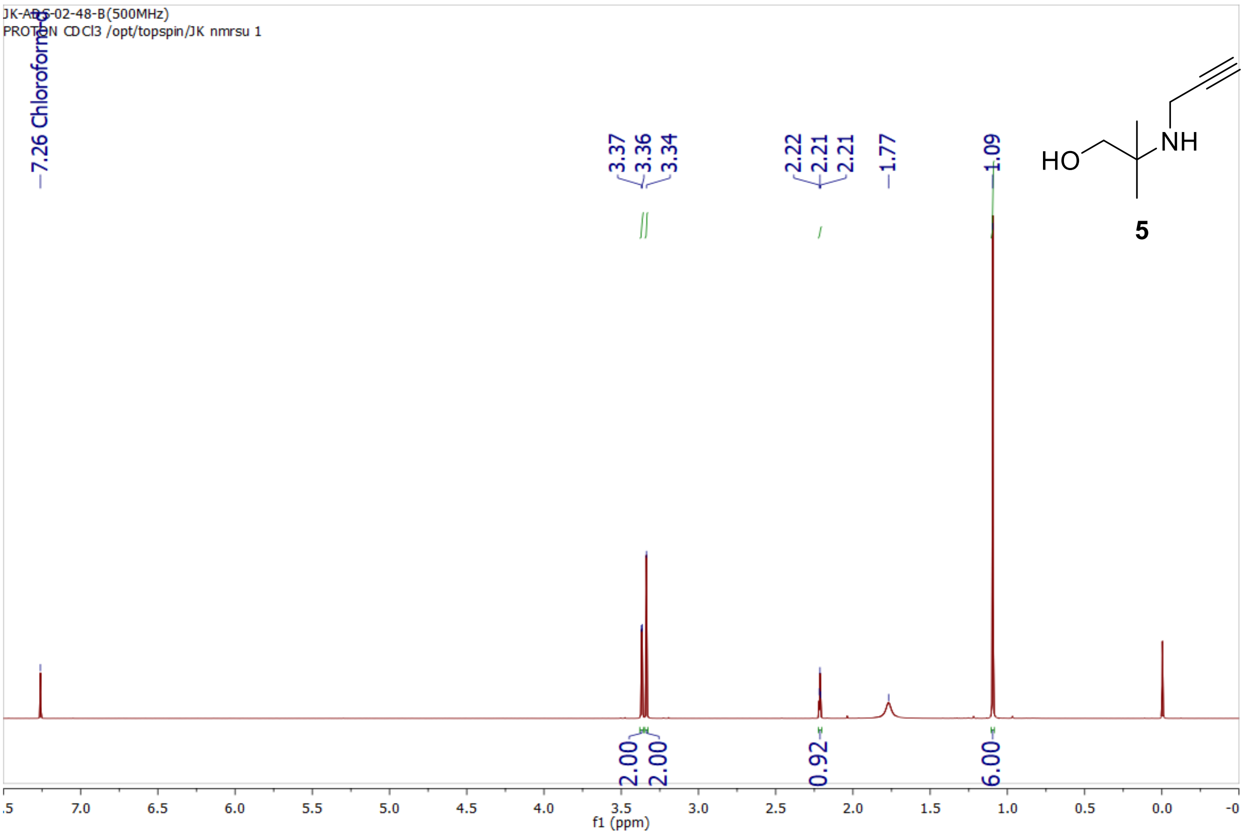
JK-ADS-02-48-B(500MHz)
PROTON CDCl3 /opt/topspin/JK nmrsu 1
7.26 Chloroform-d
3.37
3.36
3.34
2.22
2.21
2.21
1.77
1.09
HO
NH
5
2.00
2.00
0.92
6.00
f1 (ppm)

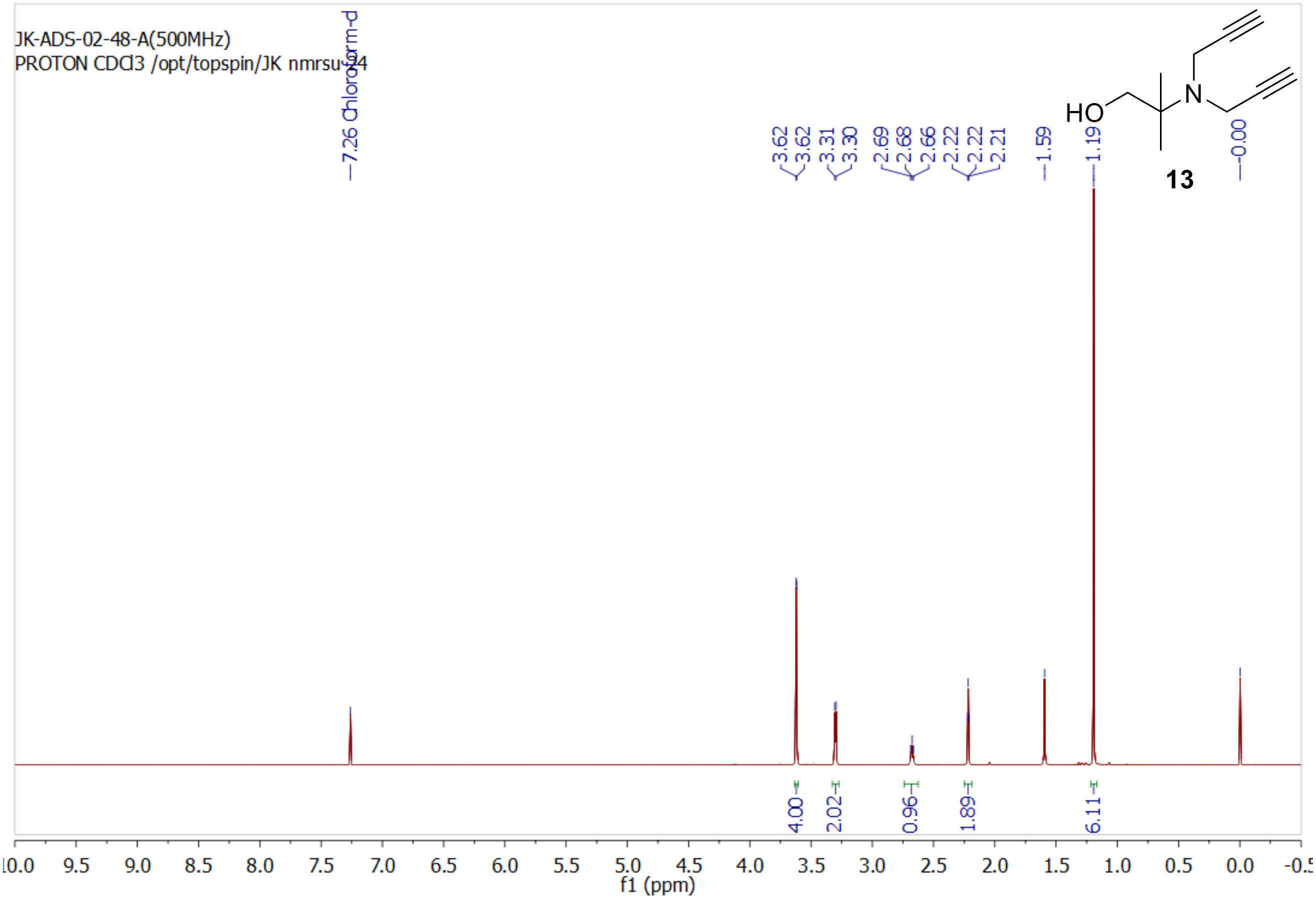
JK-ADS-02-48-A(500MHz)
PROTON CDCl3 /opt/topspin/JK nmrsu 24
7.26 Chloroform-d
3.62
3.62
3.31
3.30
2.69
2.68
2.66
2.22
2.22
2.21
1.59
1.19
0.00
HO
N
13
4.00
2.02
0.96
1.89
6.11
f1 (ppm)

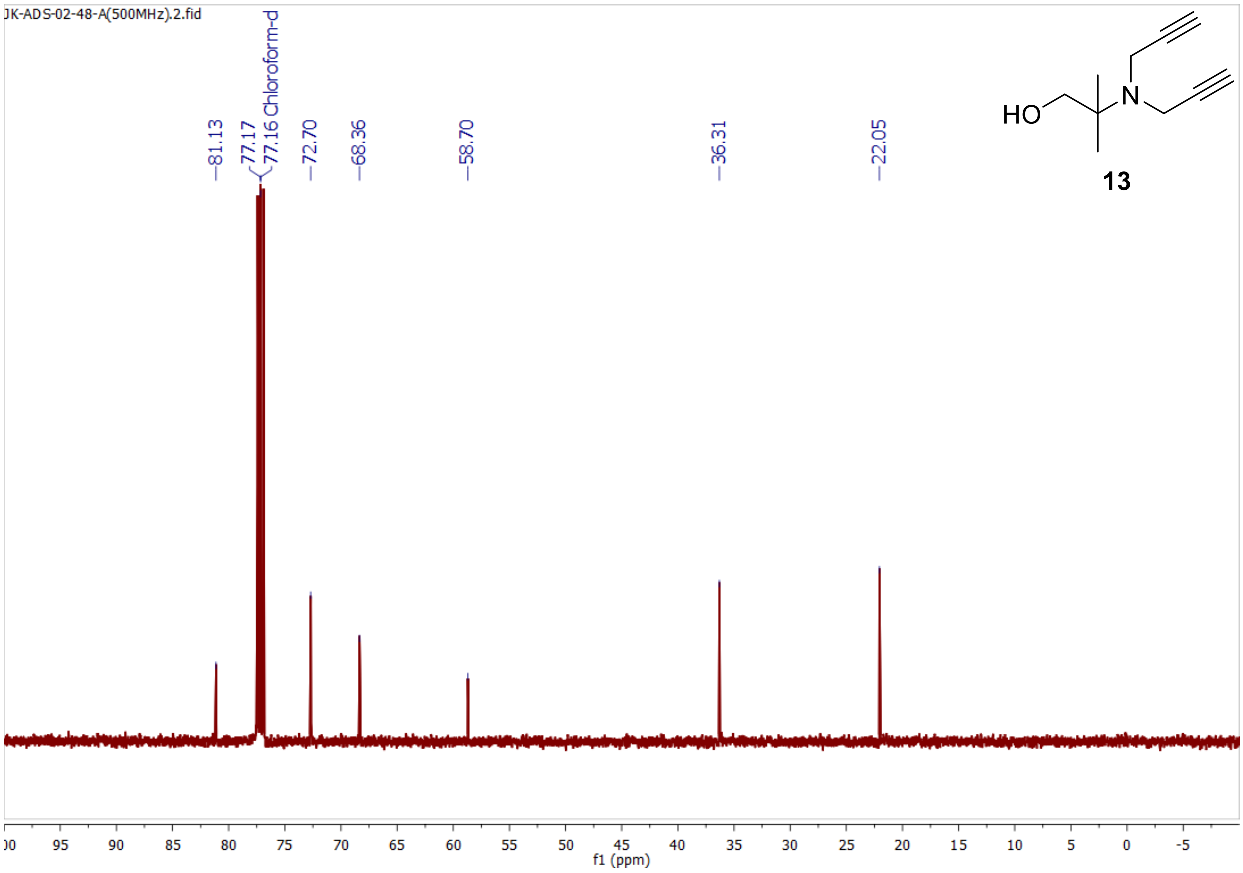
JK-ADS-02-48-A(500MHz).2.fid
81.13
77.17
77.16 Chloroform-d
72.70
68.36
58.70
36.31
22.05
HO
N
13
f1 (ppm)

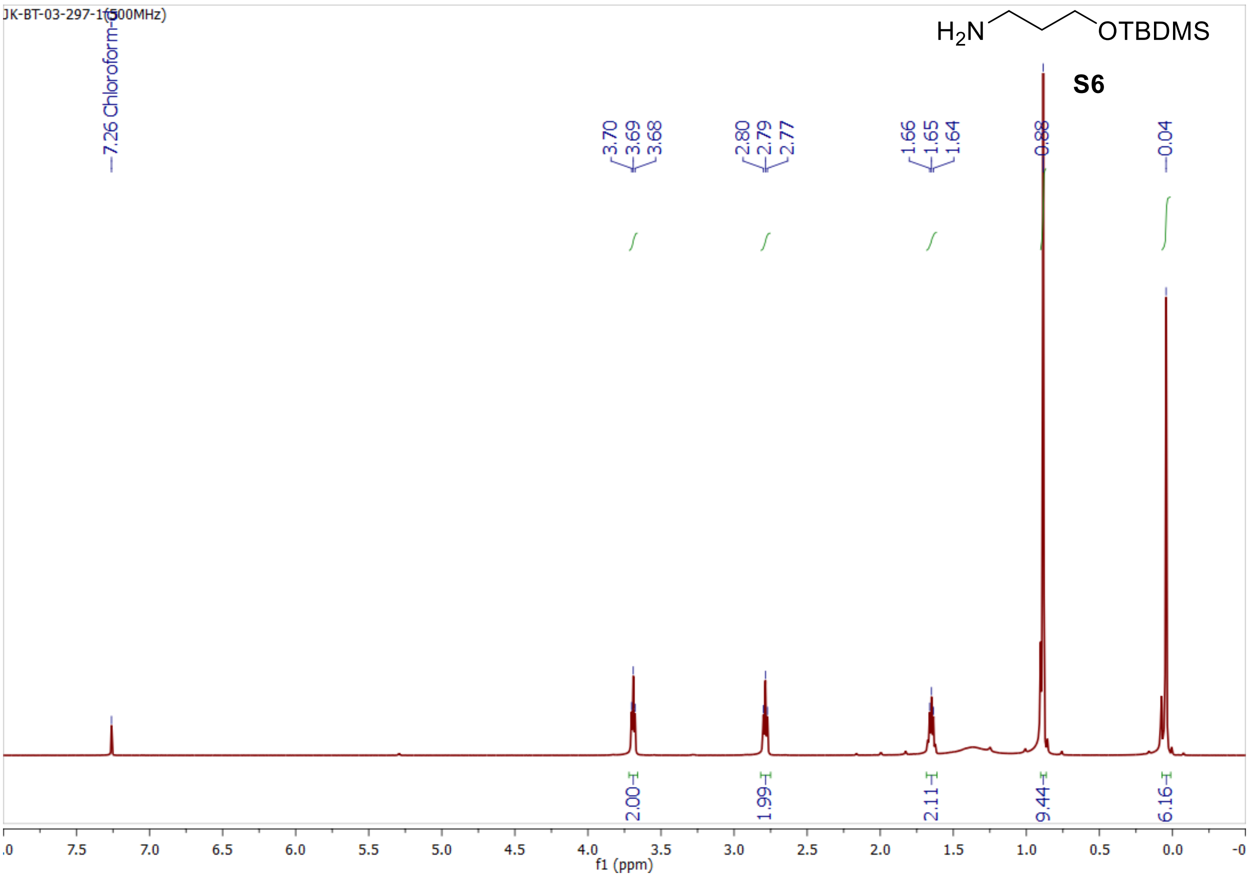

JK-BT-03-297-1(500MHz)
H2N OTBDMS
S6
7.26 Chloroform-d
3.70
3.69
3.68
2.80
2.79
2.77
1.66
1.65
1.64
0.88
0.04
2.00
1.99
2.11
9.44
6.16
f1 (ppm)


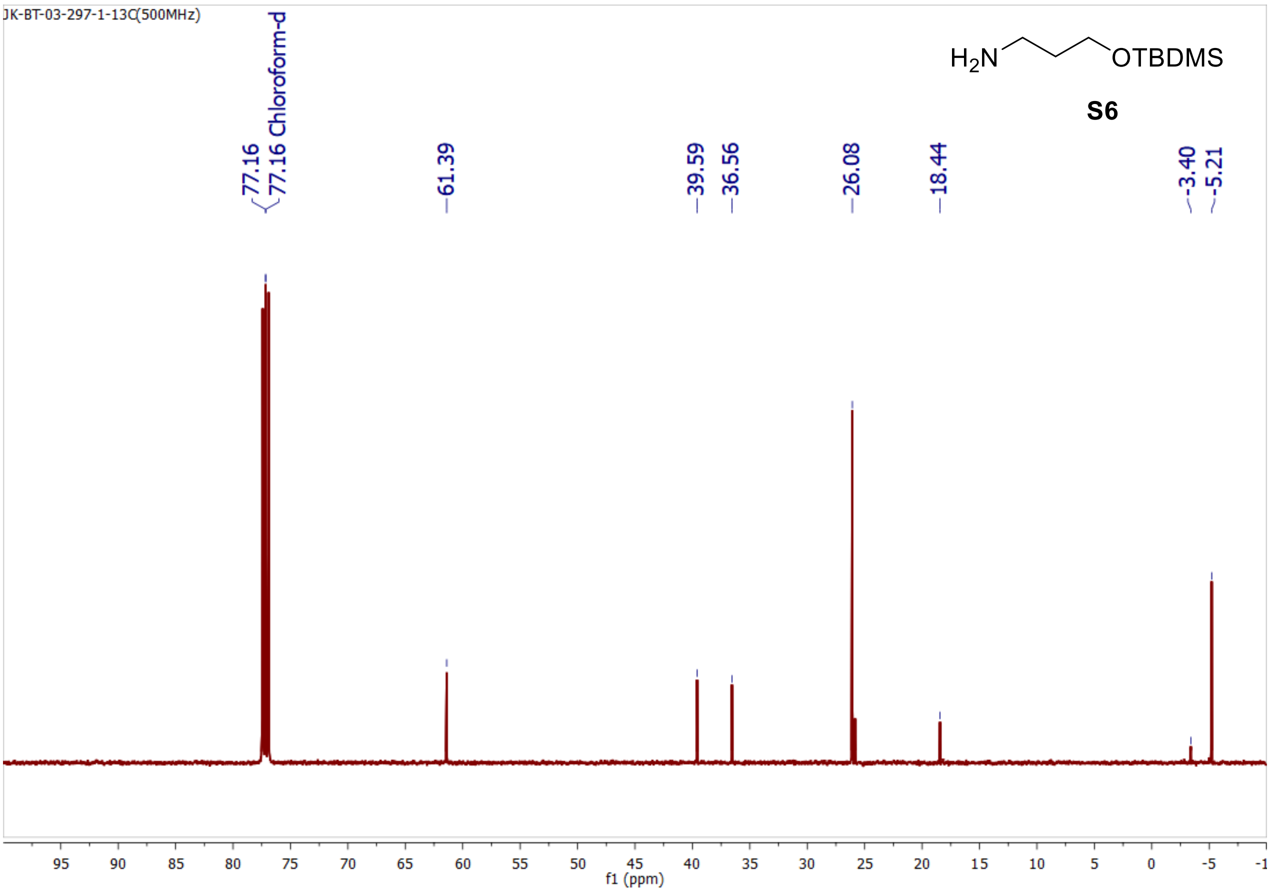

JK-BT-03-297-1-13C(500MHz)
H2N OTBDMS
S6
77.16
77.16 Chloroform-d
61.39
39.59
36.56
26.08
18.44
-3.40
-5.21
f1 (ppm)

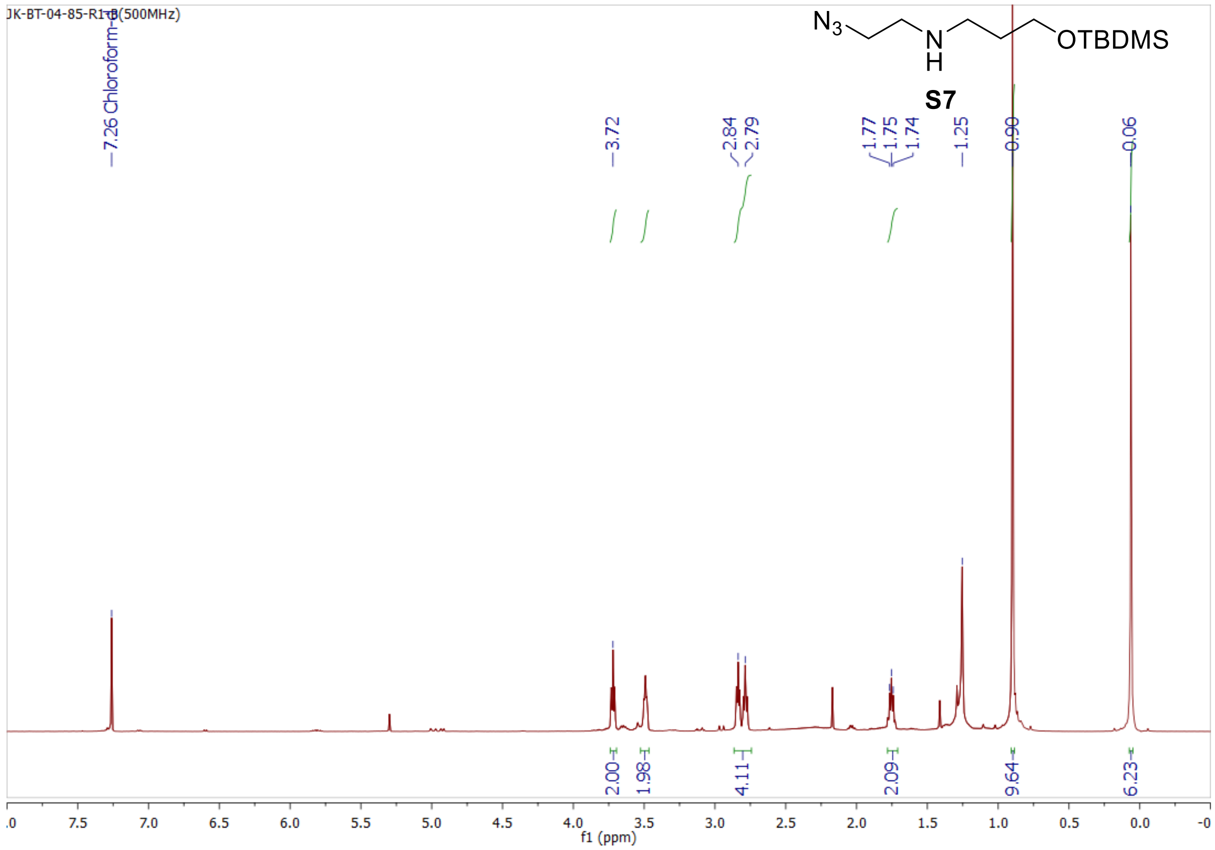
JK-BT-04-85-R1-B(500MHz)
N3 N H OTBDMS
S7
7.26 Chloroform-d
3.72
2.84
2.79
1.77
1.75
1.74
1.25
0.90
0.06
2.00
1.98
4.11
2.09
9.64
6.23
f1 (ppm)

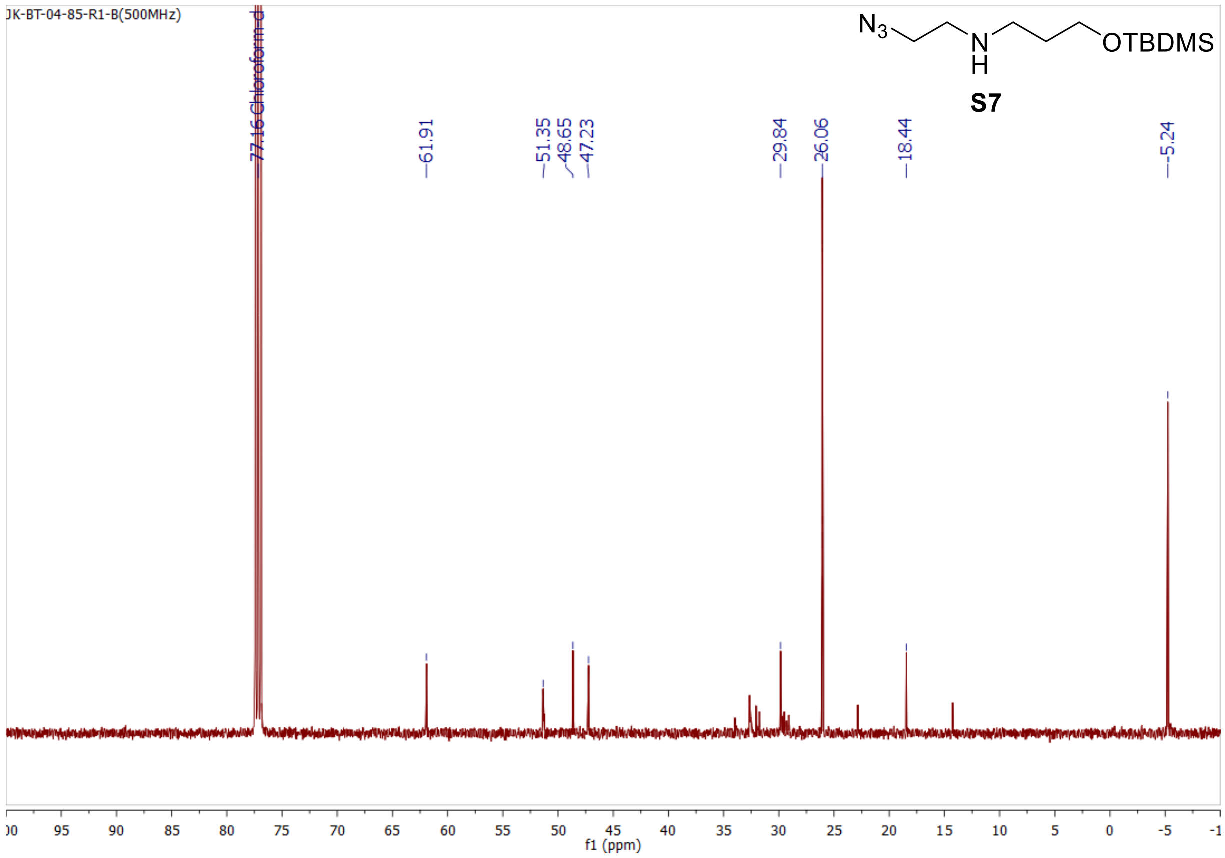
JK-BT-04-85-R1-B(500MHz)
N3 N H OTBDMS
S7
77.16 Chloroform-d
61.91
51.35
48.65
47.23
29.84
26.06
18.44
-5.24
f1 (ppm)

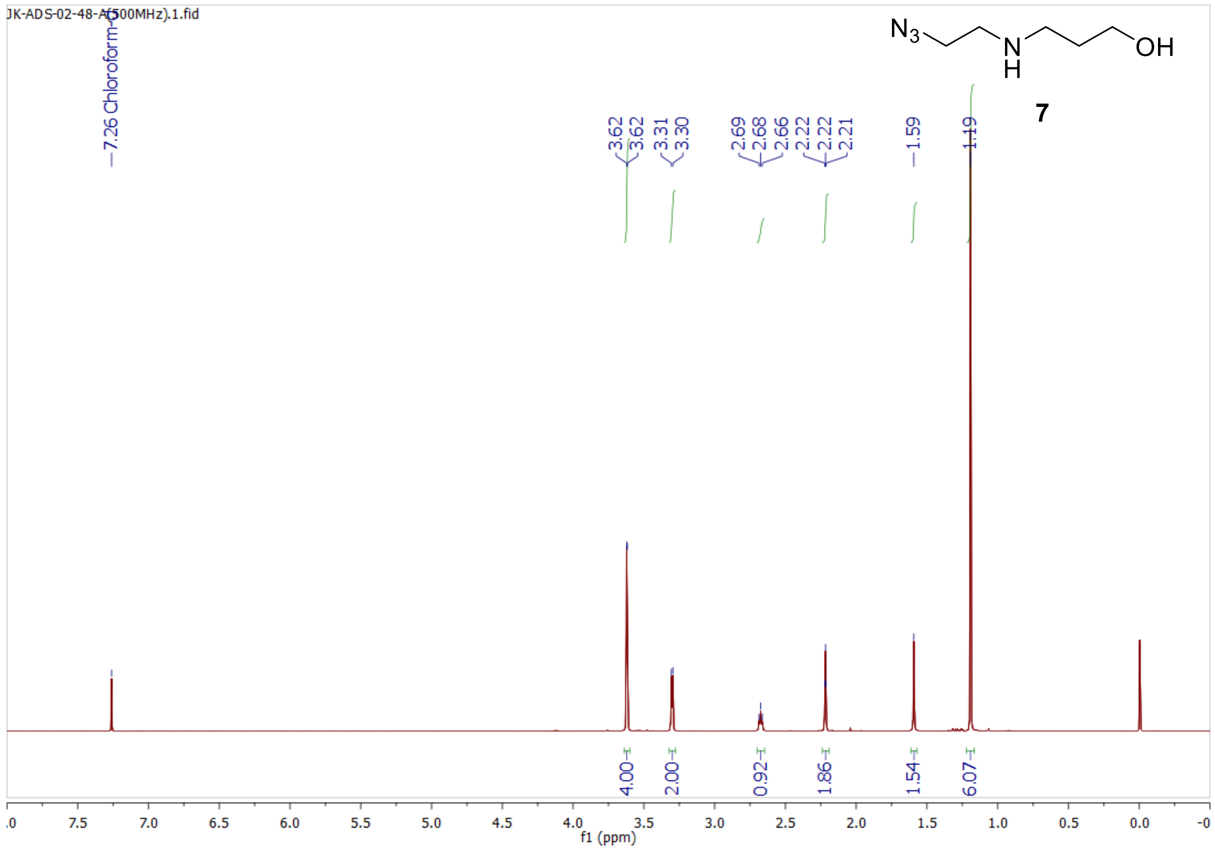
JK-ADS-02-48-A(500MHz).1.fid
7.26 Chloroform-d
3.62
3.62
3.31
3.30
2.69
2.68
2.66
2.22
2.22
2.21
1.59
1.19
N3
N
H
OH
7
4.00
2.00
0.92
1.86
1.54
6.07
f1 (ppm)

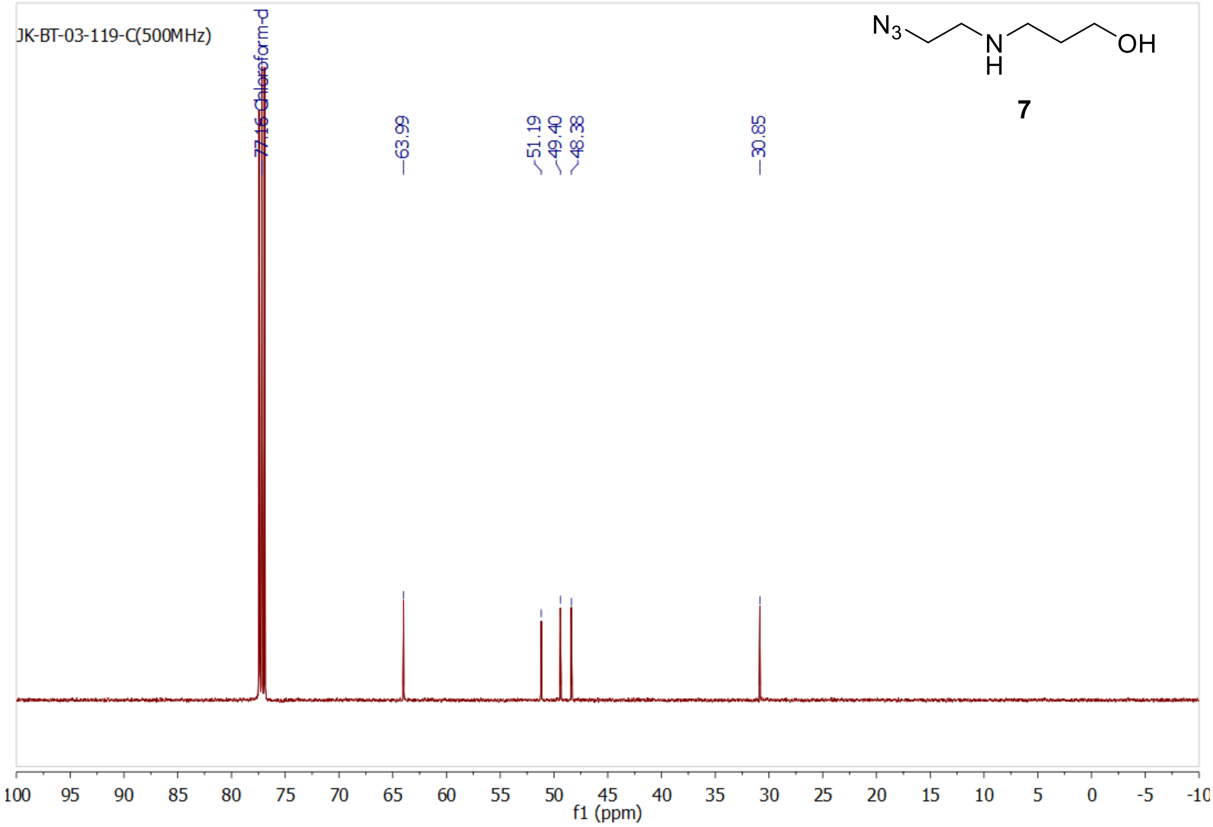
JK-BT-03-119-C(500MHz)
77.16 Chloroform-d
63.99
51.19
49.40
48.38
30.85
N3
N
H
OH
7
f1 (ppm)

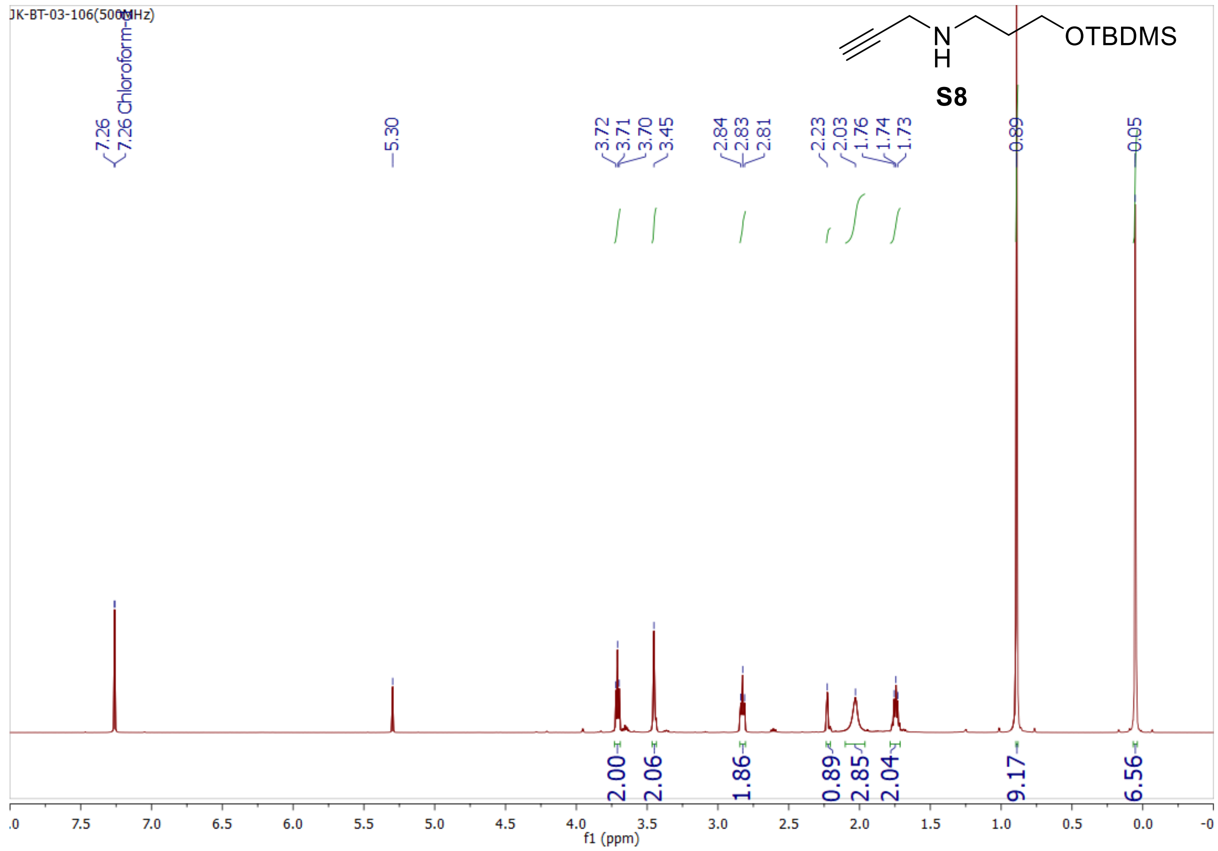
JK-BT-03-106(500MHz)
S8
OTBDMS
N
H
7.26
7.26 Chloroform-d
5.30
3.72
3.71
3.70
3.45
2.84
2.83
2.81
2.23
2.03
1.76
1.74
1.73
0.89
0.05
2.00
2.06
1.86
0.89
2.85
2.04
9.17
6.56
f1 (ppm)

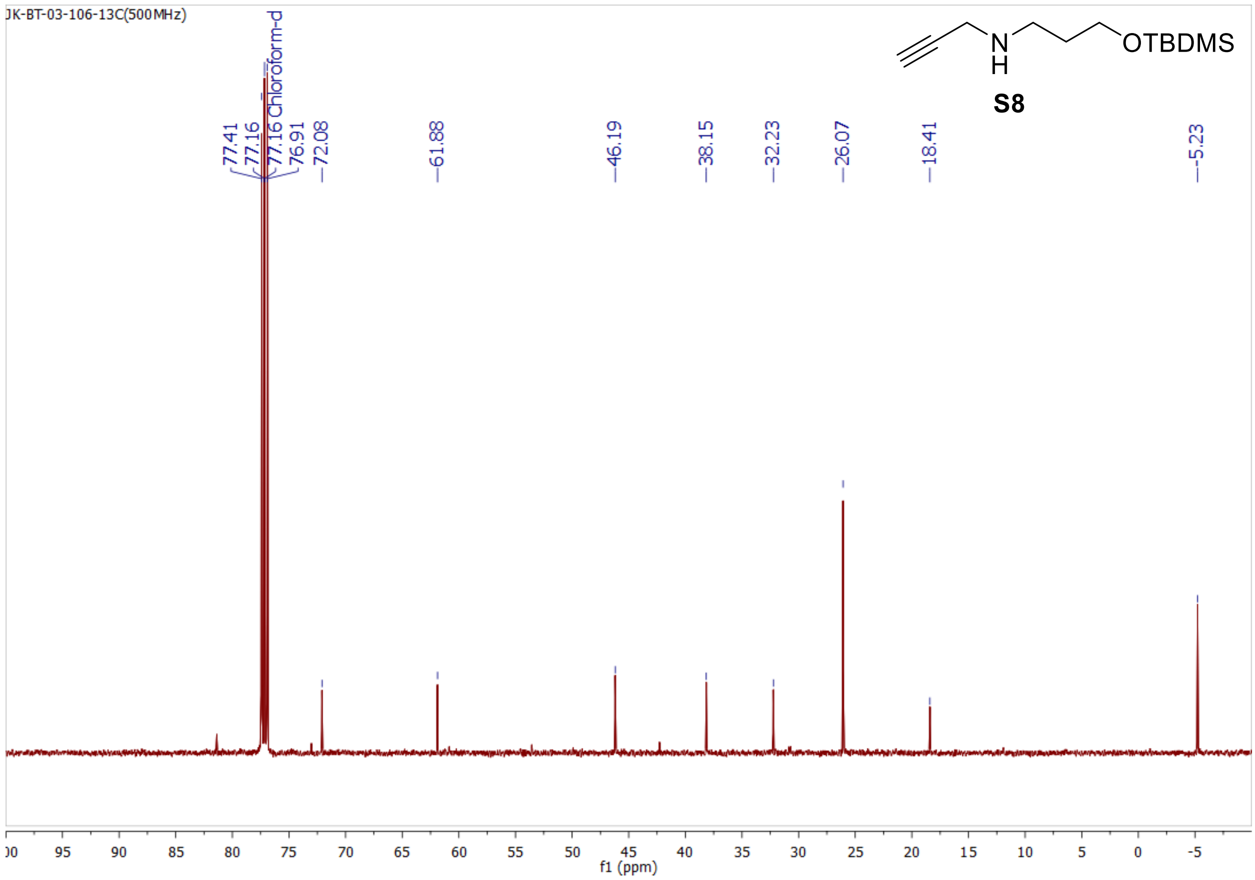
JK-BT-03-106-13C(500MHz)
S8
OTBDMS
N
H
77.41
77.16
77.16 Chloroform-d
76.91
72.08
61.88
46.19
38.15
32.23
26.07
18.41
-5.23
f1 (ppm)

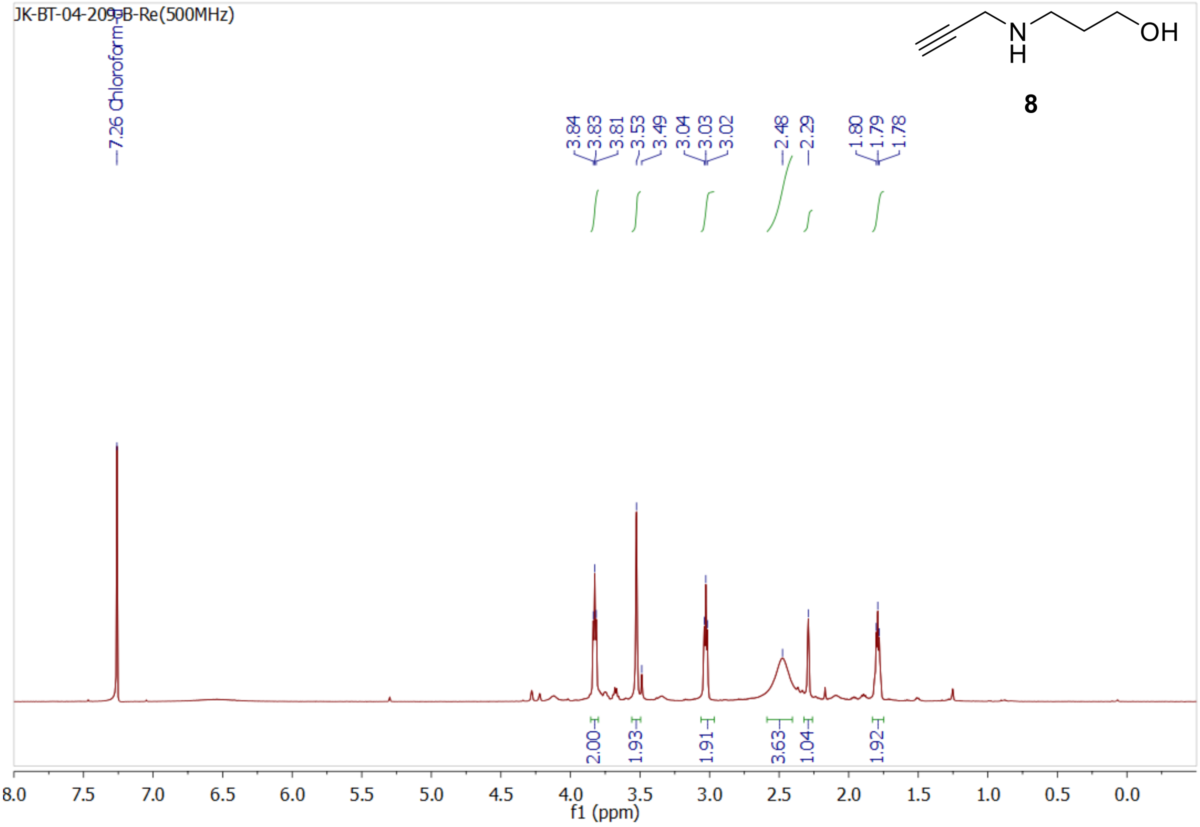
JK-BT-04-209-B-Re(500MHz)
7.26 Chloroform-d
3.84
3.83
3.81
3.53
3.49
3.04
3.03
3.02
2.48
2.29
1.80
1.79
1.78
OH
N
H
8
2.00
1.93
1.91
3.63
1.04
1.92
8.0 7.5 7.0 6.5 6.0 5.5 5.0 4.5 4.0 3.5 3.0 2.5 2.0 1.5 1.0 0.5 0.0
f1 (ppm)

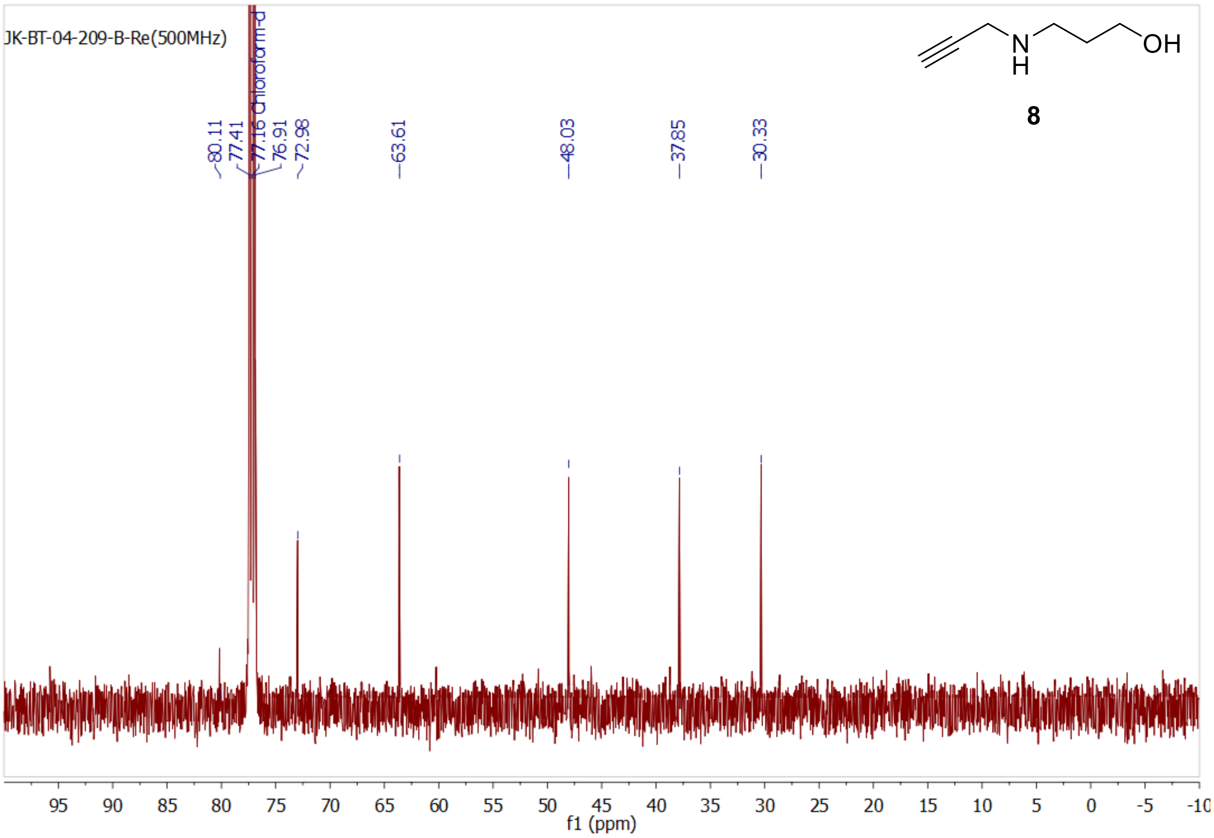
JK-BT-04-209-B-Re(500MHz)
Chloroform-d
80.11
77.41
77.16
76.91
72.98
63.61
48.03
37.85
30.33
OH
N
H
8
95 90 85 80 75 70 65 60 55 50 45 40 35 30 25 20 15 10 5 0 -5 -10
f1 (ppm)

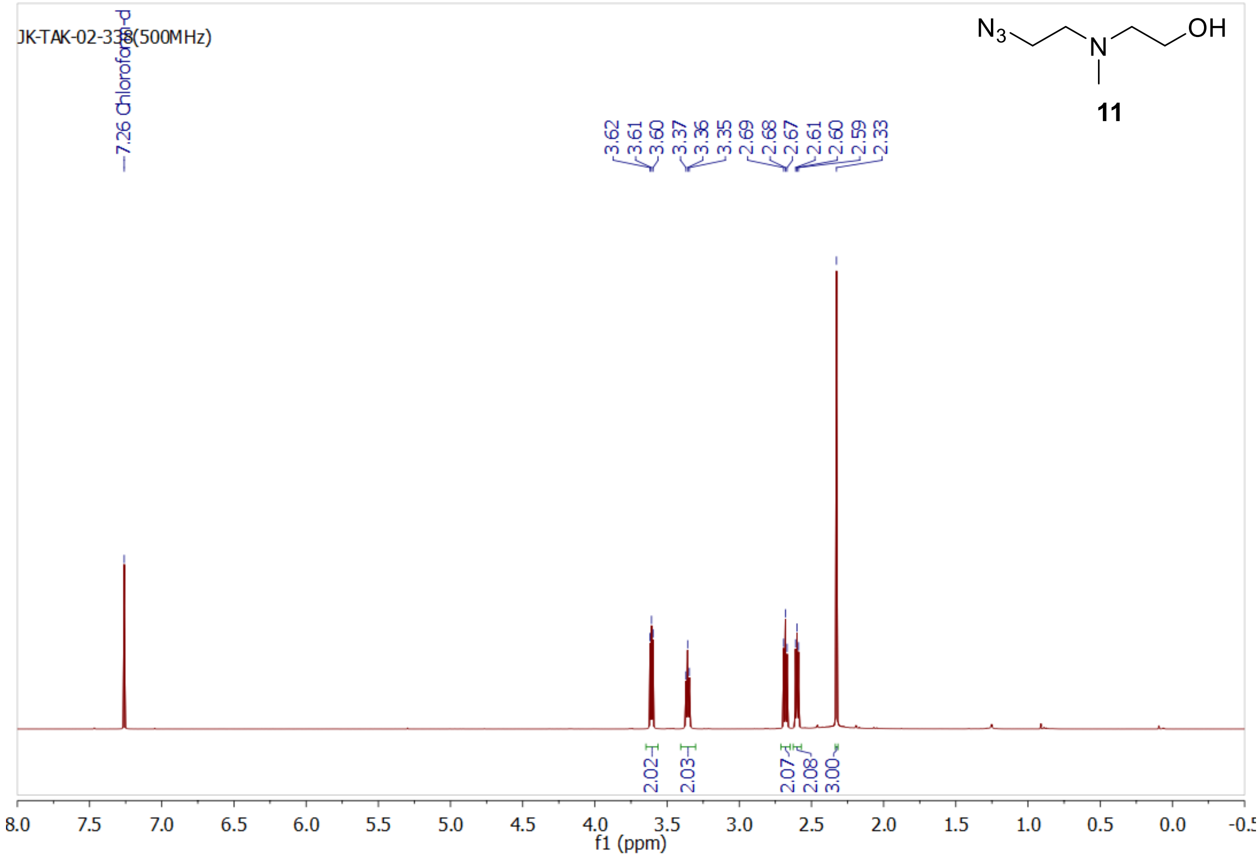
JK-TAK-02-338(500MHz)
7.26 Chloroform-d
3.62
3.61
3.60
3.37
3.36
3.35
2.69
2.68
2.67
2.61
2.60
2.59
2.33
2.02
2.03
2.07
2.08
3.00
N3
N
OH
11
f1 (ppm)

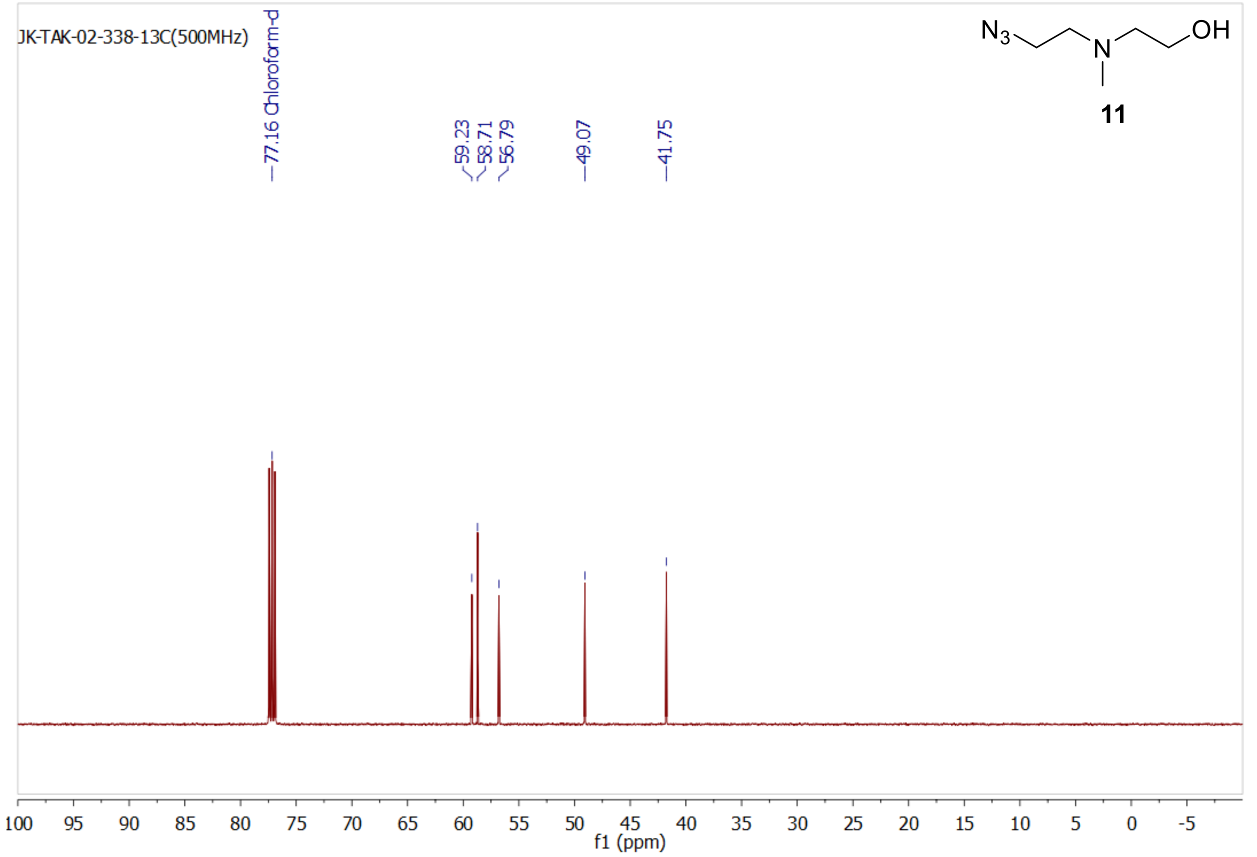
JK-TAK-02-338-13C(500MHz)
77.16 Chloroform-d
59.23
58.71
56.79
49.07
41.75
N3
N
OH
11
f1 (ppm)

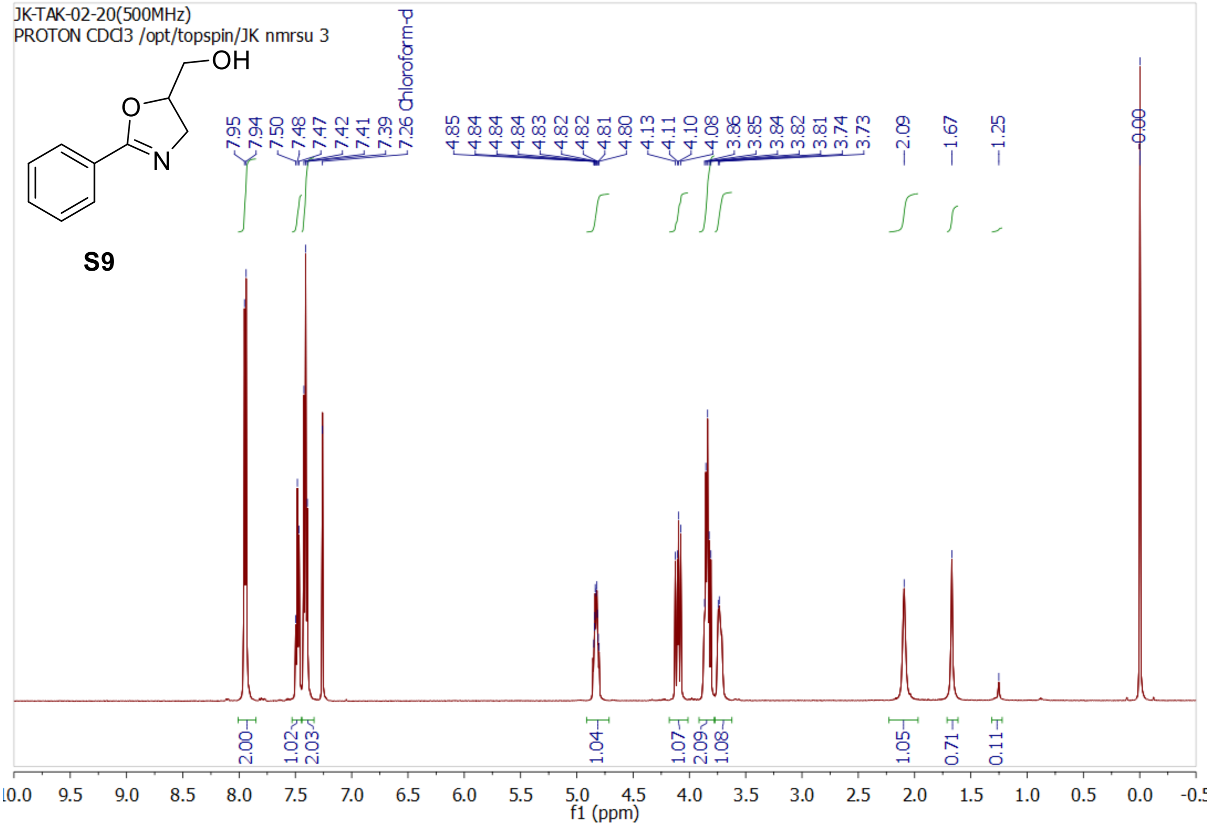
JK-TAK-02-20(500MHz)
PROTON CDCl3 /opt/topspin/JK nmrsu 3
OH
O
N
S9
Chloroform-d
f1 (ppm)

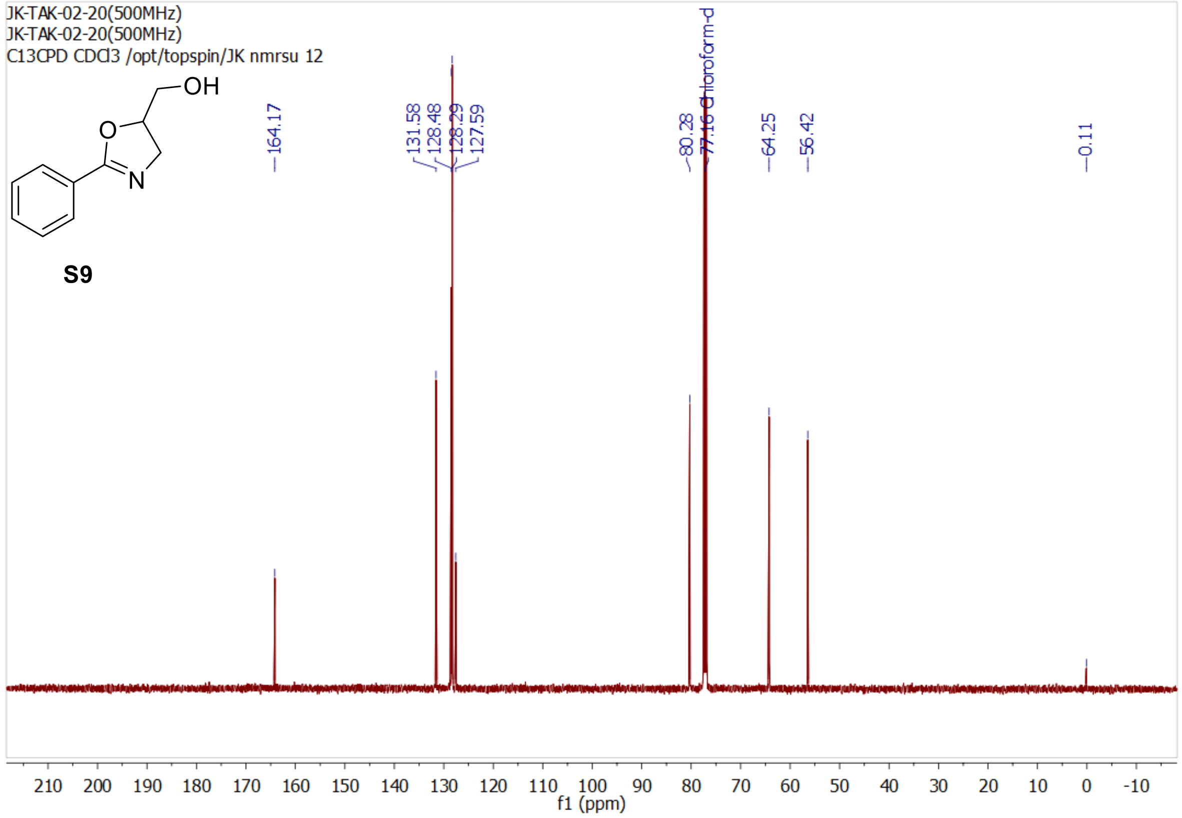
JK-TAK-02-20(500MHz)
JK-TAK-02-20(500MHz)
C13CPD CDCl3 /opt/topspin/JK nmrsu 12
OH
O
N
S9
164.17
131.58
128.48
128.29
127.59
80.28
77.16 Chloroform-d
64.25
56.42
0.11
f1 (ppm)

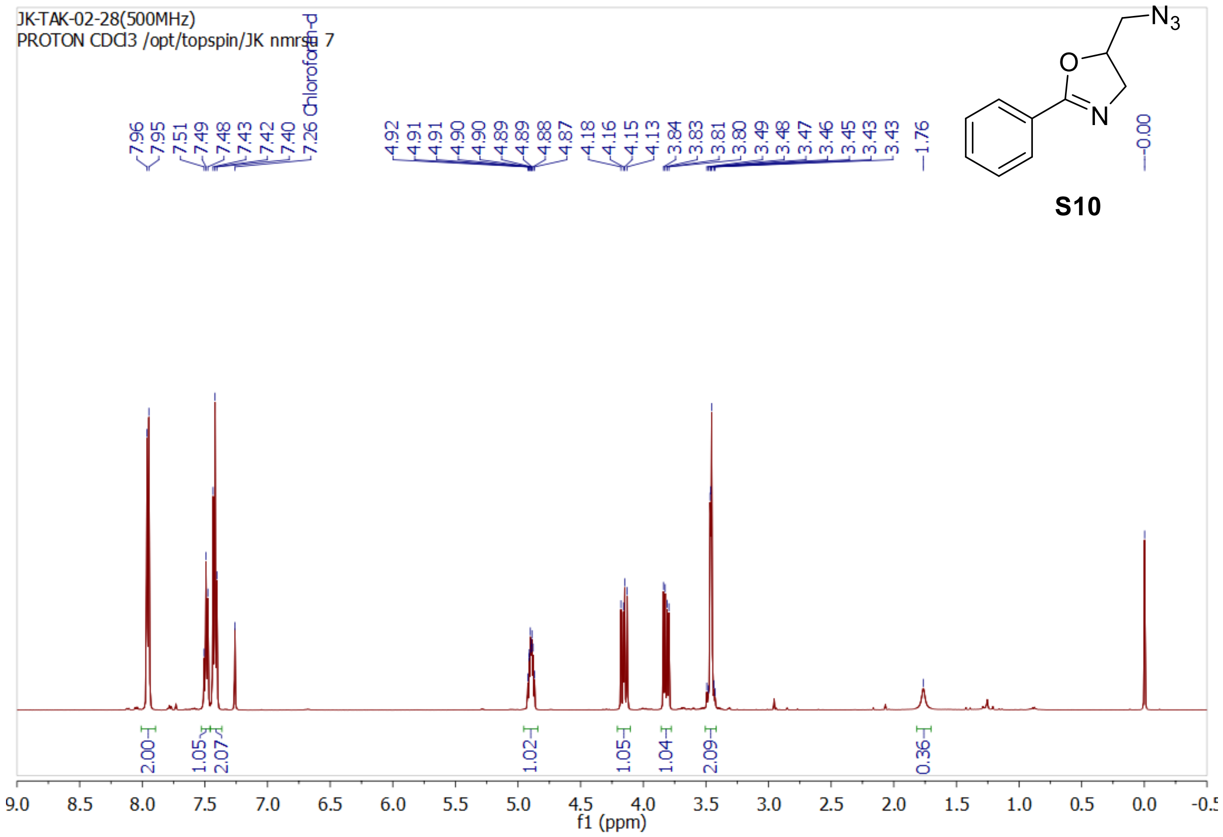
JK-TAK-02-28(500MHz)
PROTON CDCl3 /opt/topspin/JK nmrsu 7
7.96
7.95
7.51
7.49
7.48
7.43
7.42
7.40
7.26 Chloroform-d
4.92
4.91
4.91
4.90
4.90
4.89
4.89
4.88
4.87
4.18
4.16
4.15
4.13
3.84
3.83
3.81
3.80
3.49
3.48
3.47
3.46
3.45
3.43
3.43
1.76
0.00
N3
O
N
S10
2.00
1.05
2.07
1.02
1.05
1.04
2.09
0.36
f1 (ppm)

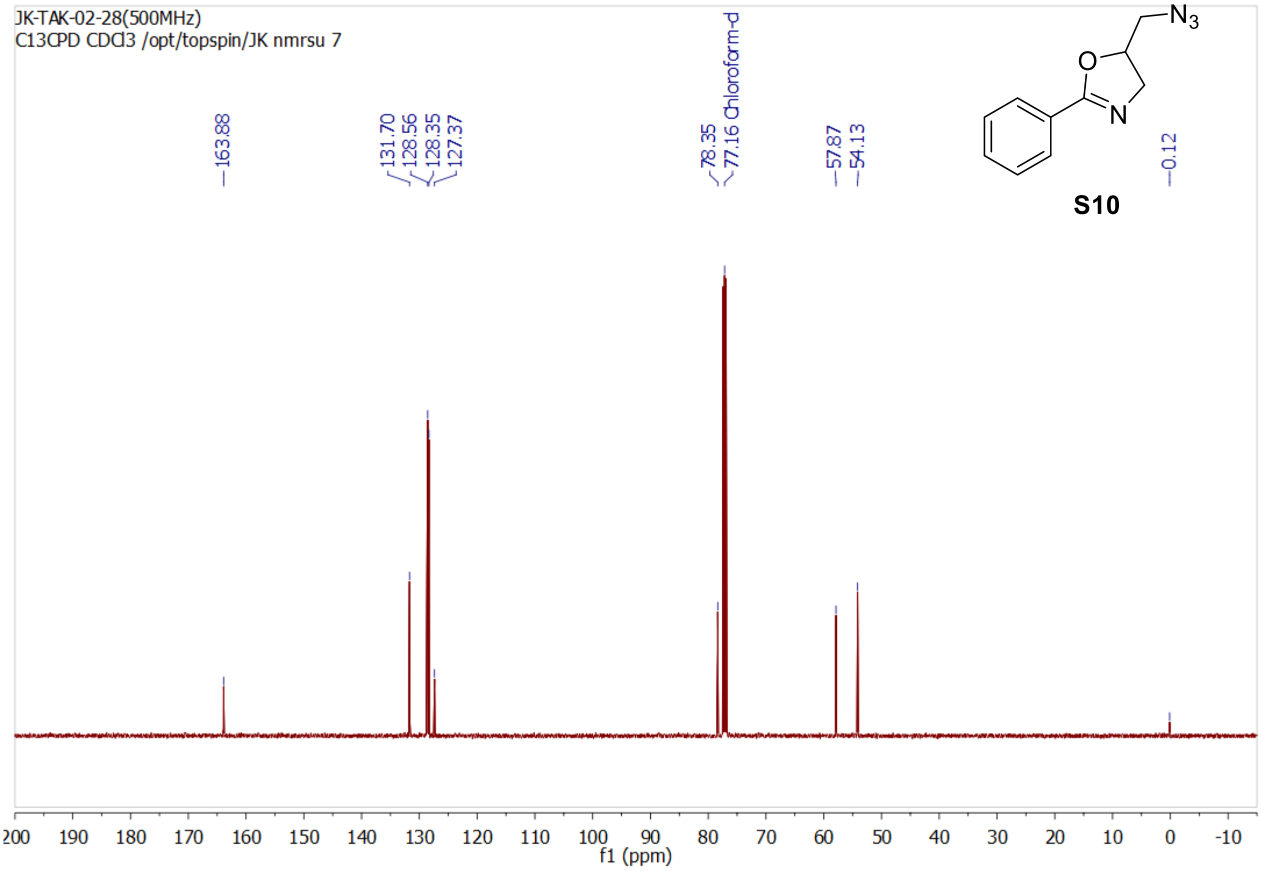
JK-TAK-02-28(500MHz)
C13CPD CDCl3 /opt/topspin/JK nmrsu 7
163.88
131.70
128.56
128.35
127.37
78.35
77.16 Chloroform-d
57.87
54.13
0.12
N3
O
N
S10
f1 (ppm)

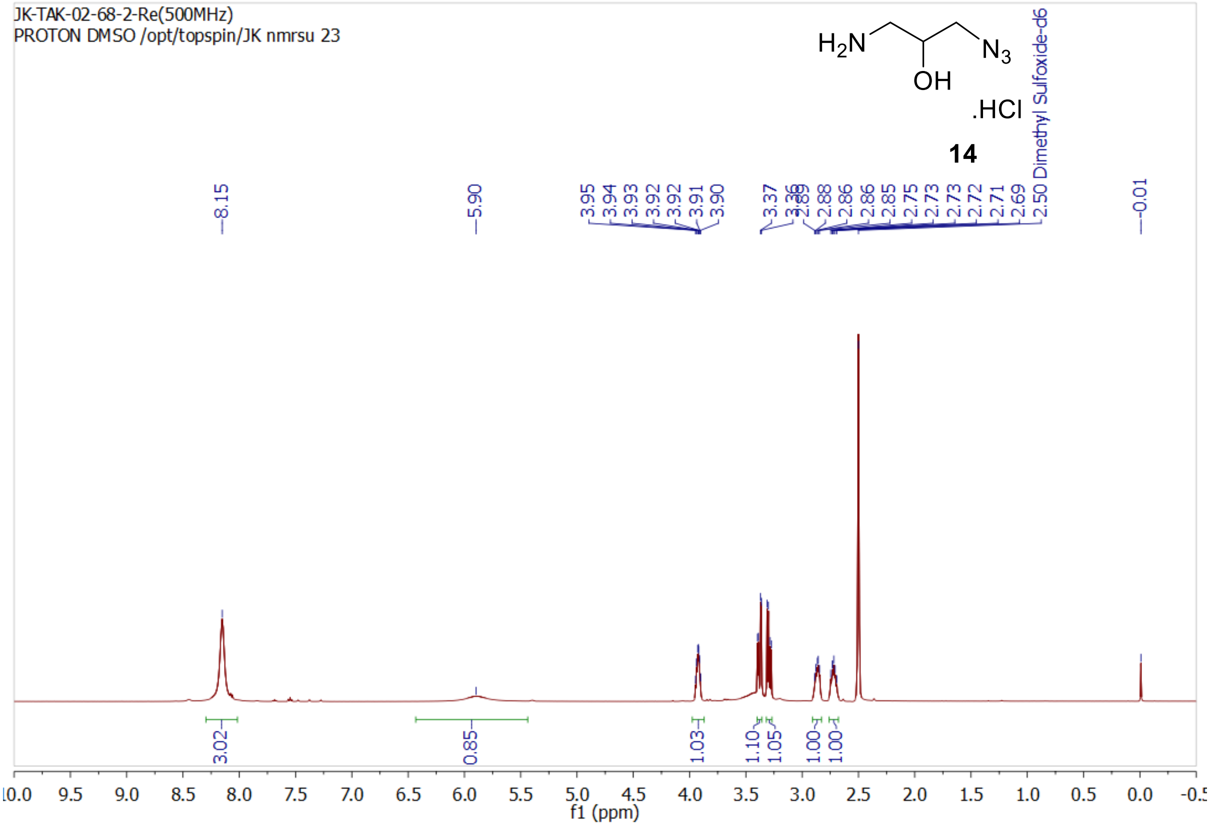
JK-TAK-02-68-2-Re(500MHz)
PROTON DMSO /opt/topspin/JK nmrsu 23
$H_2N$
OH
$N_3$
.HCl
14
2.50 Dimethyl Sulfoxide-d6
8.15
5.90
3.95
3.94
3.93
3.92
3.92
3.91
3.90
3.37
3.36
3.29
2.88
2.86
2.85
2.75
2.73
2.73
2.72
2.71
2.69
-0.01
3.02
0.85
1.03
1.10
1.05
1.00
1.00
f1 (ppm)

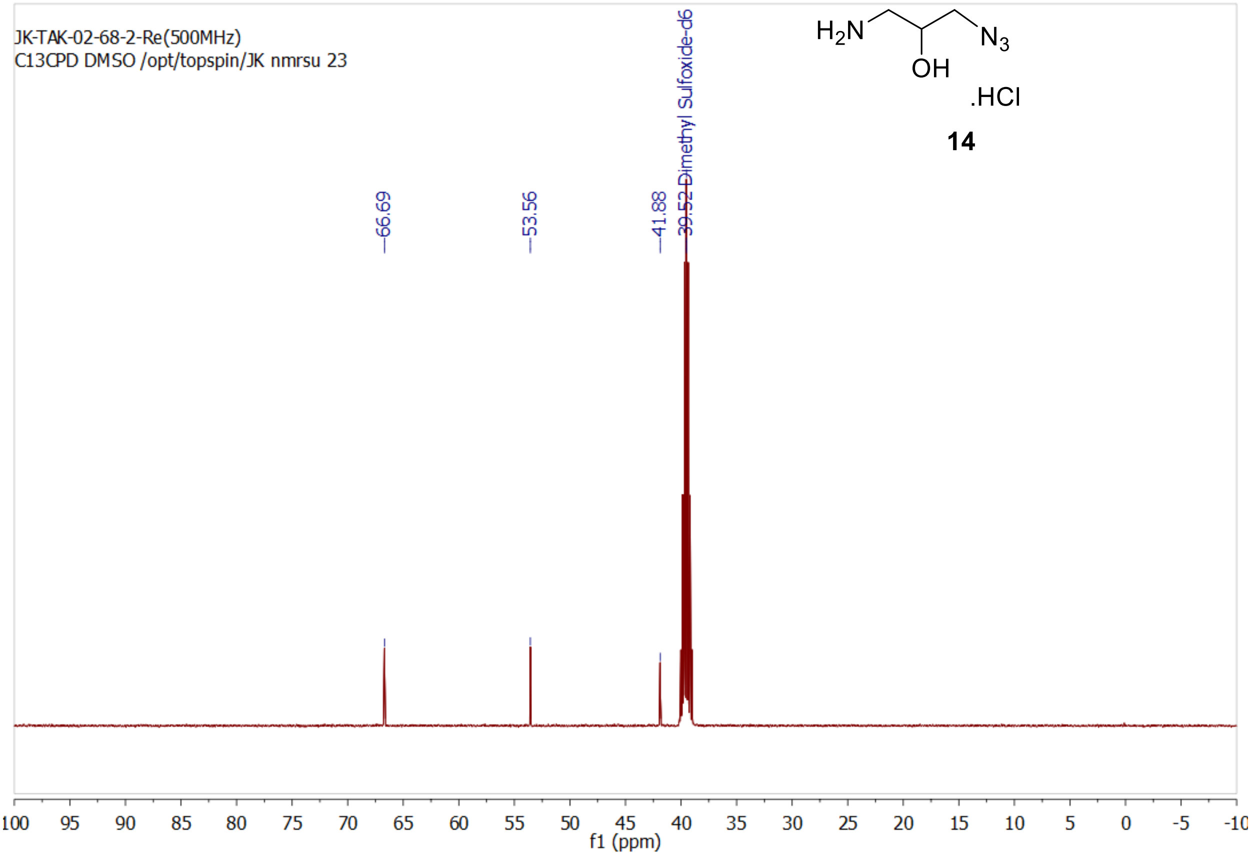
JK-TAK-02-68-2-Re(500MHz)
C13CPD DMSO /opt/topspin/JK nmrsu 23
$H_2N$
OH
$N_3$
.HCl
14
39.52 Dimethyl Sulfoxide-d6
66.69
53.56
41.88
f1 (ppm)

000

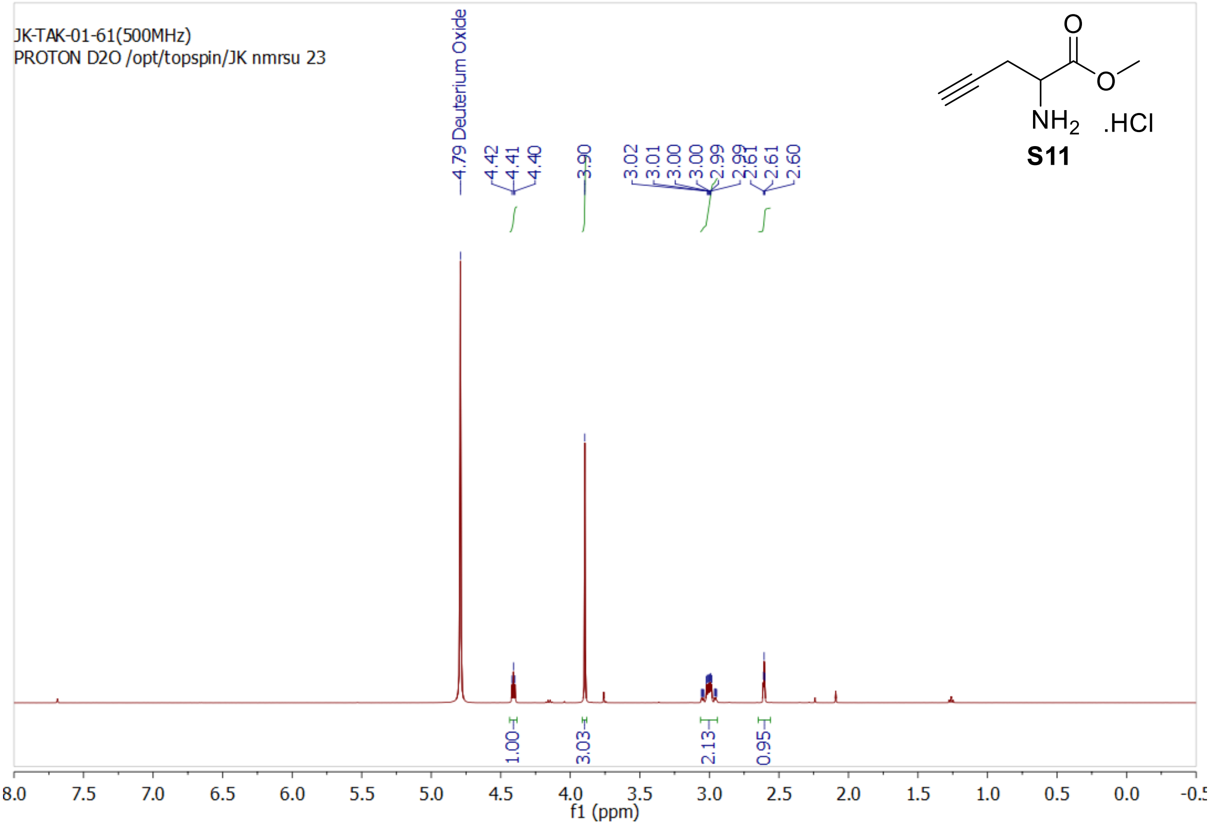
JK-TAK-01-61(500MHz)
PROTON D2O /opt/topspin/JK nmrsu 23
4.79 Deuterium Oxide
4.42
4.41
4.40
3.90
3.02
3.01
3.00
3.00
2.99
2.99
2.61
2.61
2.60
1.00
3.03
2.13
0.95
NH2
.HCl
S11
f1 (ppm)

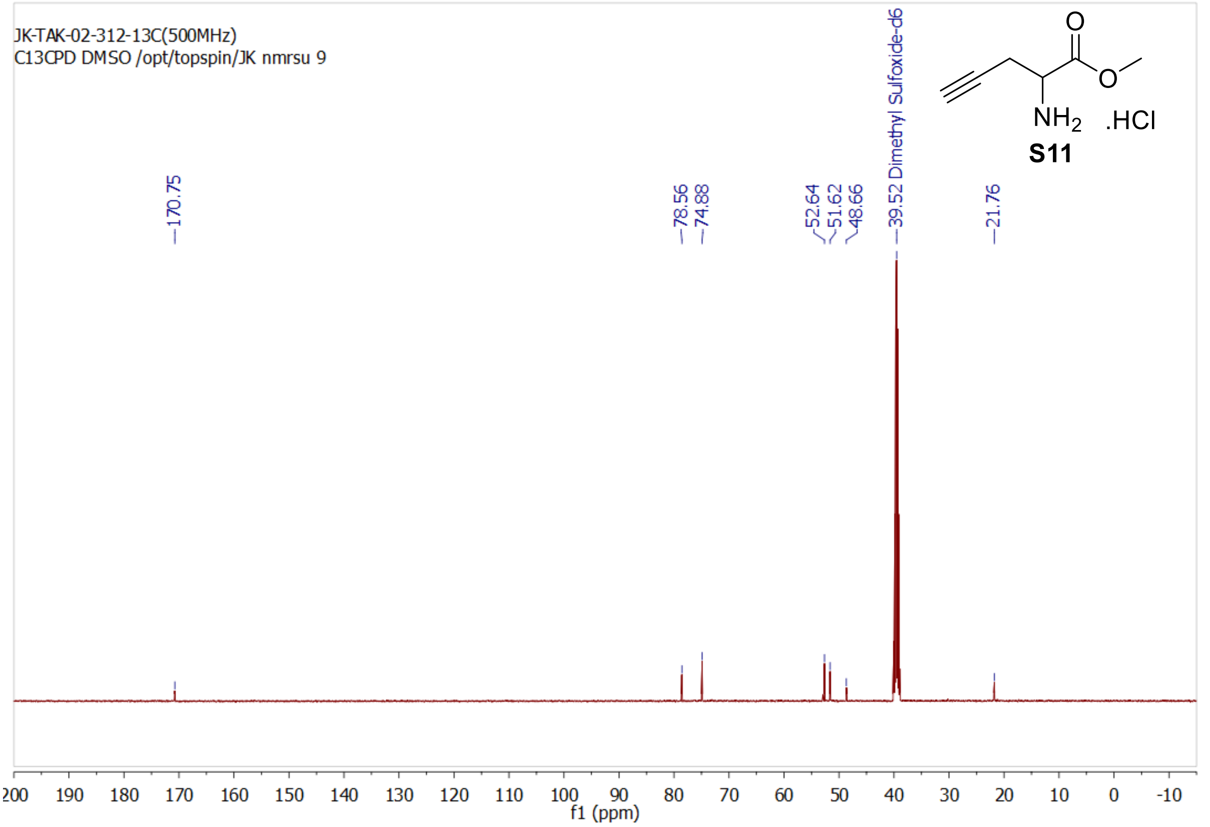
JK-TAK-02-312-13C(500MHz)
C13CPD DMSO /opt/topspin/JK nmrsu 9
170.75
78.56
74.88
52.64
51.62
48.66
39.52 Dimethyl Sulfoxide-d6
21.76
NH2
.HCl
S11
f1 (ppm)

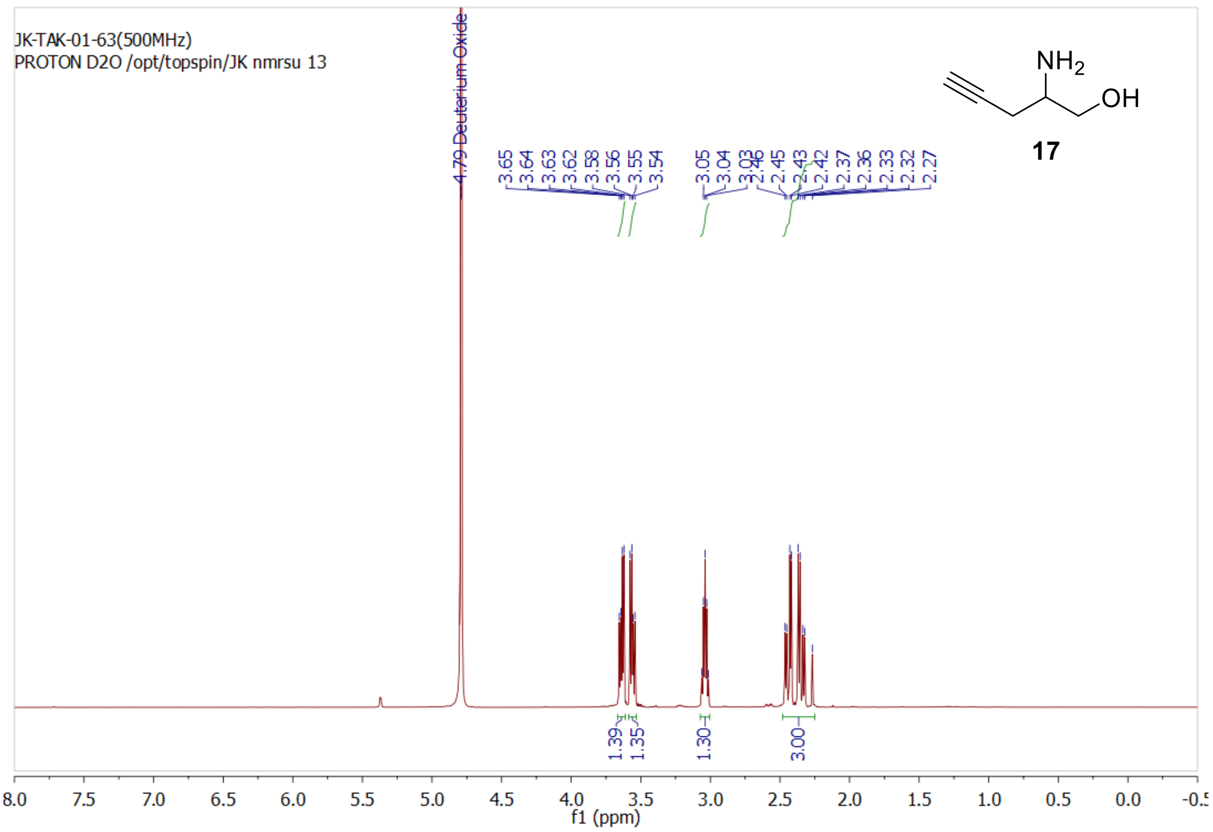
JK-TAK-01-63(500MHz)
PROTON D2O /opt/topspin/JK nmrsu 13
4.79 Deuterium Oxide
3.65
3.64
3.63
3.62
3.58
3.56
3.55
3.54
3.05
3.04
3.03
2.46
2.45
2.43
2.42
2.37
2.36
2.33
2.32
2.27
NH2
OH
17
1.39
1.35
1.30
3.00
8.0
7.5
7.0
6.5
6.0
5.5
5.0
4.5
4.0
3.5
3.0
2.5
2.0
1.5
1.0
0.5
0.0
f1 (ppm)

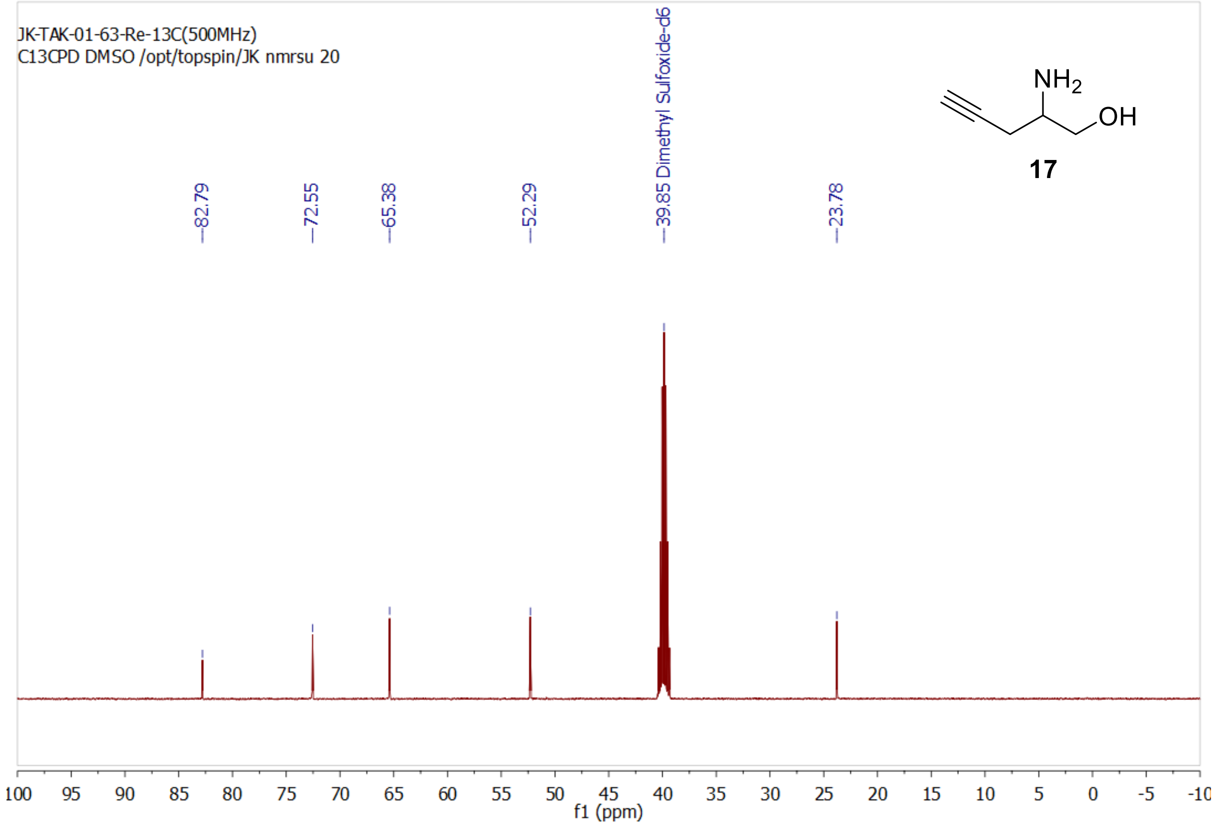
JK-TAK-01-63-Re-13C(500MHz)
C13CPD DMSO /opt/topspin/JK nmrsu 20
82.79
72.55
65.38
52.29
39.85 Dimethyl Sulfoxide-d6
23.78
NH2
OH
17
100
95
90
85
80
75
70
65
60
55
50
45
40
35
30
25
20
15
10
5
0
-5
f1 (ppm)

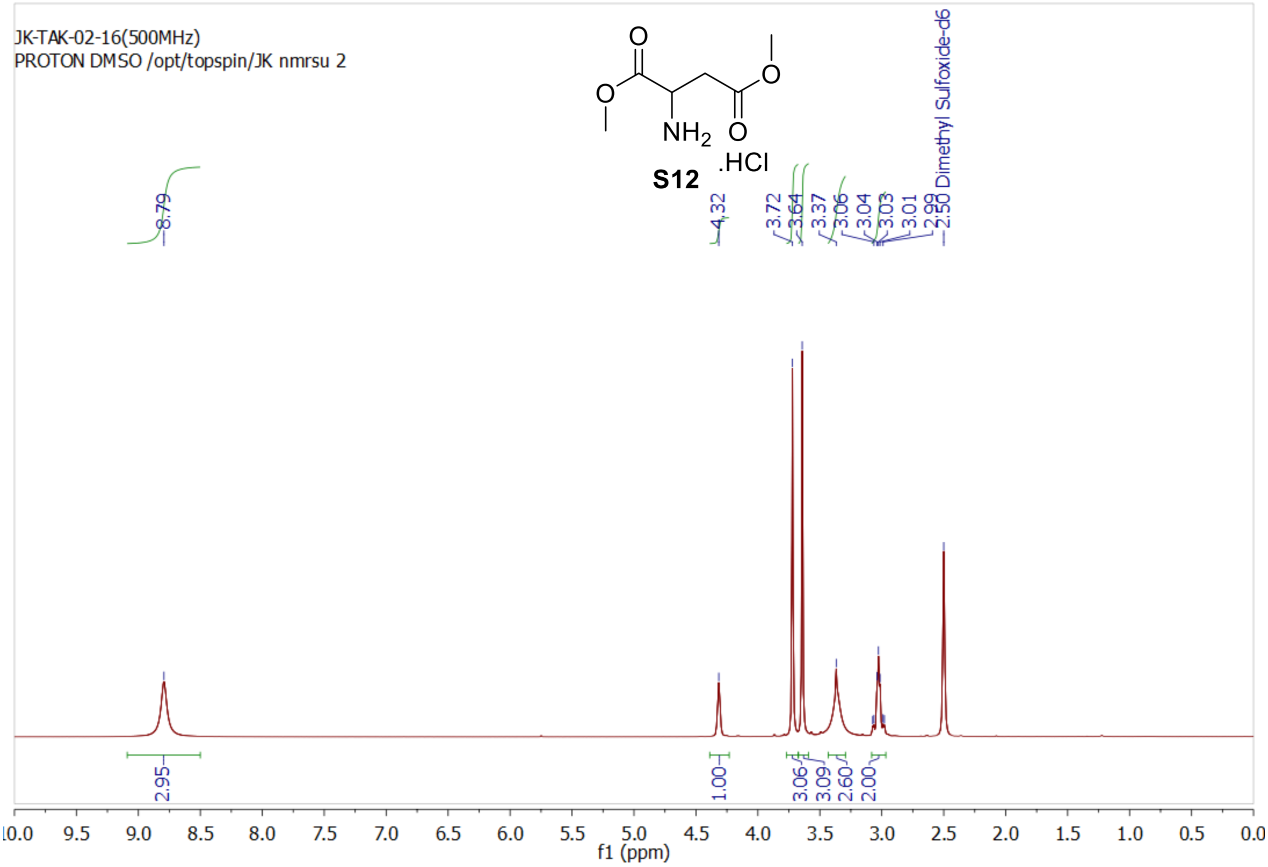
JK-TAK-02-16(500MHz)
PROTON DMSO /opt/topspin/JK nmrsu 2
NH$_2$
.HCl
S12
8.79
4.32
3.72
3.64
3.37
3.06
3.04
3.03
3.01
2.99
2.50 Dimethyl Sulfoxide-d6
2.95
1.00
3.06
3.09
2.60
2.00
f1 (ppm)

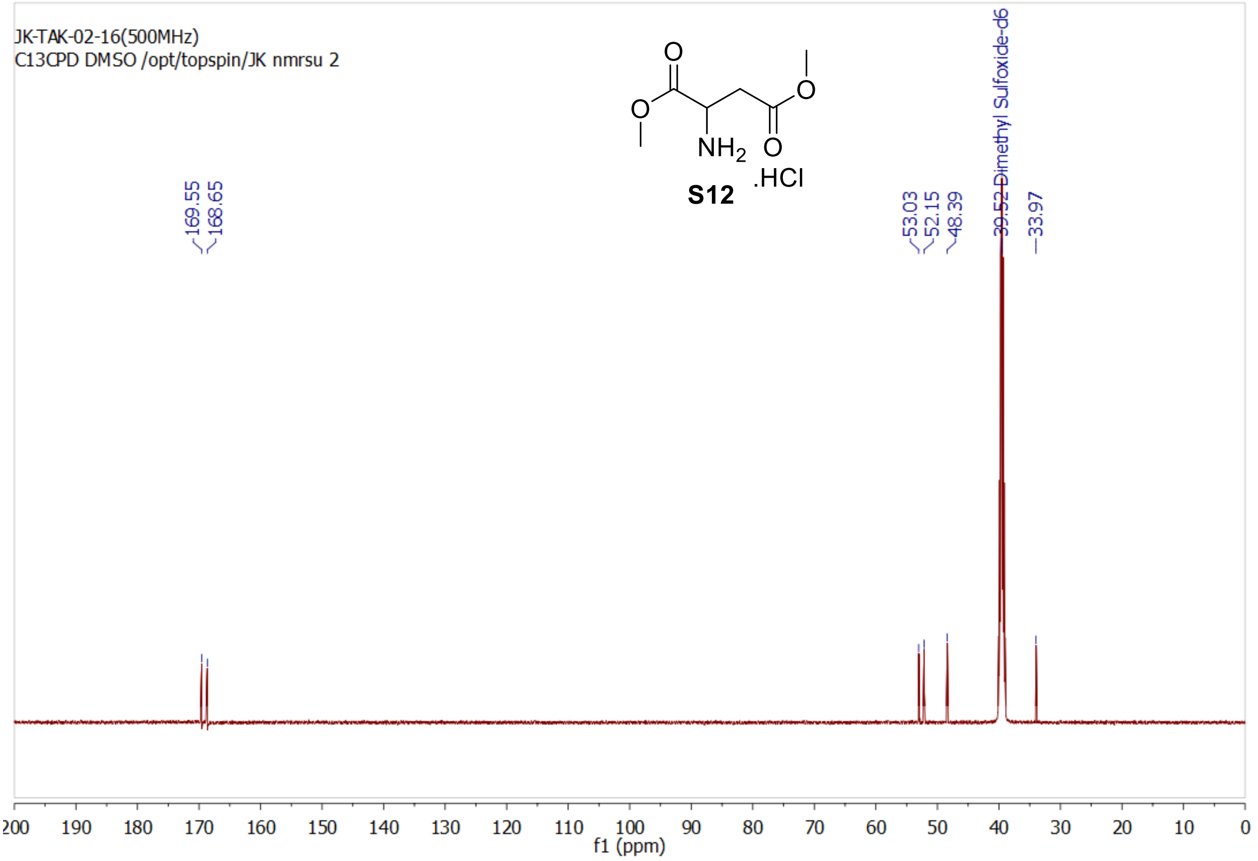
JK-TAK-02-16(500MHz)
C13CPD DMSO /opt/topspin/JK nmrsu 2
NH$_2$
.HCl
S12
169.55
168.65
53.03
52.15
48.39
39.52 Dimethyl Sulfoxide-d6
33.97
f1 (ppm)

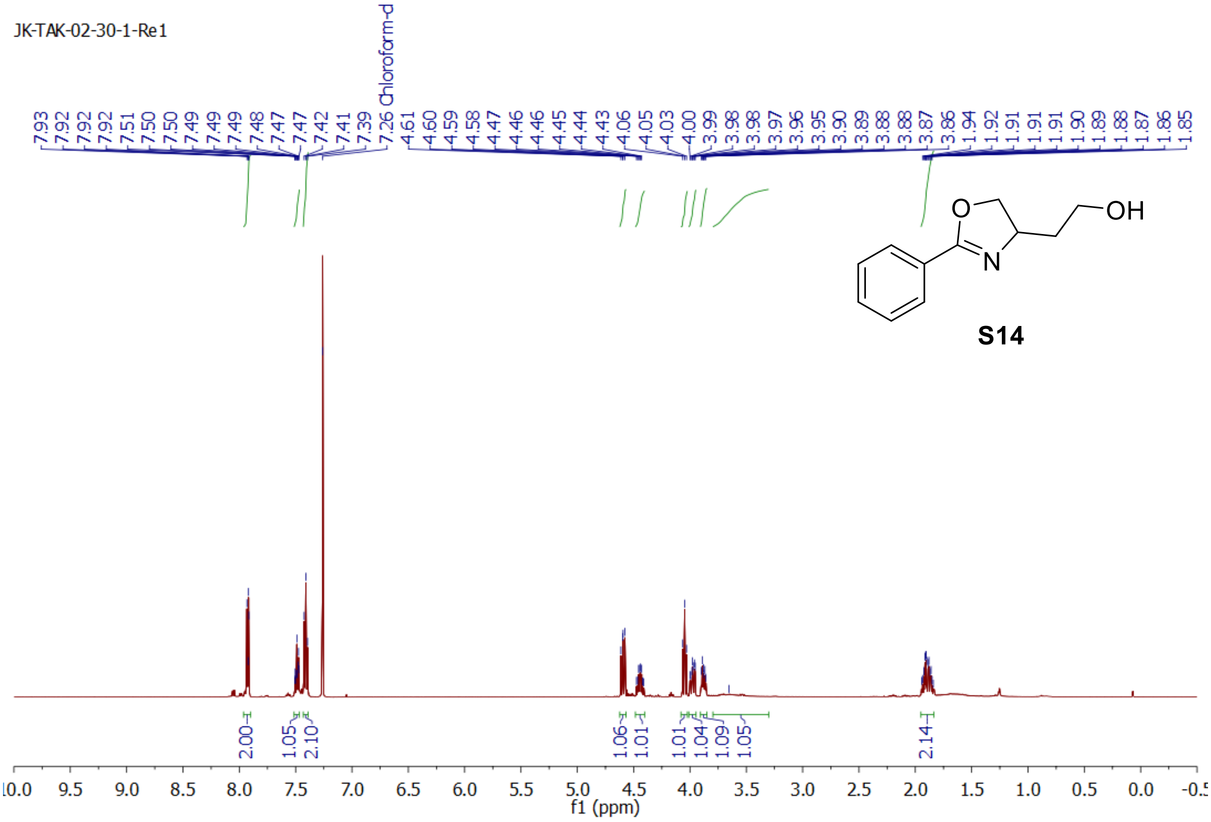
JK-TAK-02-30-1-Re1
7.93 7.92 7.92 7.92 7.51 7.50 7.50 7.49 7.49 7.49 7.48 7.47 7.47 7.42 7.41 7.39 7.26 Chloroform-d 4.61 4.60 4.59 4.58 4.47 4.46 4.46 4.45 4.44 4.43 4.06 4.05 4.03 4.00 3.99 3.98 3.98 3.97 3.96 3.95 3.90 3.89 3.88 3.88 3.87 3.86 1.94 1.92 1.91 1.91 1.90 1.89 1.88 1.87 1.86 1.85
2.00 1.05 2.10 1.06 1.01 1.01 1.04 1.09 1.05 2.14
f1 (ppm)
O
N
OH
S14

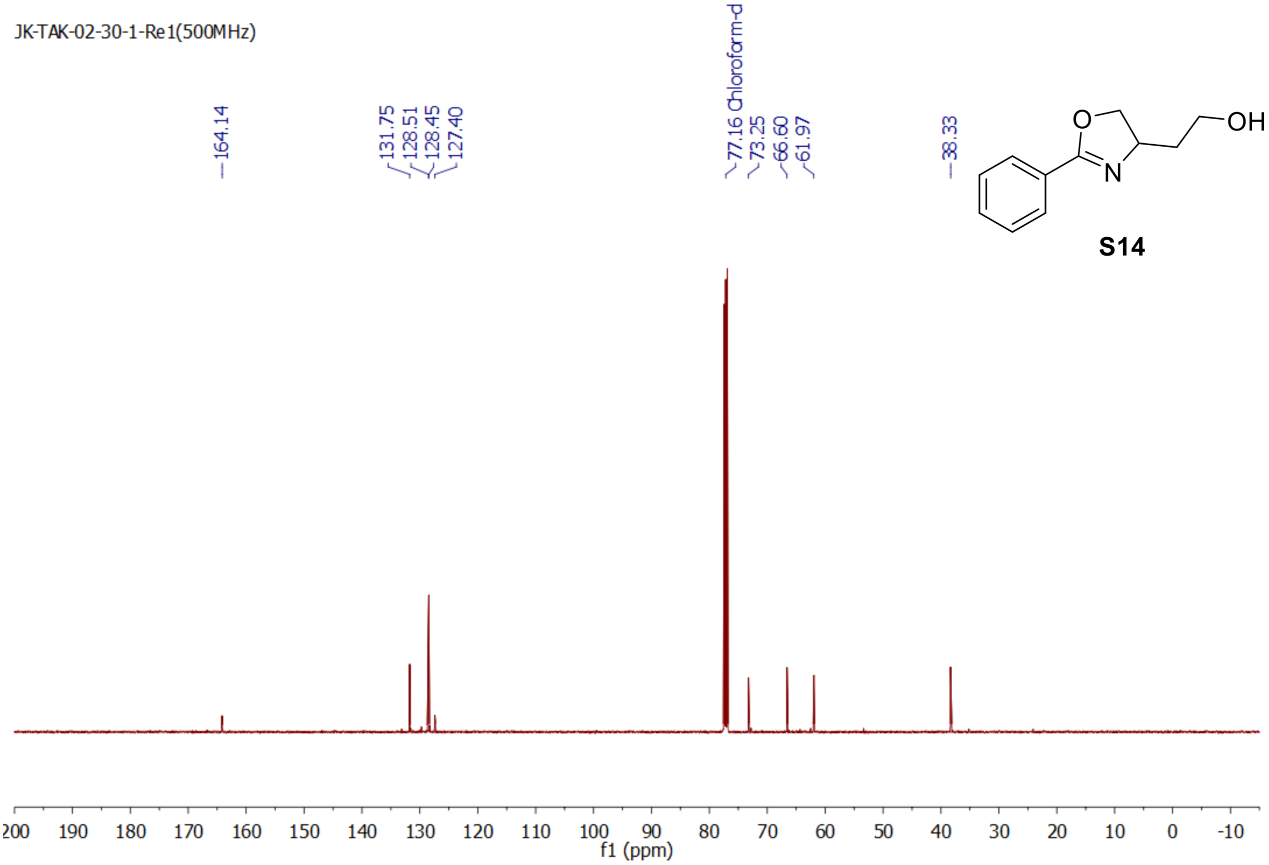
JK-TAK-02-30-1-Re1(500MHz)
164.14
131.75 128.51 128.45 127.40
77.16 Chloroform-d 73.25 66.60 61.97
38.33
f1 (ppm)
O
N
OH
S14

JK-TAK-02-59(500MHz)
PROTON CDCl3 /opt/topspin/JK nmrsu 4

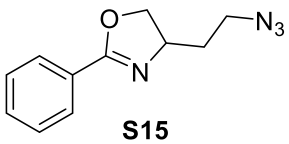

**S15**

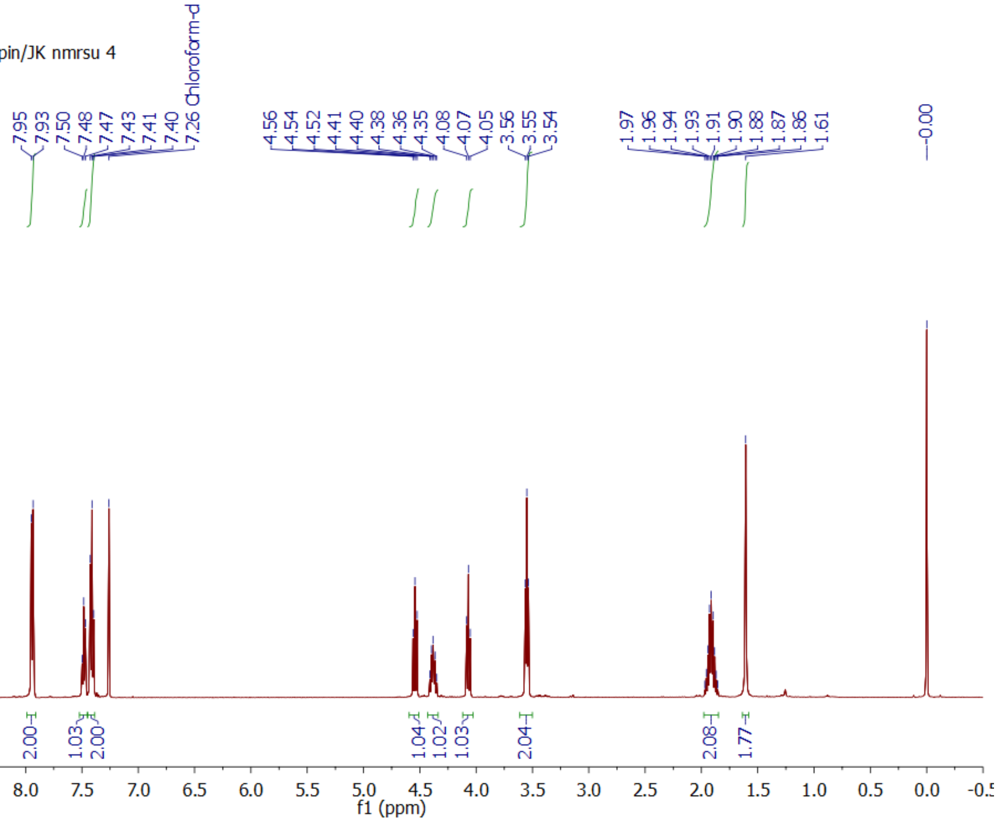

JK-TAK-02-59-13C(500MHz)
C13CPD CDCl3 /opt/topspin/JK nmrsu 2

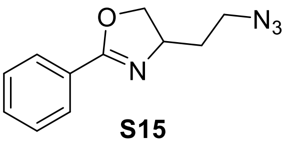

**S15**

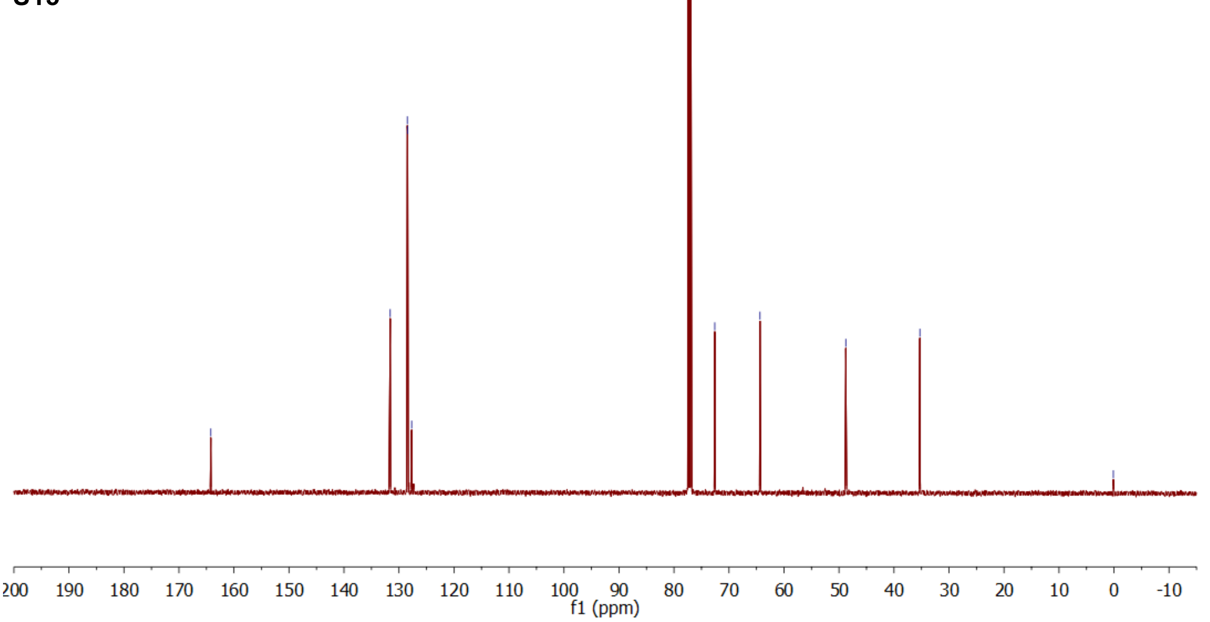

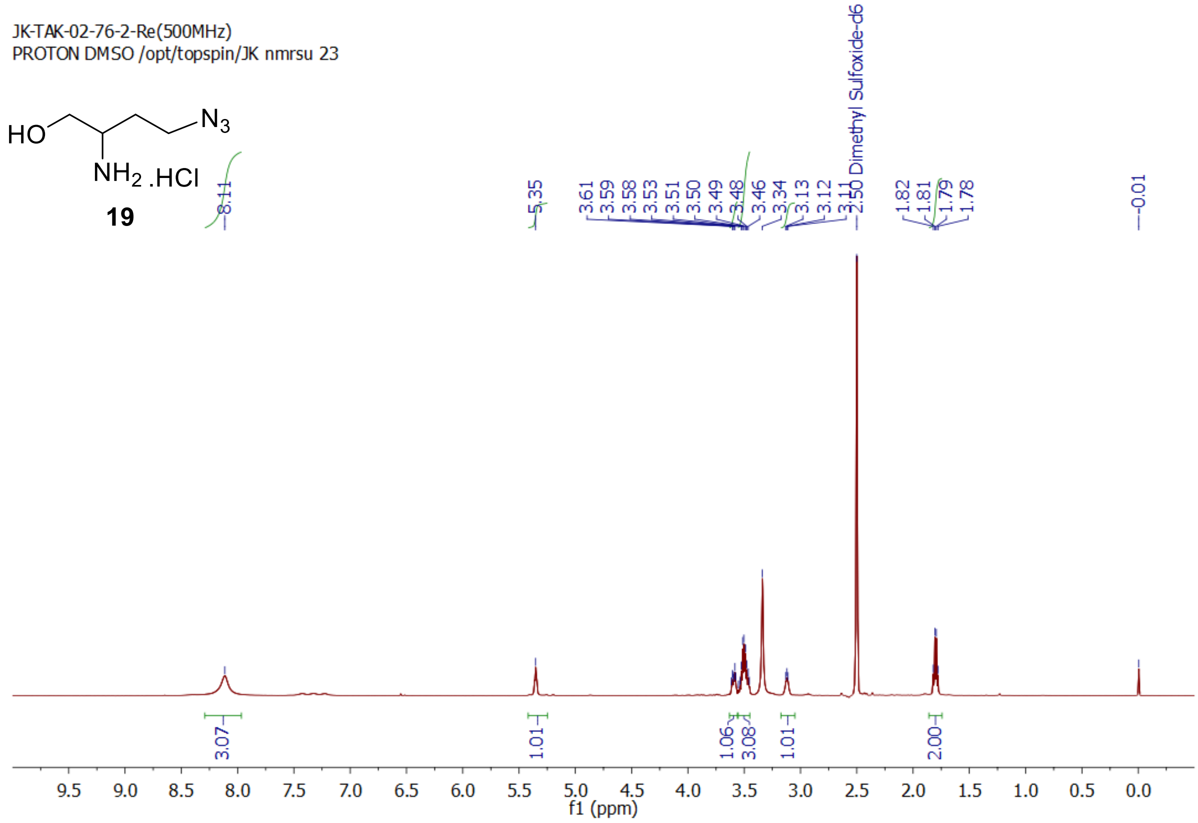
JK-TAK-02-76-2-Re(500MHz)
PROTON DMSO /opt/topspin/JK nmrsu 23
HO
N3
NH2 .HCl
19
8.11
5.35
3.61 3.59 3.58 3.53 3.51 3.50 3.49 3.48 3.46 3.34 3.13 3.12 3.11
2.50 Dimethyl Sulfoxide-d6
1.82 1.81 1.79 1.78
-0.01
3.07
1.01
1.06
3.08
1.01
2.00
f1 (ppm)

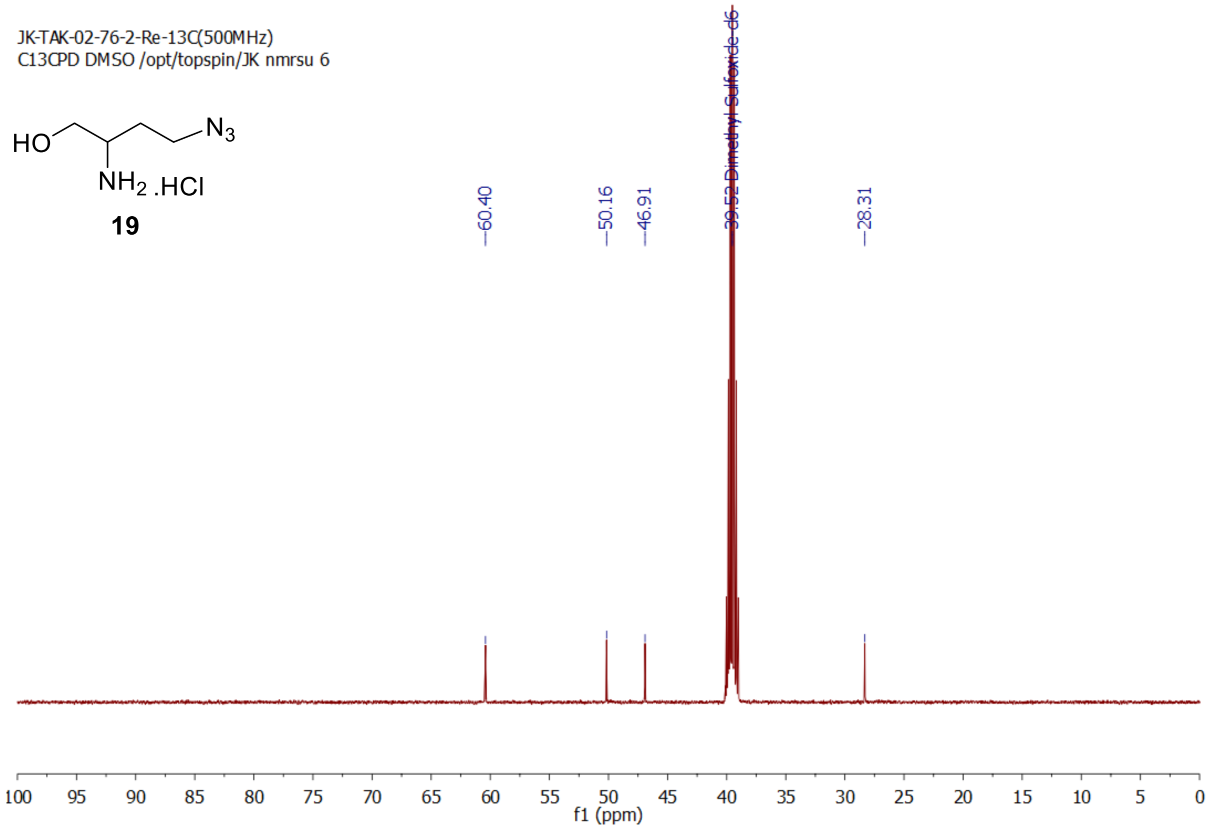
JK-TAK-02-76-2-Re-13C(500MHz)
C13CPD DMSO /opt/topspin/JK nmrsu 6
HO
N3
NH2 .HCl
19
60.40
50.16
46.91
Dimethyl Sulfoxide-d6
28.31
f1 (ppm)
